# Winter Foraging Selectivity of Blackbuck in Semi-Arid Agricultural Landscapes

**DOI:** 10.1101/2025.07.27.666993

**Authors:** Mohd Yunus, Orus Ilyas

## Abstract

The blackbuck (*Antilope cervicapra*), a native grazer of the Indian subcontinent, is increasingly confined to semi-arid, human-modified landscapes. Understanding its seasonal foraging patterns is essential for conservation planning in such habitats. This study examined winter dietary preferences of blackbuck across three agro-pastoral sites—Palla Salu, Rathgaon, and Sikandra Rao—in Aligarh district, Uttar Pradesh, India. Vegetation sampling recorded 4,125 plant individuals, with ***Prosopis juliflora*** and ***Cynodon dactylon*** dominating the landscape. However, micro-histological analysis of 45 fecal pellet samples identified **18 distinct plant species “**comprising **a total of 860** identifiable fragments”. Of these, ***C. dactylon, Poa annua, Rumex pulcher*, and *Atriplex patula*** were significantly preferred. Despite high field abundance, *P. juliflora* was consistently avoided, likely due to structural or chemical deterrents. Interestingly, rare species such as ***Chenopodium vulvaria*** were overrepresented in the diet, suggesting nutrient-specific selectivity. Bonferroni-adjusted confidence intervals confirmed significant foraging preferences across all sites. These findings highlight the blackbuck’s adaptive and selective winter feeding strategy and the crucial role of native grasses and forbs during resource-scarce periods. We recommend the control of invasive species and restoration of native forage to enhance habitat quality and support long-term blackbuck conservation in fragmented, human-dominated ecosystems.

## INTRODUCTION

The blackbuck (*Antilope cervicapra*), a native grazer of the Indian subcontinent, is emblematic of open grassland ecosystems. Once widespread, its populations have declined sharply due to extensive habitat loss, agricultural expansion, and human encroachment (Ranjitsinh, 1989; Jhala, 1997). As a primarily grazing species, blackbuck exhibit strong seasonal dietary shifts driven by forage availability and landscape structure (Prasad, 1981; Ghosh et al., 1984). During the monsoon, they congregate in grasslands and consume tender shoots, occasionally browsing on newly sprouted foliage. However, in winter, when grass protein content declines, they increasingly shift to crop fields, feeding on on wheat, barley, gram, and legumes, particularly in areas like Velavadar (Gujarat) and Mudmal (Tamil Nadu) (Ranjitsinh, 1989; Jhala, 1993a). They are also known to consume seed pods of native *Prosopis cineraria* and exotic *P. juliflora*, which may comprise a significant portion of the winter diet (Jhala, 1991). In late winter and early summer, when crops are harvested and grasses are depleted, blackbuck utilize moist patches and browse species like *Acacia* and *Phoenix*. Despite lower abundance in the diet, browse provides essential protein and energy during resource-limited periods (Goyal et al., 1988; Prasad, 1981).

Habitat use in blackbuck is influenced not only by forage availability but also by anti-predator strategies. The species avoids tall vegetation and dense shrub lands due to obstructed visibility and heightened predation risk, preferring open short-grass habitats that allow early predator detection and rapid escape (Isvaran, 2007; Mungall, 1978). This behavioral ecology, while adaptive, makes blackbuck more vulnerable in fragmented agro-pastoral landscapes where crop fields dominate and suitable forage patches are patchy and seasonal. In Aligarh district, Uttar Pradesh—a semi-arid agricultural matrix—blackbuck persist in scattered habitats embedded within farmlands, village edges, and fallow grasslands. Although previous studies in other regions have documented their seasonal diet composition (Jhala, 1991; Mishra et al., 2020), data specific to winter foraging ecology in northern India remain limited. Understanding blackbuck’s winter diet and selectivity is essential not only for species conservation but also for mitigating crop damage and human-wildlife conflict in this heavily cultivated region. This study investigates the winter foraging behavior of blackbuck across three agro-pastoral landscapes in Aligarh using vegetation surveys and fecal micro-histology to assess diet composition and selectivity. The findings will inform conservation strategies focused on habitat restoration, invasive species management, and co-existence in agricultural landscapes.

## STUDY AREA

This study was conducted in Aligarh district, located western Prades in Uttar, India, on the southern margin Upper of the Ganga– Yamuna Doab. The region lies approximately 100 km southeast of Delhi and about 40 km southwest of the Ganges River, covering an area of 5,019 km^2^ at an elevation of around 178 meters above sea level. The climate is classified as hot semi-arid (Köppen BSh), characterized by scorching summers (April–June), monsoon rainfall (∼800–1,000 mm annually) from late June to September, and mild winters (2°C–22°C). Soils are primarily loamy with patches of sandy deposits, shaped by aeolian activity from the Thar Desert. These conditions support a mosaic of agricultural fields, fallow lands, and remnant vegetation patches—ideal for winter foraging by large herbivores like blackbuck (*Antilope cervicapra*).

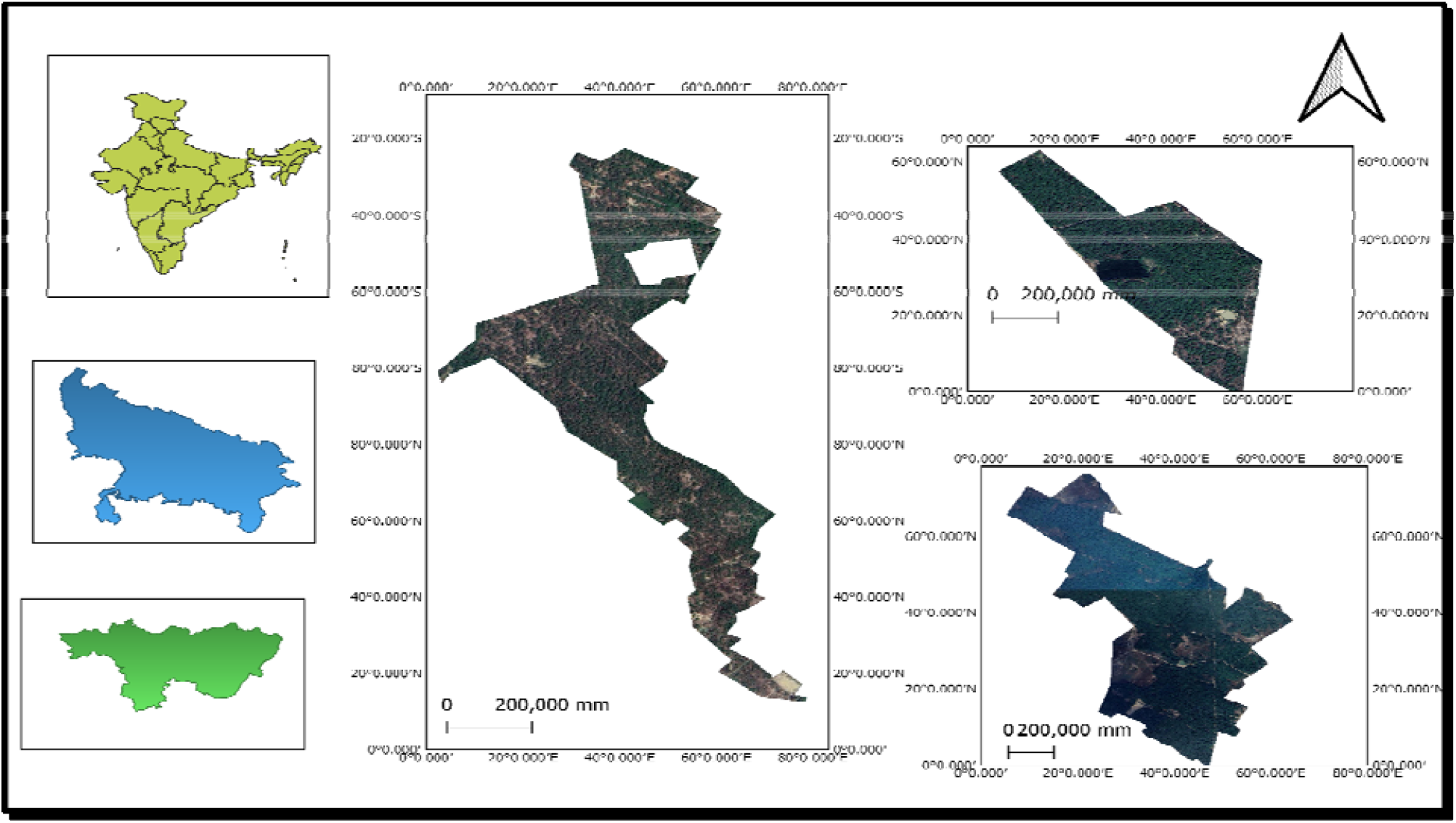

Fieldwork was conducted at three key winter habitats within Aligarh: **Palla Salu** (0.3 km^2^), **Rathgaon** (0.4 km^2^), and **Sikandra Rao** (0.9 km^2^). These sites were selected based on prior observations of consistent blackbuck activity during the winter season. All three sites are embedded in human-dominated landscapes composed of farmland, scrub vegetation, and seasonal grasses. Despite their small size, they serve as important winter foraging grounds, offering a variety of native and invasive plant species. Satellite imagery and field verification confirmed their extent and accessibility, making them suitable for ecological assessment involving quadrat-based vegetation surveys and fecal pellet sampling for diet analysis.

## METHODOLOGY

To comprehensively understand the winter foraging behavior of blackbuck (*Antilope cervicapra*) in semi-arid agro-pastoral systems, an extensive field-based investigation was undertaken in **Aligarh district**, located in western Uttar Pradesh, India. The research focused on three key sites Palla Salu (0.33lJkm^2^), Sikandra Rao (0.90lJkm^2^), and athgaon (0.40lJkm^**2**^) which were purposively selected based on long-term observational data and published literature indicating the consistent seasonal presence and grazing activity of blackbuck populations in these areas during winter (Jhala, 1991). These sites represent a typical gradient of land-use patterns in the region, encompassing natural grassland, fallow agricultural fields, and semi-managed scrublands.

To capture temporal variability in vegetation structure and availability during the dry season, monthly vegetation surveys were conducted over a span of three months—January, February, and March 2025. Surveys were deliberately scheduled between the 15th and 17th of each month to ensure temporal consistency in sampling and to reduce phenological variation that could confound ecological interpretation. This time frame also coincided with the period when resource scarcity intensifies, compelling herbivores such as blackbuck to make strategic foraging decisions.Vegetation sampling was conducted using a stratified line transect method, which is widely recognized for its ability to provide representative coverage across habitat types. At each of the three study sites, three 200-meter transects were systematically laid out. Along each transect, sampling quadrats were positioned at 20-meter intervals, with a 10-meter offset perpendicular to the transect line. To avoid spatial bias and ensure comprehensive spatial representation, the quadrats alternated between the left and right sides of the transect path (Mishra, 1968).The dimensions of the quadrats were standardized according to vegetation strata following established ecological conventions (Kershaw, 1973). Specifically, 1 × 1 meter quadrats were used to sample herbaceous species and graminoids, 5 × 5 meter quadrats for woody shrubs, and 10 × 10 meter quadrats for tree species. Within each quadrat, all plant species were identified, and their individual counts, phenological stages (e.g., vegetative, flowering, fruiting). These data provided a detailed picture of available forage resources in terms of both quantity and quality.To assist with real-time plant identification in the field, the PlantNet mobile application was employed. This tool uses machine learning-based image recognition to enhance accuracy in species identification, particularly in biodiverse and poorly documented semi-arid ecosystems (PlantNet, n.d.). Following identification, phytosociological parameters—including frequency, density, and relative abundance—were computed for each species using formulas described by Mishra (1968). These quantitative measures were later used to estimate plant availability in the environment and to compare with dietary preferences obtained from fecal analysis.

In parallel with vegetation sampling, reference leaf samples were opportunistically collected (encounter-based sampling) from the three focal sites—Palla Salu, Sikandra Rao, and Rathgaon—to build a comprehensive reference library of potential forage species. These samples were gathered across various habitat types, including natural grasslands, scrublands, field margins, and anthropogenically influenced edges, ensuring broad ecological coverage and capturing spatial heterogeneity in plant availability (Sankaran et al., 2005; Augustine & McNaughton, 1998; Rahmani, 2001). This method allowed inclusion of rarely encountered or seasonally limited plant species that might not appear in quadrat surveys but could be consumed by blackbuck.

Each collected plant sample was carefully tagged with site location, date, and habitat type, and stored in labeled, breathable paper envelopes to minimize decomposition. The plant materials were later used to prepare reference microscope slides, which are essential for accurate species identification during fecal microhistological analysis. Slide preparation followed standard microhistological protocols (Sparks & Malechek, 1968; Holechek et al., 1982), involving leaf fragment treatment to preserve distinct cuticular features.To assess the actual dietary composition of blackbuck, fecal pellet sampling was conducted alongside habitat surveys. At each site, three 500-meter transects were established across representative grazing habitats, including open fields, scrub zones, and ecotonal areas. Along these transects, blackbuck fecal pellets were located and collected opportunistically, ensuring samples reflected the animal’s natural foraging patterns with minimal human disturbance (Buckland et al., 2001; Putman, 1984). A total of 45 fecal samples—15 from each site—were collected, representing adequate sample size thresholds for dietary studies in medium-sized ungulates (Gotelli & Ellison, 2013; Levy & Lemeshow, 2013).Upon collection, fecal samples were air-dried at room temperature, then ground into fine powder using a mortar and pestle. The powdered material was passed through ASTM No. 30 and 60 mesh sieves to isolate fragments of optimal size for microscopic analysis. The sieved material was then quartered to ensure randomness and uniformity in subsampling. Samples were treated with a 1:3 solution of nitric acid and distilled water to clear organic debris, then dehydrated through a graded ethanol series and mounted permanently in Canada balsam on clean glass slides (Baumgardt et al., 1964; Storr, 1961).Each prepared slide was examined under a compound microscope, with five randomly selected fields of view per slide analyzed for consistent representation. Plant fragments were identified based on distinctive cuticular features such as epidermal cell patterns, stomatal structures, and trichomes, and were matched with the prepared reference slides to ensure accurate species-level identification (Stewart, 1967; Sparks & Malechek, 2004). This integrative and comparative microhistological technique provided a robust dataset to determine the composition and diversity of the blackbuck’s winter diet across habitat types.

To assess blackbuck dietary selectivity, proportional utilization of each plant species identified from fecal samples was statistically compared with its proportional availability in the environment. Specifically, the frequency of occurrence for each plant taxon in the fecal material was calculated and denoted as P□□, while its corresponding availability in the field was represented as PDD. These two proportions were then compared using Bonferroni-adjusted 95% confidence intervals, a method widely employed in herbivore dietary studies (Neu et al., 1974; Byers et al., 1984).

Plant species were categorized into three groups: preferred, avoided, or neutrally used. A species was considered preferred if P□□ fell below the lower bound of the confidence interval for PDD; avoided if P□□: exceeded the upper bound; and neutrally used if PDD fell within the interval.

The confidence interval for proportional utilization was calculated using the formula: P□□ ± Z (α/2k) × √[P□□ (1 − P□□) / n]

Where:

- P□□ is the proportion of fragments of a particular species in the fecal sample
- Z(α/2k) is the Bonferroni-adjusted Z-value based on the number of comparisons (k)
- n is the total number of fragments identified across all samples
- k is the number of plant species included in the analysis
- α is the significance level, typically set at 0.05

In this context, b refers to the number of fragments of a particular species in the fecal dataset, and ΣB is the total number of plant fragments observed. Likewise, n represents the count of individuals for that species in the field vegetation data, and ΣN is the total number of individual plants recorded. This statistical approach allowed for an objective, quantitative classification of plant species based on their selection or avoidance, offering key insight into the winter foraging preferences of blackbuck (Zar, 1999; Fowler & Cohen, 1986).

## RESULT

### 1 Important Value Index

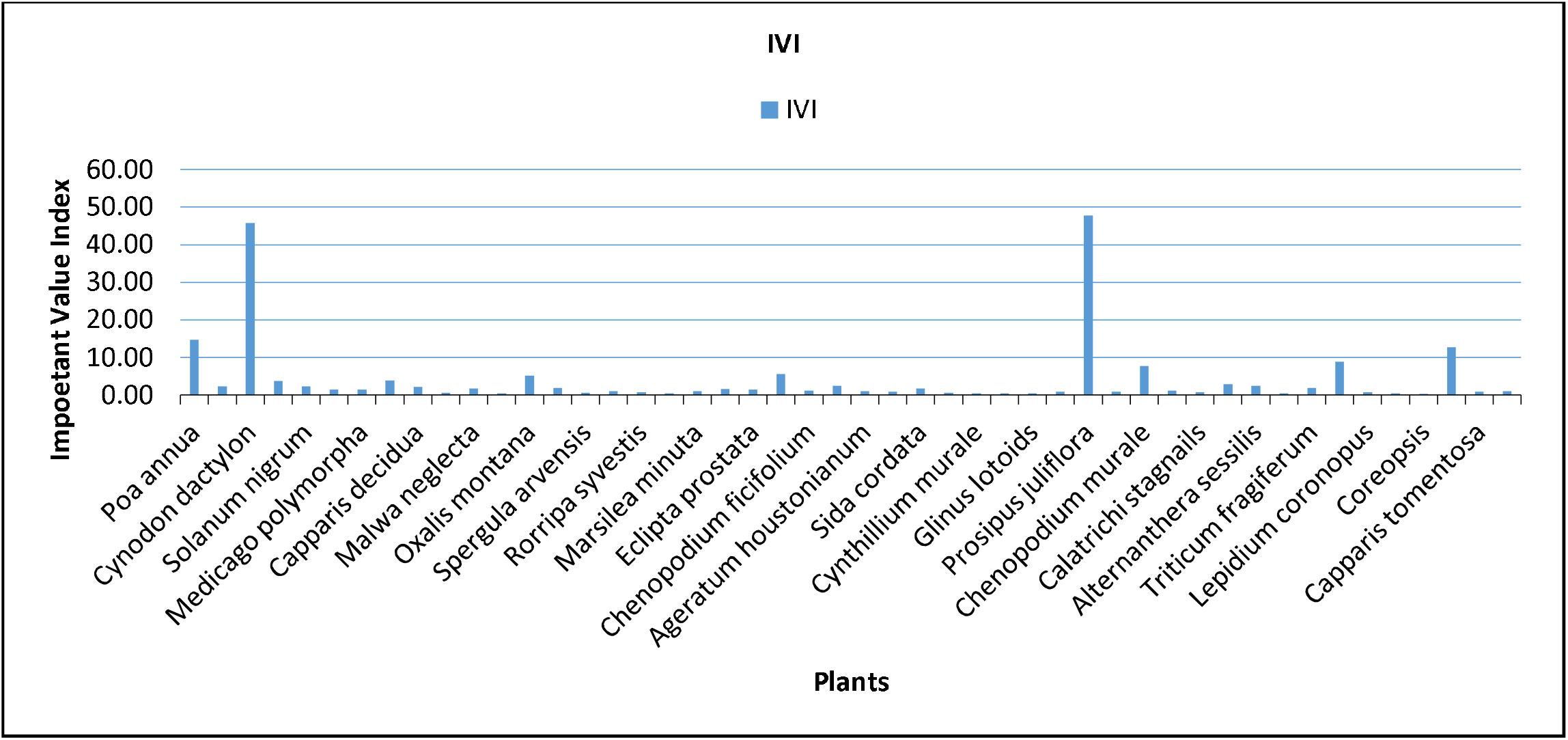

A total of 4,125 individual plants belonging to a diverse assemblage of herbaceous and shrub species were recorded across the study sites of Palla Salu, Rathgaon, and Sikandra Rao. Among these, *Prosopis juliflora* (698 individuals), *Cynodon dactylon* (754 individuals), and *Poa annua* (314 individuals) emerged as the most abundant species within the sampled quadrats. These taxa demonstrated a high degree of ecological adaptability to semi-arid conditions and were consistently encountered across multiple sampling units, highlighting their dominance in the study landscape. In contrast, several species such as *Rorripa sylvestris* (2 individuals), *Atriplex patula, Datura innoxia*, and *Carduus tenuiflorus* (each represented by 3 individuals) were recorded infrequently. Their low occurrence may indicate either ecological rarity, narrow habitat specificity, or sensitivity to environmental fluctuations and anthropogenic disturbances.

The Importance Value Index (IVI) analysis further underscored the dominance of a few key species in structuring the vegetation community. *Prosopis juliflora* (IVI = 47.81), *Cynodon dactylon* (IVI = 45.84), and *Poa annua* (IVI = 14.62) ranked highest in IVI scores, affirming their prevalence and relative ecological importance. While *C. dactylon* is a well-documented forage grass commonly preferred by large herbivores, *P. juliflora* is an invasive alien species known for its aggressive expansion and ability to displace native flora. Several other taxa, including *Calamagrostis epigejos, Chenopodium vulvaria*, and *Chenopodium murale*, exhibited moderate IVI values, indicating a secondary role in the plant community. In contrast, species such as *Coreopsis, Rumex obtusifolius*, and *Cyanthillium cinereum* (formerly listed as *Cynthillium murale*) had very low IVI scores, suggesting a restricted or patchy distribution, likely influenced by site-specific microhabitat conditions or competitive exclusion by dominant species.

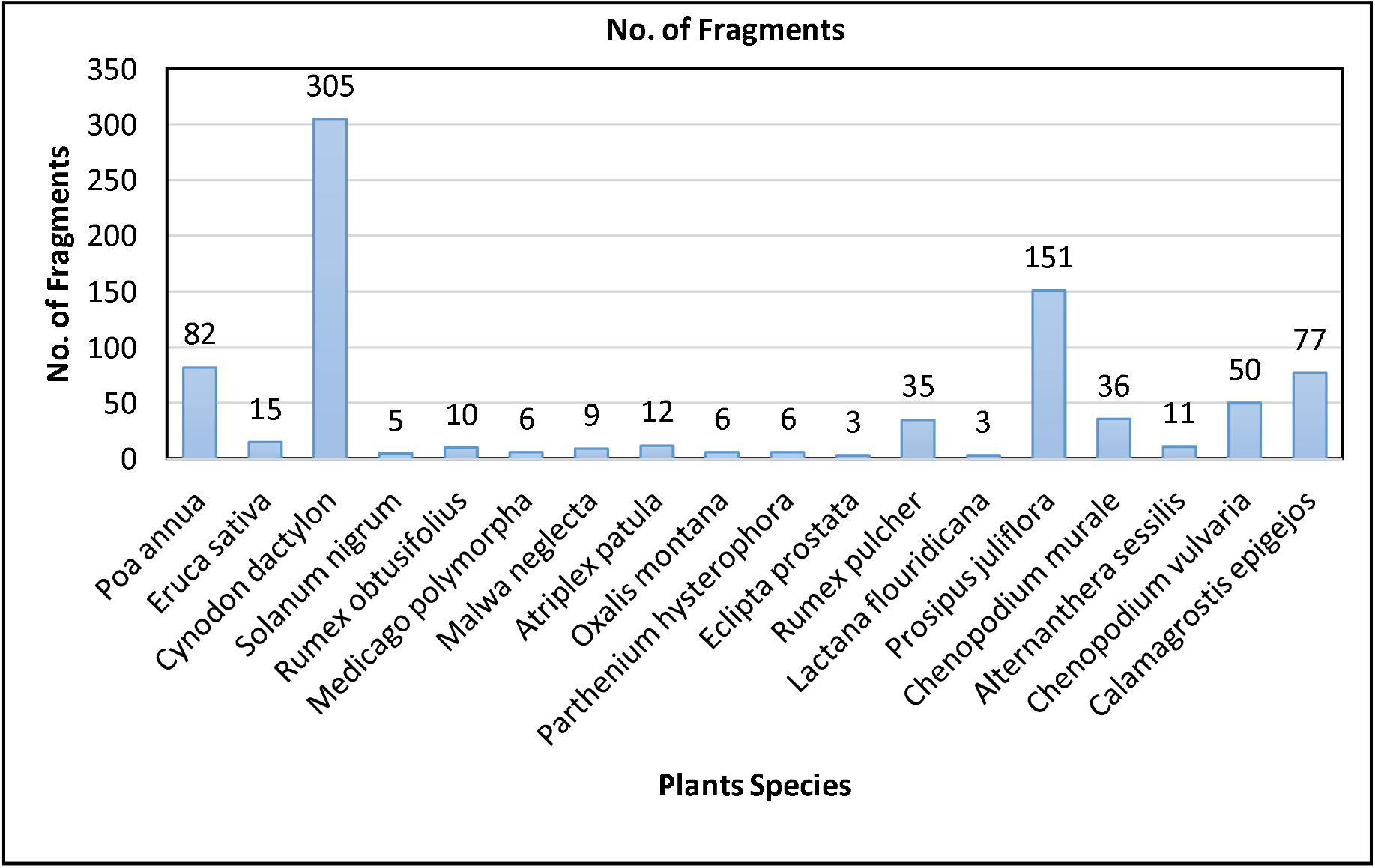

Overall, the vegetation structure across the study sites revealed a distinctly skewed composition, characterized by a few ecologically dominant and widely distributed species, alongside a larger cohort of localized, less abundant taxa. This pattern aligns with typical ecological profiles observed in intensively grazed or anthropogenically disturbed grassland systems, where grazing pressure and habitat fragmentation contribute to uneven species distribution and reduced community evenness.

Microhistological analysis of fecal pellets revealed a total of 860 plant fragments, representing 18 distinct species consumed by blackbuck across the study area. Application of Bonferroni-adjusted confidence intervals provided clear evidence of selective foraging behavior. Among the species identified, *Cynodon dactylon* was the most preferred forage, constituting 34.9% of the identified dietary fragments, a proportion significantly exceeding its availability in the environment (27.7%). Other species such as *Poa annua* (10.4% use vs. 7.5% availability), *Rumex pulcher* (4.0% use vs. 1.6% availability), and *Atriplex patula* also demonstrated significant positive selection, likely attributed to their high palatability, favorable digestibility, or superior nutritional content.

In contrast, *Prosopis juliflora*, despite its ecological dominance and high field availability (25.7%), was underrepresented in the fecal samples, accounting for only 17.3% of the dietary fragments. This pattern indicates active avoidance, potentially due to the species’ spiny morphology, lignified structure, or the presence of allelopathic compounds and chemical deterrents that reduce its desirability as a forage resource.

Other species that were noticeably underutilized included *Solanum nigrum, Lactuca floridana*, and *Parthenium hysterophorus*, which are known to contain toxic alkaloids or other secondary metabolites that discourage herbivory. On the other hand, species such as *Chenopodium vulvaria, Oxalis acetosella*, and *Calamagrostis epigejos* were consumed in proportions that closely matched their field abundance, indicating neutral selection. These may function as fallback forage species, especially during periods of resource scarcity when preferred plants are less available.

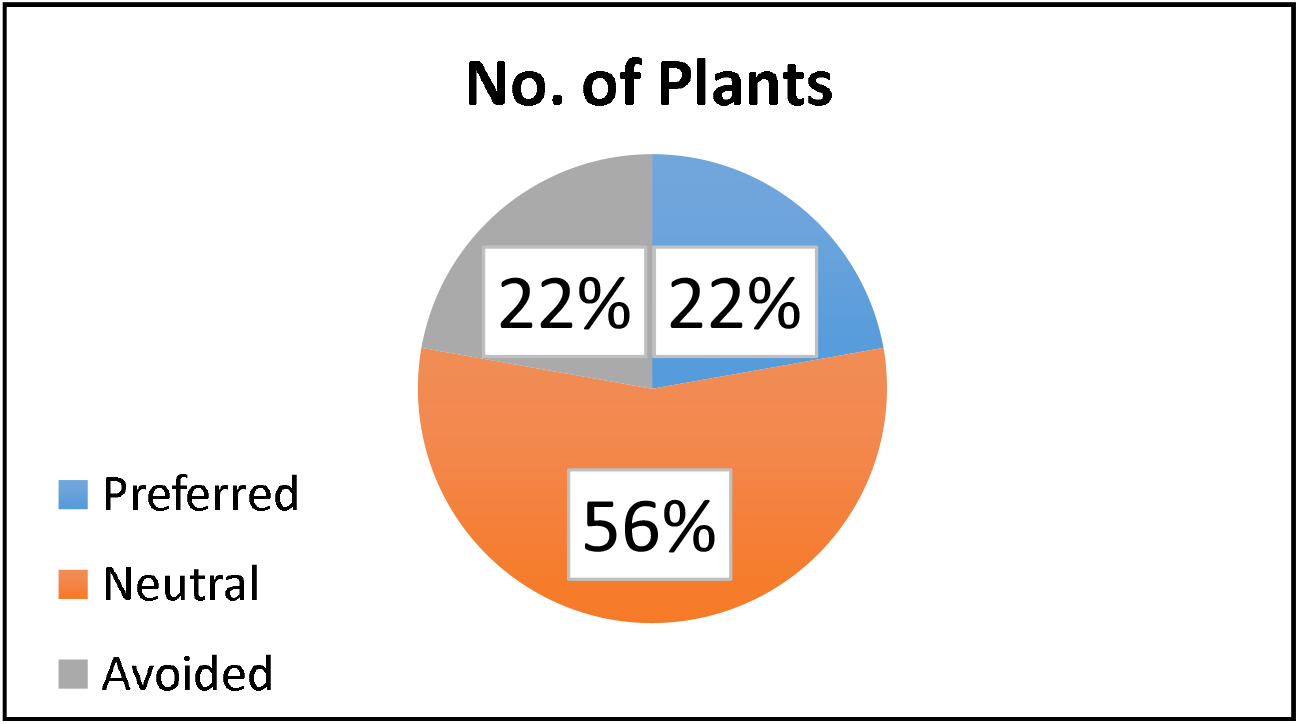

The observed dietary patterns were consistent across all three study sites, underscoring the blackbuck’s ability to forage selectively within resource-constrained semi-arid environments during winter. The preference for nutrient-dense forage species such as *Cynodon dactylon*, alongside the active avoidance of fibrous or chemically defended plants, reflects a foraging strategy aimed at maximizing nutritional gain while minimizing the intake of suboptimal or potentially harmful vegetation. Such adaptive behavior is particularly critical in human-modified landscapes, where habitat quality and forage distribution are often heterogeneous due to agricultural expansion and habitat fragmentation.

The widespread occurrence and selective use of species like *C. dactylon* and *Poa annua* across all surveyed sites suggest that these taxa may function as ecological indicators of suitable blackbuck foraging habitats. Their presence could guide conservation interventions focused on habitat restoration and forage management.

This study further highlights the utility of microhistological fecal analysis as a robust, non-invasive method for assessing herbivore diet composition at the species level. Detailed understanding of blackbuck dietary preferences contributes valuable ecological insight into habitat use and can inform evidence-based management strategies. Specifically, maintaining a diverse assemblage of palatable native forage species while controlling the spread of invasive plants such as *Prosopis juliflora* is crucial for sustaining blackbuck populations in semi-arid agro-ecosystems.

## Discussion

The present study reveals that blackbuck (*Antilope cervicapra*) exhibit a distinct and selective foraging pattern during the winter season, driven by both nutritional optimization and ecological constraints. Micro-histological analysis of fecal samples, reinforced through Bonferroni-adjusted confidence intervals, clearly indicates that the species does not graze randomly but displays pronounced dietary preferences. Out of the 18 plant species identified across the landscape, certain grasses such as *Cynodon dactylon* and *Poa annua* were significantly preferred over their relative availability, underlining their crucial role as core forage species in the blackbuck’s winter diet.The preferential selection of *Cynodon dactylon*, a C4 perennial grass known for its high nutrient content, digestibility, and energetic efficiency, aligns with earlier findings that establish its value as a keystone forage species in semi-arid ecosystems (Jhala, 1997; Sankaran, 2005). These qualities make it particularly important during resource-limited periods, such as the dry winter season, when energy demands are high, and food quality is often compromised. The consistent selection of *Poa annua*, a cool-season annual grass, suggests an adaptive dietary response to seasonal availability, with blackbuck potentially targeting it for its softer texture and favorable leaf-to-stem ratio during colder months. Interestingly, certain low-abundance plant species, including *Chenopodium vulvaria*, were disproportionately consumed, despite their limited representation in the environment. This suggests that blackbuck may be employing a selective foraging strategy aimed not merely at maximizing caloric intake but also targeting specific micronutrients, minerals, or secondary plant metabolites that aid in physiological regulation or parasite resistance (Goyal et al., 1988; Robbins, 1993). The preference for such species may also reflect self-medicative behavior or compensatory foraging to meet mineral deficits, a strategy observed in several wild ungulate species inhabiting nutrient-stressed habitats.

Conversely, the avoidance of *Prosopis juliflora*, despite its ecological dominance across large patches of the study area, is ecologically noteworthy. This invasive woody species, although widely available, was underutilized by blackbuck, likely due to its poor palatability, physical defenses such as thorns, or presence of allelopathic and anti-nutritional compounds (Pasiecznik et al., 2001; Rai & Tripathi, 2018). The fact that such a prolific species contributes minimally to the diet suggests that availability alone does not dictate consumption. Instead, chemical and mechanical deterrents, coupled with poor nutritive quality, may actively discourage herbivory. This underlines the broader ecological risks posed by *Prosopis juliflora*, as its spread can lead to homogenization of the landscape, reducing the diversity of palatable and nutritionally valuable species, thereby lowering the carrying capacity of rangelands for native herbivores. Other species such as *Solanum nigrum, Lactuca floridana*, and *Parthenium hysterophorus* were also largely avoided, despite being moderately available in some transects. The selective avoidance of these plants is likely due to their content of alkaloids, nitrates, or other secondary toxic metabolites that can affect digestion, reproduction, or general health in large herbivores (Robbins, 1993; Kumar & Shahabuddin, 2005). Such dietary exclusions further reinforce the blackbuck’s capacity for selective avoidance based on evolved physiological and sensory mechanisms to detect and reject harmful forage. Several species, including *Eruca sativa* and *Rumex pulcher*, were consumed roughly in proportion to their availability in the landscape. This pattern of neutral selection may point to their role as fallback or buffer species, consumed when preferred forages are locally absent or seasonally depleted. Opportunistic foraging of this nature is consistent with behavioral flexibility observed in ungulates inhabiting unpredictable, semi-arid environments (Isvaran, 2007). The ability to shift between preferred and neutral forage may be crucial for maintaining energy balance in fluctuating environments, particularly in human-dominated landscapes where habitat fragmentation and grazing competition with livestock further complicate resource availability.

Overall, these findings provide compelling evidence that blackbuck employ a foraging strategy that seeks to optimize nutrient intake while minimizing ingestion of suboptimal or chemically defended plant species. This adaptive selectivity is ecologically significant, especially in landscapes under increasing anthropogenic pressure. By actively selecting for high-quality forage while avoiding ecologically dominant but nutritionally inferior invasives, blackbuck not only demonstrate behavioral plasticity but also signal the ecological importance of maintaining native grassland diversity. Conservation strategies must therefore prioritize the protection, restoration, and active propagation of high-value forage species such as *Cynodon dactylon* and *Poa annua*, while simultaneously controlling the spread of invasive taxa like *Prosopis juliflora*. This is essential for sustaining blackbuck populations and promoting the overall resilience of semi-arid grassland ecosystems.

## Conclusion

This study highlights the winter dietary preferences and selective foraging strategies of blackbuck (*Antilope cervicapra*) inhabiting the semi-arid grasslands of Aligarh, Uttar Pradesh. Utilizing micro-histological analysis of fecal pellets, in conjunction with Bonferroni-adjusted confidence intervals, the research provides strong evidence that blackbuck exhibit non-random foraging behavior and display a clear preference for specific plant species based on nutritional content and palatability. Notably, grasses such as *Cynodon dactylon* and *Poa annua* were significantly overrepresented in the diet compared to their proportional availability in the field. This indicates a strong positive selection for these nutrient-rich species, which likely serve as primary forage resources during the dry winter season when food quality and quantity decline (Jhala, 1997; Sankaran, 2005).

*Cynodon dactylon*, a C4 grass with high digestibility and energy content, has been consistently identified in previous studies as a preferred species due to its year-round availability and resilience under grazing pressure. Similarly, *Poa annua*, a cool-season grass, offers soft, palatable foliage that complements the nutritional needs of blackbuck during winter. Their consistent selection underlines the ecological importance of maintaining such species in grassland ecosystems.Conversely, the study found that *Prosopis juliflora*, an invasive and ecologically dominant species in the region, was significantly underutilized. Despite its abundance, blackbuck largely avoided it, likely due to its thorny morphology, unpalatable nature, and the presence of chemical defenses or anti-nutritional compounds (Pasiecznik et al., 2001; Rai & Tripathi, 2018). Its widespread presence and limited contribution to the blackbuck diet signal its negative impact on forage quality and biodiversity. Certain low-availability species like *Chenopodium vulvaria* were selectively consumed, suggesting that blackbuck may actively seek out less common species to fulfill micronutrient requirements or to access beneficial secondary compounds (Goyal et al., 1988; Robbins, 1993). Such targeted foraging indicates a strategic dietary behavior rather than opportunistic grazing.

Overall, these findings underscore that blackbuck are not generalist grazers but highly selective feeders. Their dietary choices are influenced by both forage quality and ecological availability. Therefore, conservation and grassland management strategies must prioritize the protection of native, nutrient-rich species while simultaneously curbing the spread of invasive plants like *Prosopis juliflora*. Ensuring a diverse and high-quality plant community is essential to support the long-term viability of blackbuck populations in semi-arid landscapes.

### Conservation and Future Recommendations

**Invasive Species Management:** Control or remove *Prosopis juliflora* from key blackbuck habitats through mechanical or biological methods to prevent further degradation of native forage diversity (Pasiecznik et al., 2001).

**Native Grassland Restoration**: Reintroduce and promote native C4 grasses such as *Cynodon dactylon, Bothriochloa spp*., and *Heteropogon spp*. that offer high nutritional value and support ungulate populations (Rodgers & Panwar, 1988).

**Protection of Microhabitats**: Conserve patches that support rare but selectively consumed plant species, ensuring access to essential micronutrients and secondary plant compounds (Robbins, 1993).

**Seasonal Diet Monitoring**: Conduct year-round fecal analysis to detect seasonal shifts in diet and adjust management strategies accordingly (Singh et al., 2014).

**Community Participation**: Engage local communities in sustainable grazing, restoration, and monitoring programs to reduce anthropogenic pressure and enhance conservation outcomes (Kumar & Shahabuddin, 2005).

## Supporting information

Blackbuck_Supplementary_Data_Yunus.zip

---

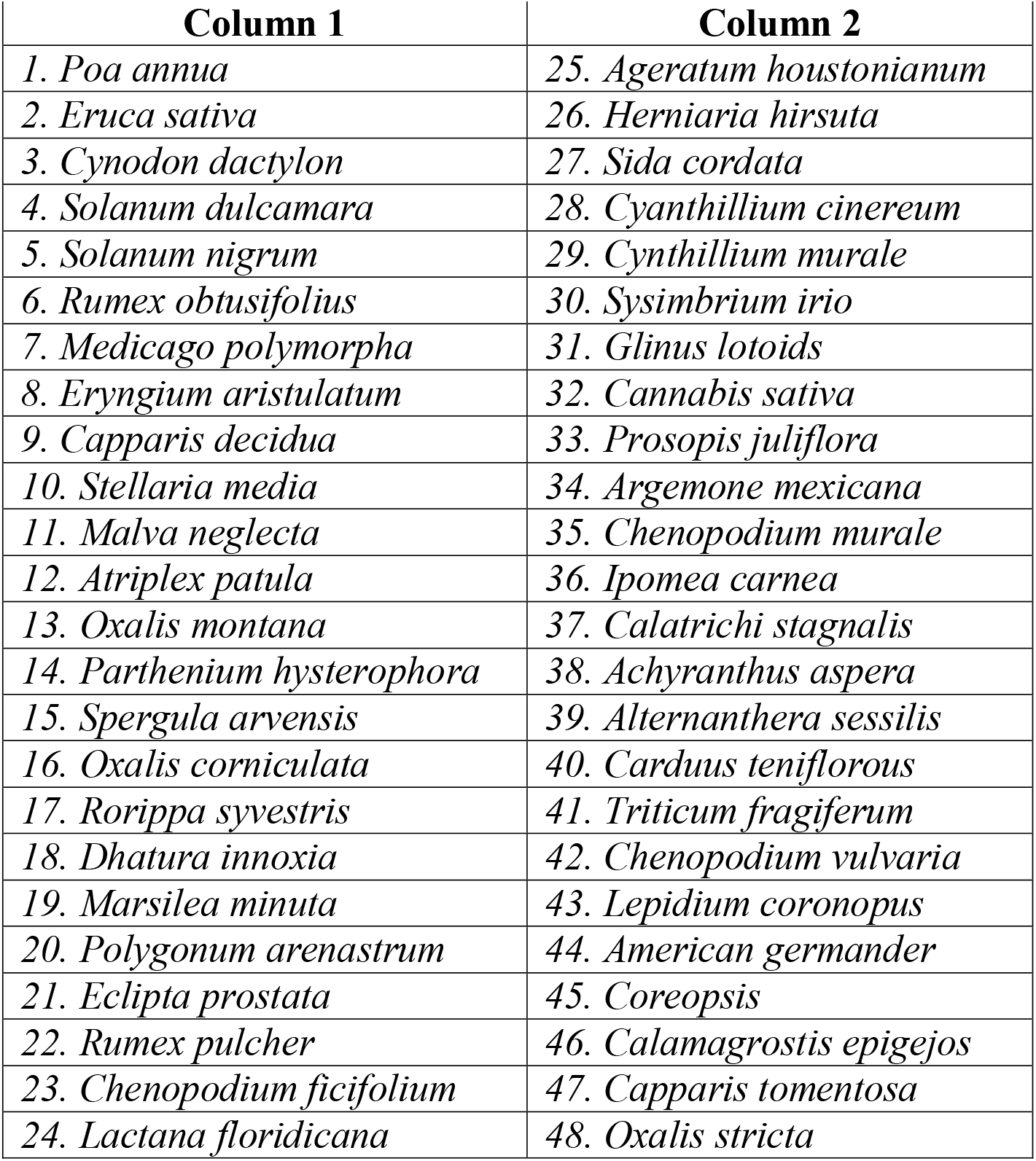
1. VEGETATION SAMPLING.

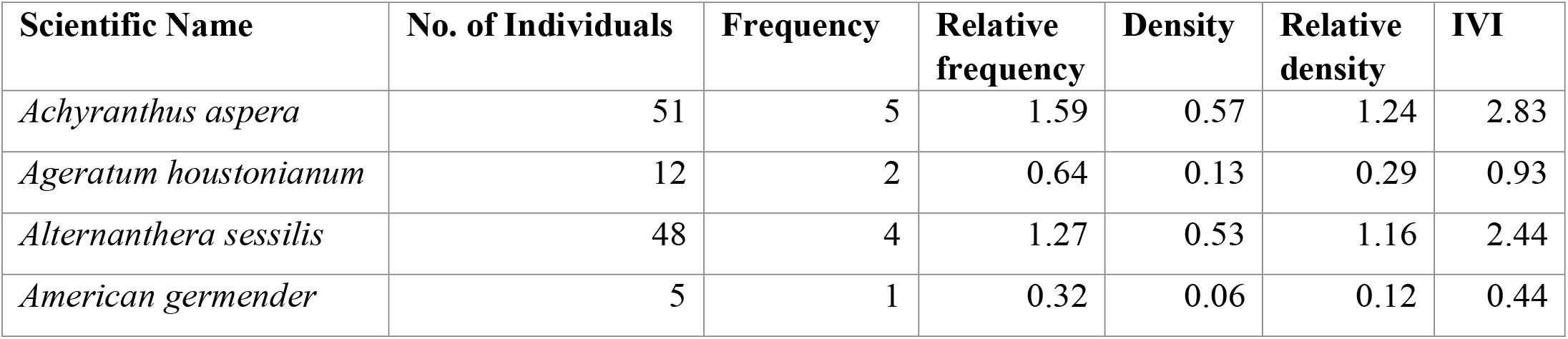
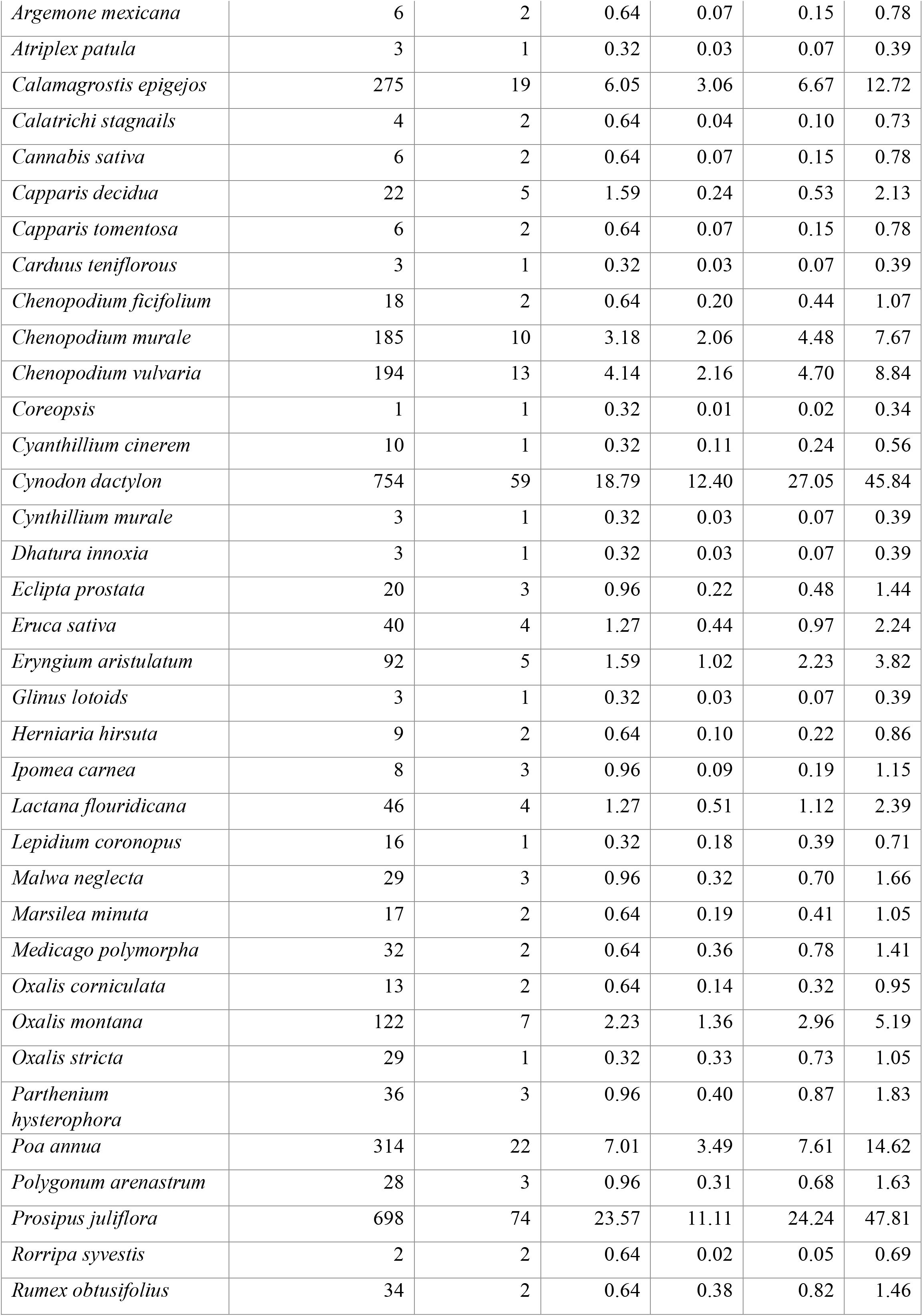
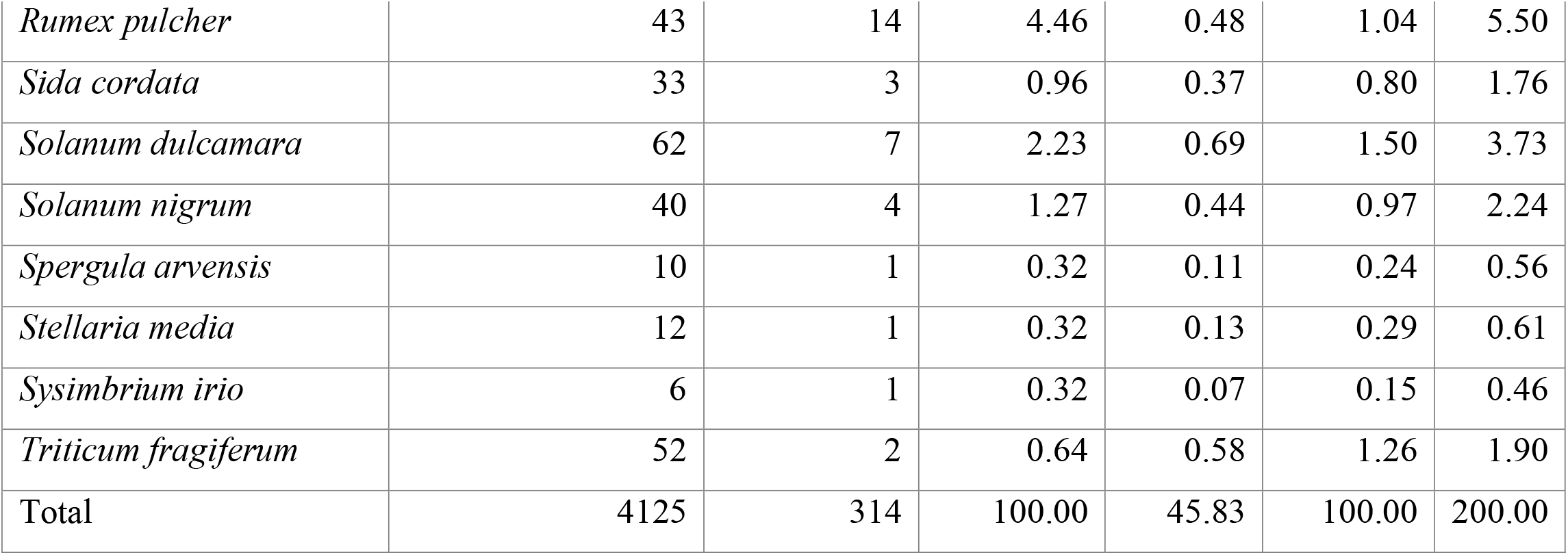
2. IVI OF PLANTS.

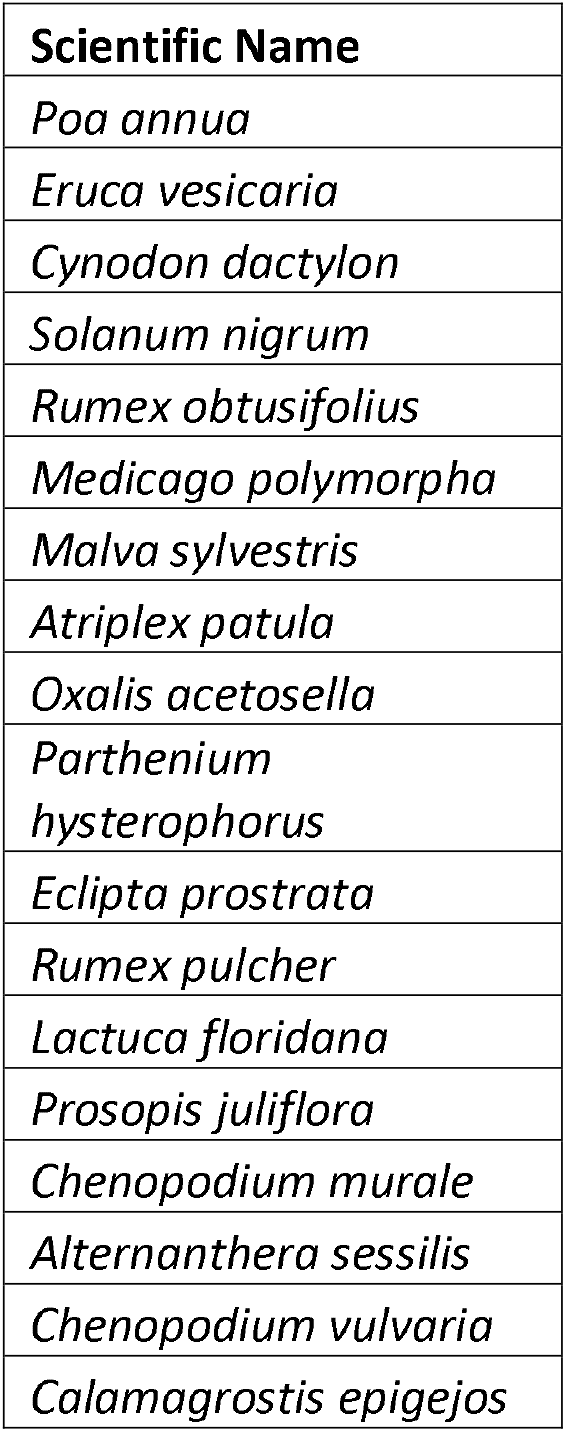
3. TOTAL PLANTS ARE FOUND IN PALLETS.

3. TOTAL PLANTS ARE FOUND IN PALLETS
| Scientific Name |
| --- |
| *Poa annua* |
| *Eruca vesicaria* |
| *Cynodon dactylon* |
| *Solanum nigrum* |
| *Rumex obtusifolius* |
| *Medicago polymorpha* |
| *Malva sylvestris* |
| *Atriplex patula* |
| *Oxalis acetosella* |
| *Parthenium hysterophorus* |
| *Eclipta prostrata* |
| *Rumex pulcher* |
| *Lactuca floridana* |
| *Prosopis juliflora* |
| *Chenopodium murale* |
| *Alternanthera sessilis* |
| *Chenopodium vulvaria* |
| *Calamagrostis epigejos* |

4. BONFERRONI CONFIDENCE INTERVAL TABLE.

4. BONFERRONI CONFIDENCE INTERVAL TABLE
| Plant Species | Fragments | Usage % | Availability % | Lower CI | Upper CI | Preference |
| --- | --- | --- | --- | --- | --- | --- |
| Annual Meadowgrass | 91 | 0.104 | 0.0746 | 0.078 | 0.131 | Preferred |
| Aragula | 15 | 0.017 | 0.0147 | 0.006 | 0.028 | Neutral |
| Bermuda Grass | 305 | 0.349 | 0.2771 | 0.307 | 0.391 | Preferred |
| Black Nightshade | 1 | 0.001 | 0.0147 | -0.002 | 0.004 | Avoided |
| Broad-Leaved Dock | 10 | 0.011 | 0.0125 | 0.002 | 0.021 | Neutral |
| Burr Medic | 6 | 0.007 | 0.0118 | 0.000 | 0.014 | Neutral |
| Common Mallow | 9 | 0.010 | 0.0107 | 0.002 | 0.019 | Neutral |
| Common Orache | 12 | 0.014 | 0.0011 | 0.004 | 0.024 | Preferred |
| Common Wood Sorrel | 29 | 0.033 | 0.0316 | 0.018 | 0.049 | Neutral |
| Congress Grass | 4 | 0.005 | 0.0132 | -0.001 | 0.010 | Avoided |
| False Daisy | 3 | 0.003 | 0.0074 | -0.002 | 0.009 | Neutral |
| Fiddle Dock | 35 | 0.040 | 0.0158 | 0.023 | 0.057 | Preferred |
| Florida Lettuce | 3 | 0.003 | 0.0169 | -0.002 | 0.009 | Avoided |
| Mesquite | 151 | 0.173 | 0.2565 | 0.140 | 0.206 | Avoided |
| Nettle-Leavedgoosefoot | 36 | 0.041 | 0.0485 | 0.024 | 0.059 | Neutral |
| Sessile Joyweed | 23 | 0.026 | 0.0176 | 0.012 | 0.040 | Neutral |
| Stinging Goosefoot | 50 | 0.057 | 0.0595 | 0.037 | 0.077 | Neutral |
| Wood Small-reed | 77 | 0.088 | 0.0643 | 0.063 | 0.113 | Neutral |

**5. Photographic Records of Plant Species and Stomatal Impressions Identified from Blackbuck (Antilope cervicapra) Pellet Analysis**

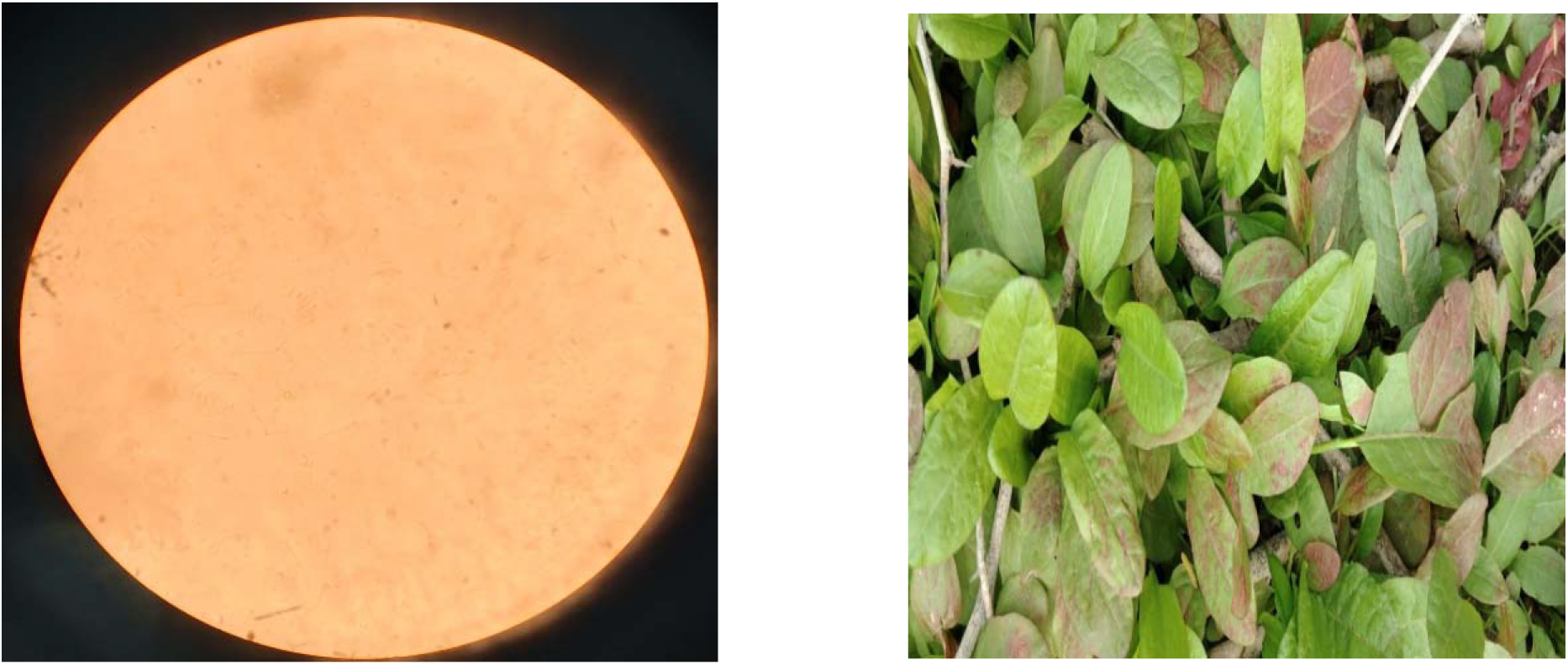
Rumex pulcher **(Fiddle Dock)**

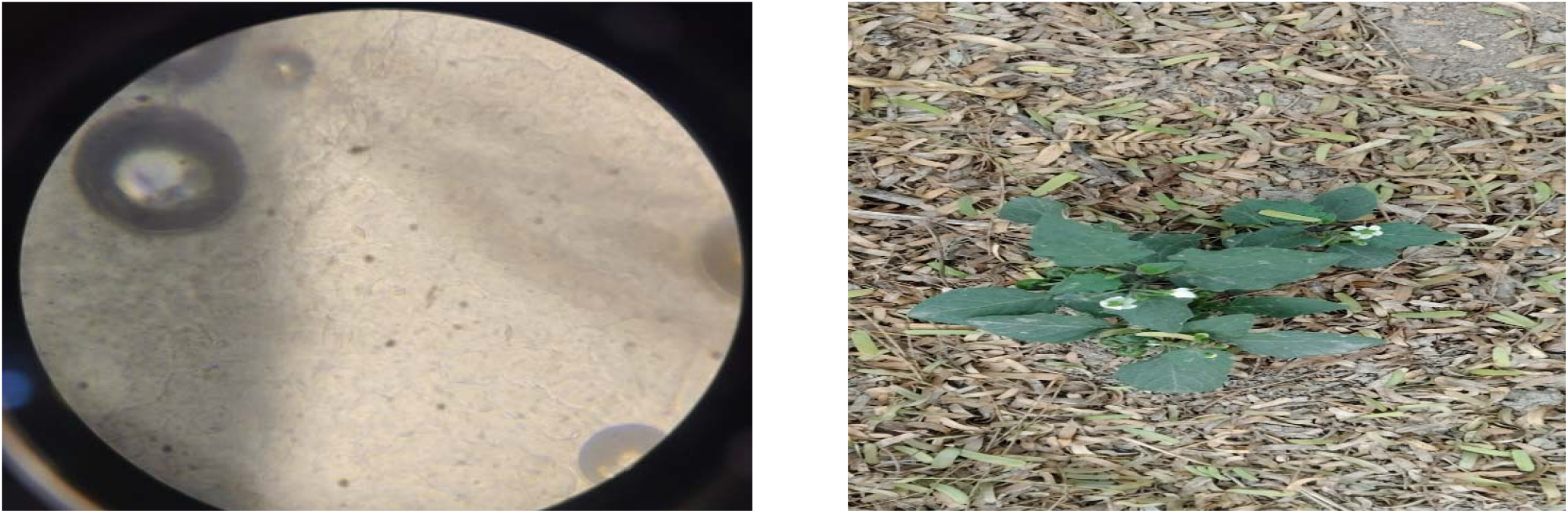
**Stomatal Type: Anomocytic**. **Distribution**: Primarily **hypostomatic**.

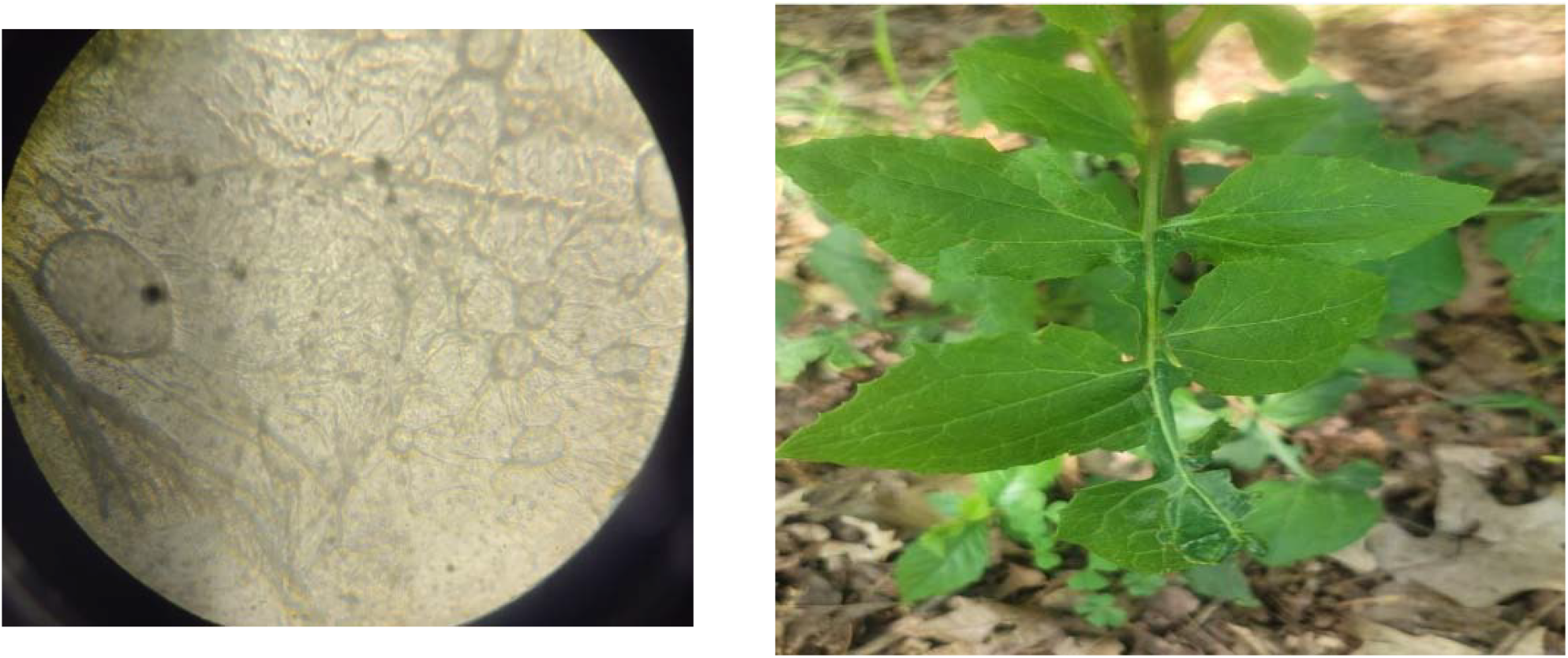
***Pictures 3 Lactana flouridicana*** **(Black Nightshade)** **Stomatal Type**: **Anisocytic** (surrounded by three unequal subsidiary cells). **Distribution**: **Amphistomatic**. **Additional Features**: Presence of glandular trichomes.

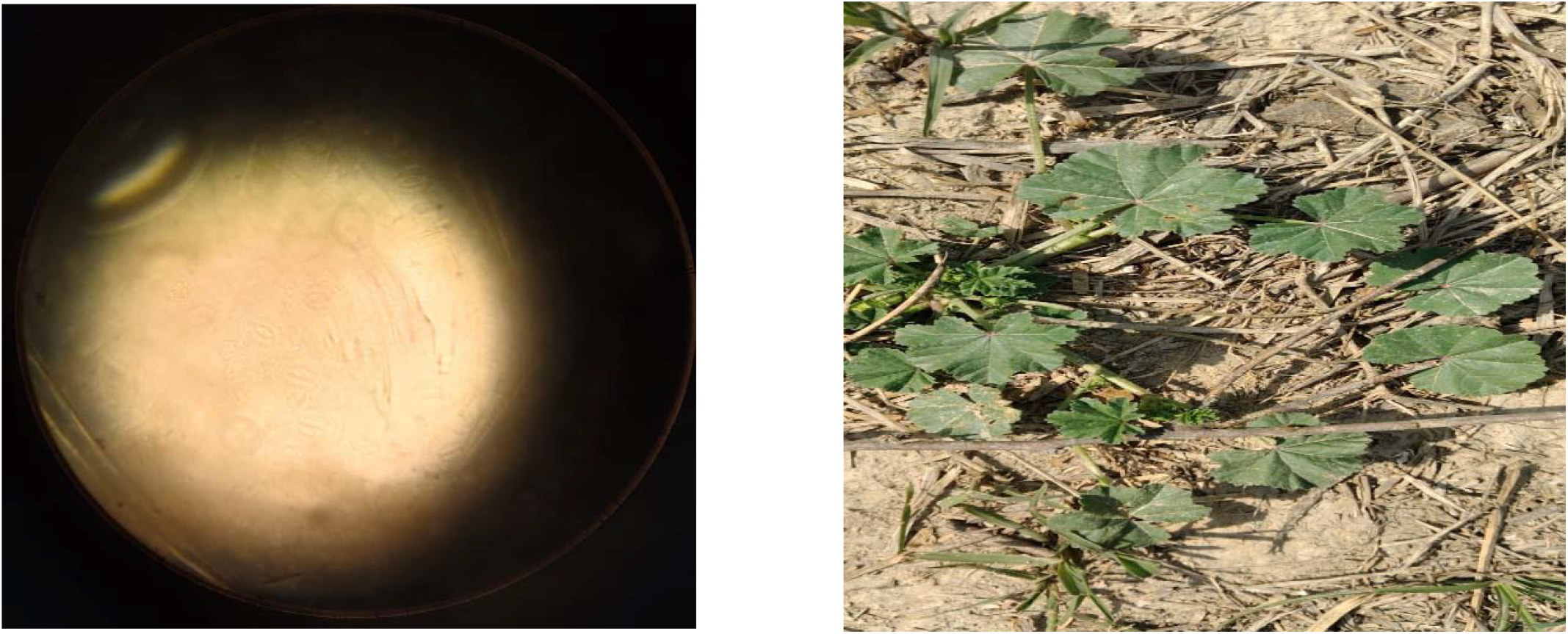
**Pictures 4 *Malwa sylvestris*** **(Common Mallow)** **Stomatal Type**: **Anisocytic. Distribution**: **Amphistomatic**

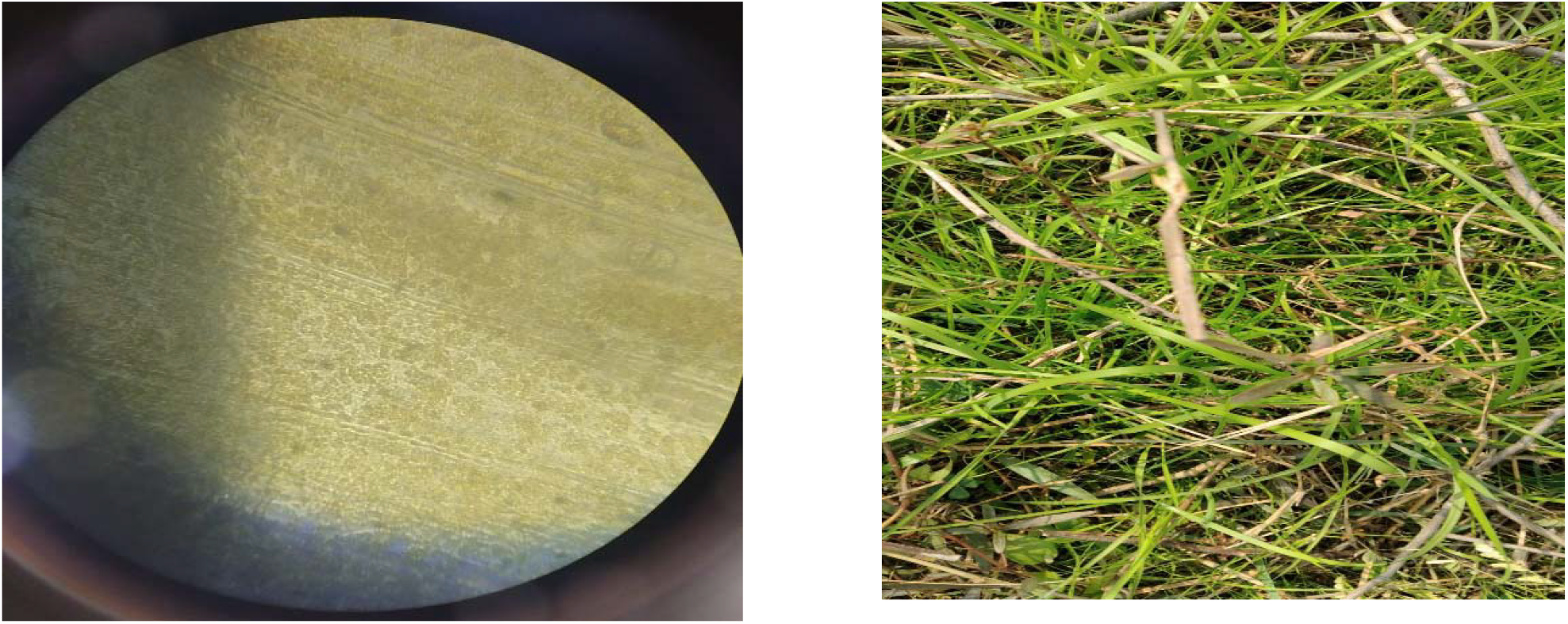
**Picture 5 *Cynodon dactylon*** **Bermuda grass** **Stomatal Type: Paracytic** (each stoma accompanied by two subsidiary cells parallel to the pore). **Distribution**: **Amphistomatic**. **Additional Features:** Epidermal cells are sinuous; silica bodies present

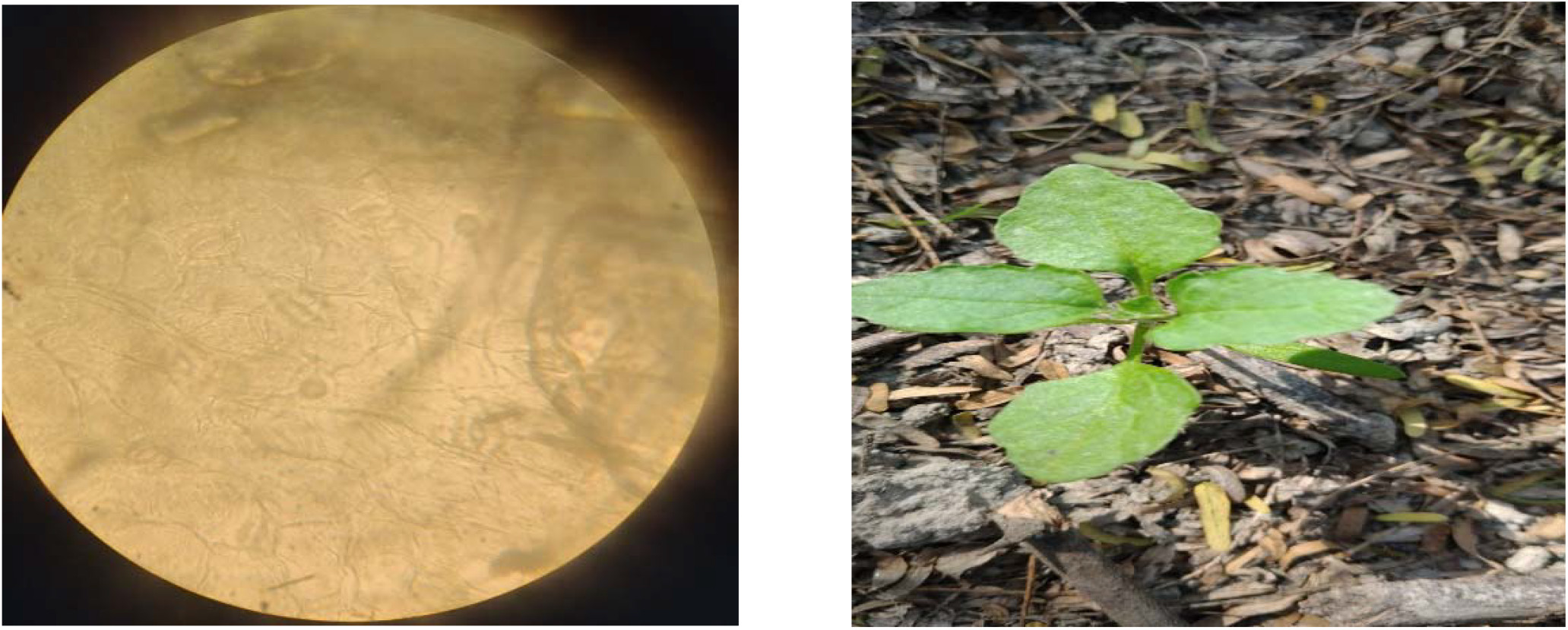
**Pictures *6 Chenopodium vulvaria*** (Stinking Goosefoot) **Stomatal Type**: **Anisocytic. Distribution**: **Amphistomatic**.

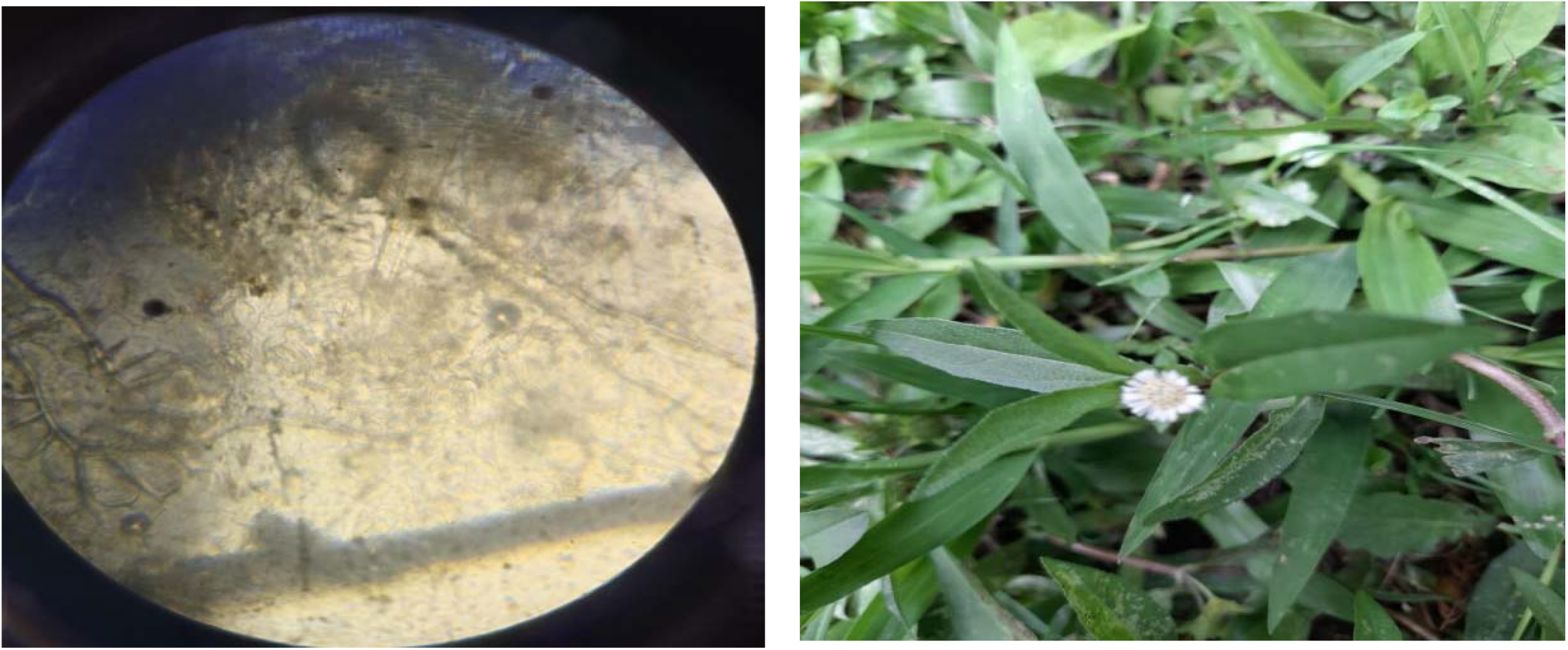
Picture 7 ***Eclipta prostrata*** (**False Daisy**) **Stomatal Type**: Anomocytic. **Distribution**: **Amphistomatic**. **Additional Features**: Presence of capitate glandular trichomes.

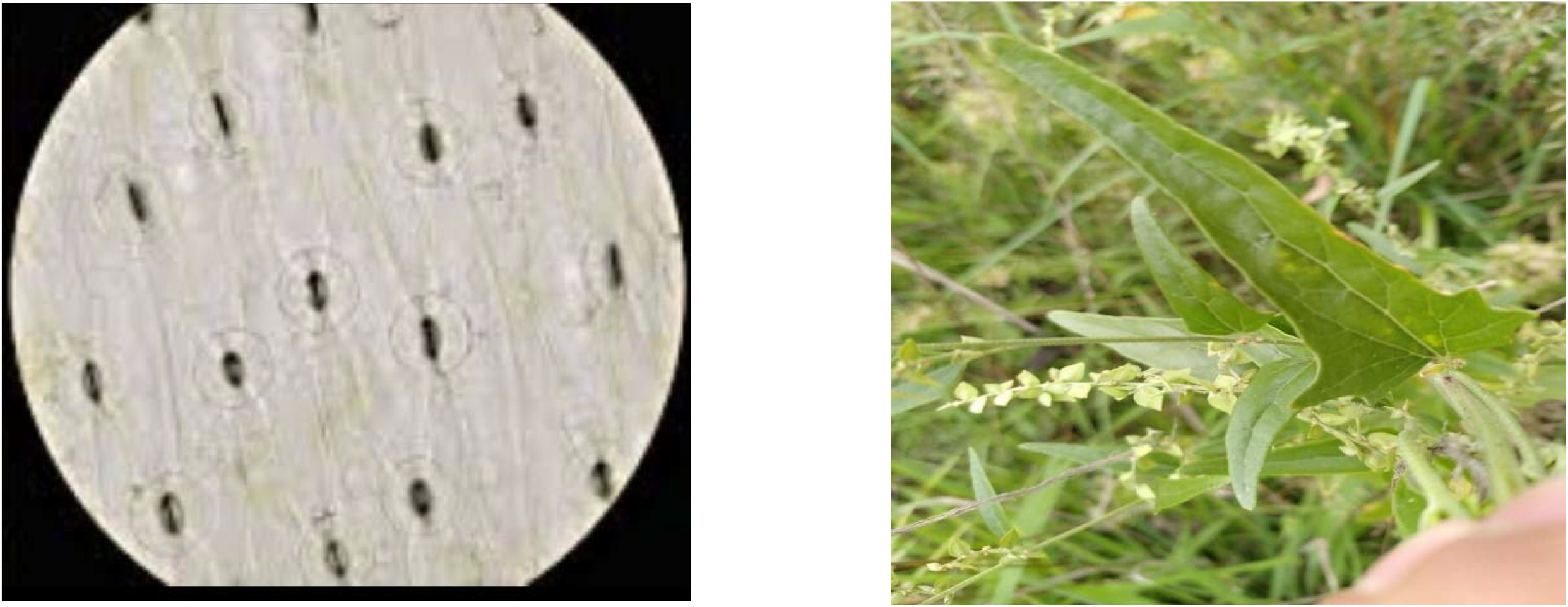
***Picture 8 Atriplex patula*** **(Common Orache)** **Stomatal Type**: **Anomocytic. Distribution**: **Amphistomatic**.

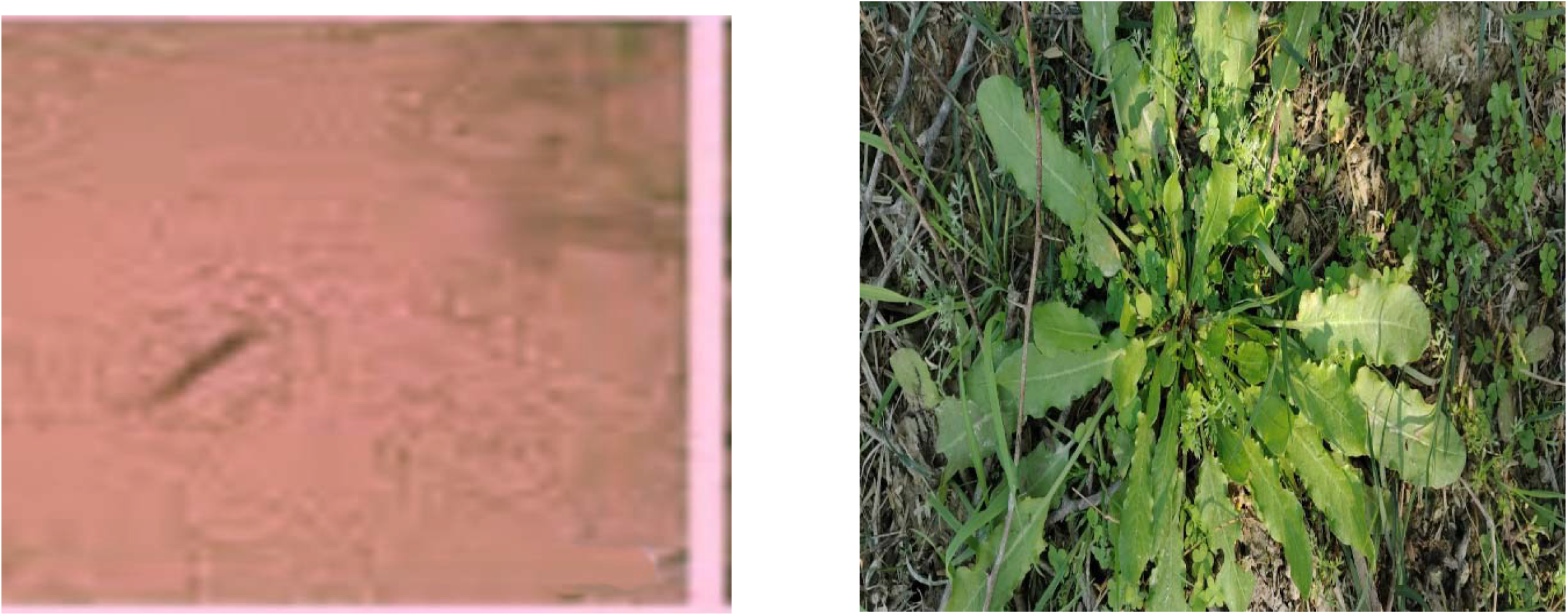
Picture 9 ***Rumex obtusifolius*** **(Broad-leaved Dock)** **Stomatal Type**: **Anomocytic**. **Distribution**: Primarily **hypostomatic**

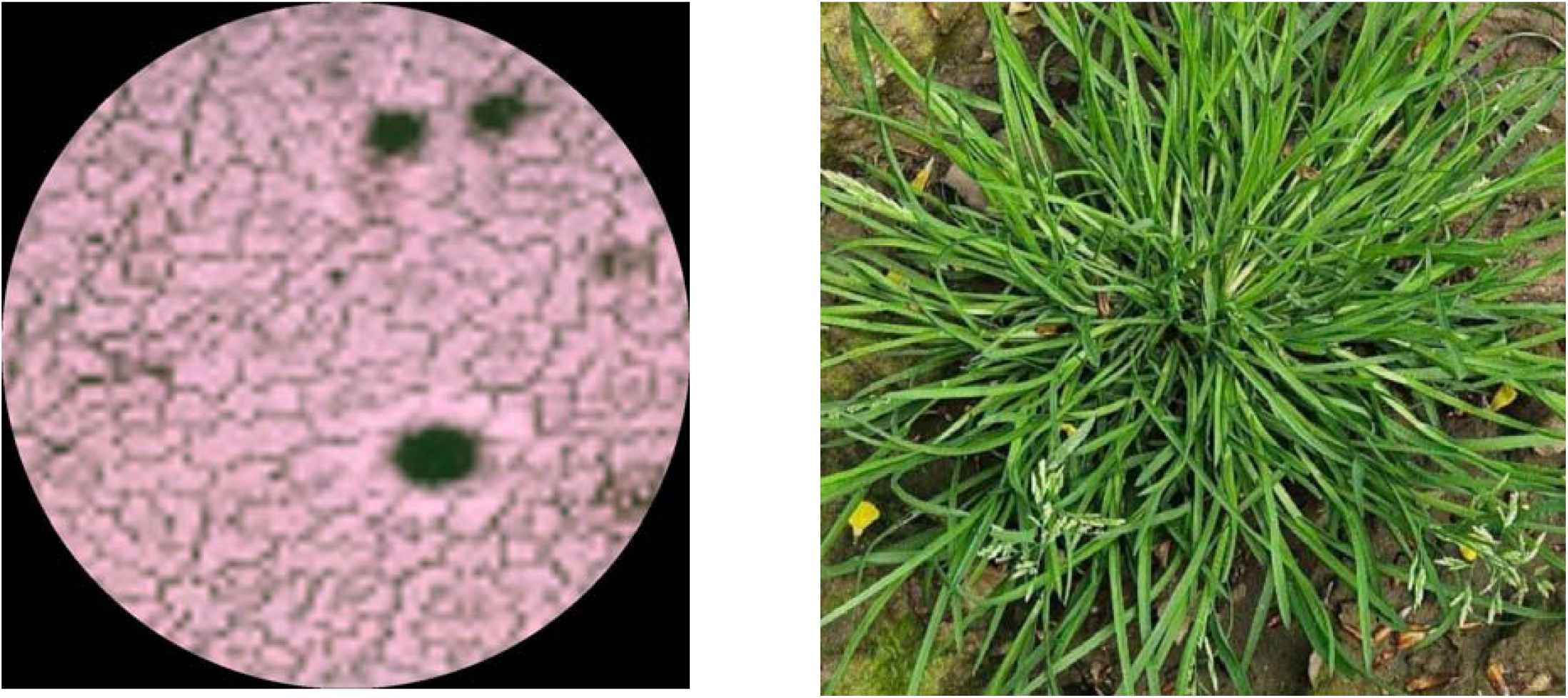
Picture 10 ***Poa annua*** (Annual Bluegrass) Stomatal Type: Typically dumbbell-shaped guard cells, common in grasses. Distribution: Amphistomatic (stomata on both leaf surfaces). Additional Features: Stomata are arranged in parallel rows between veins.

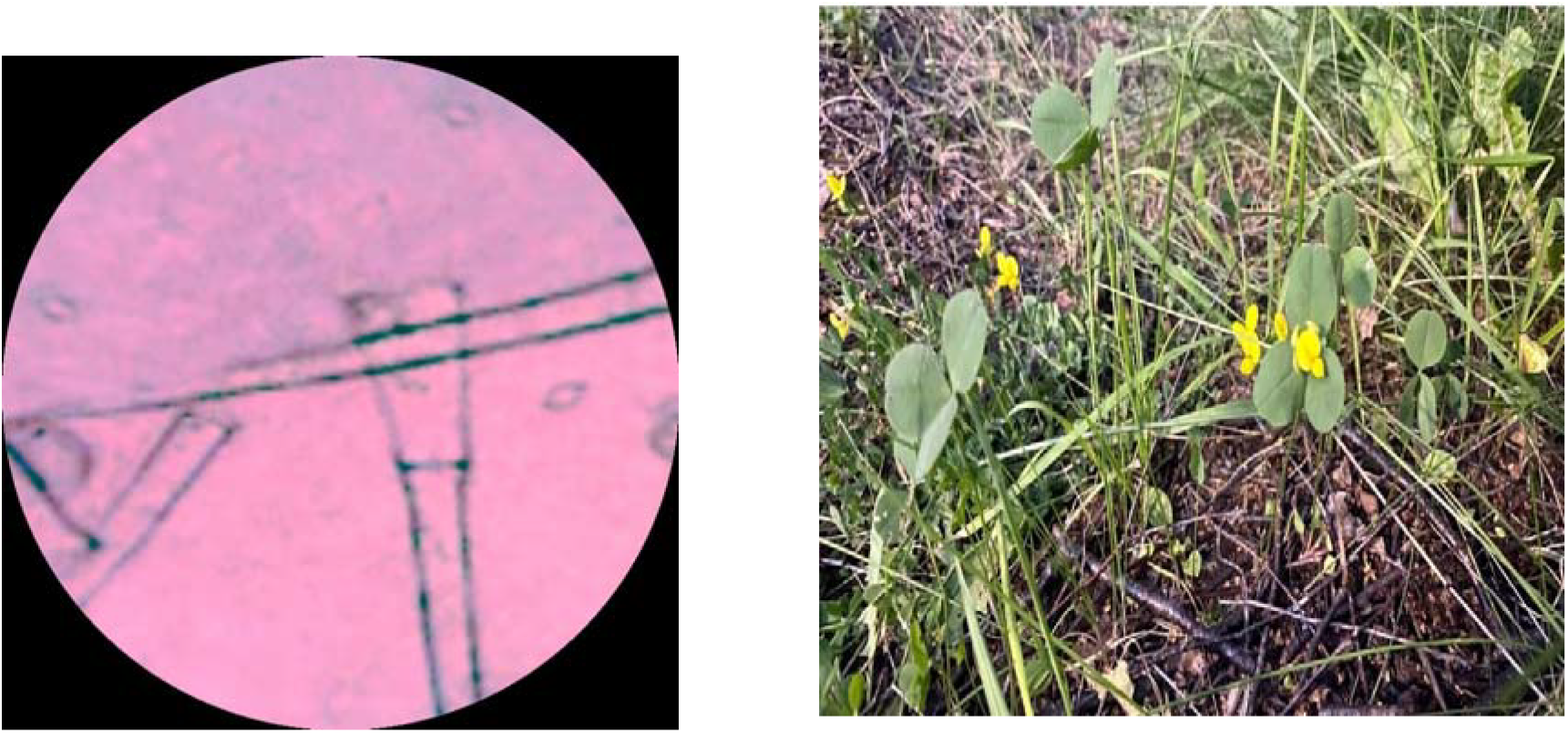
Picture ***11 Medicago polymorpha*** (Burr Medic) **Stomatal Type**: **Paracytic. Distribution**: **Amphistomatic**

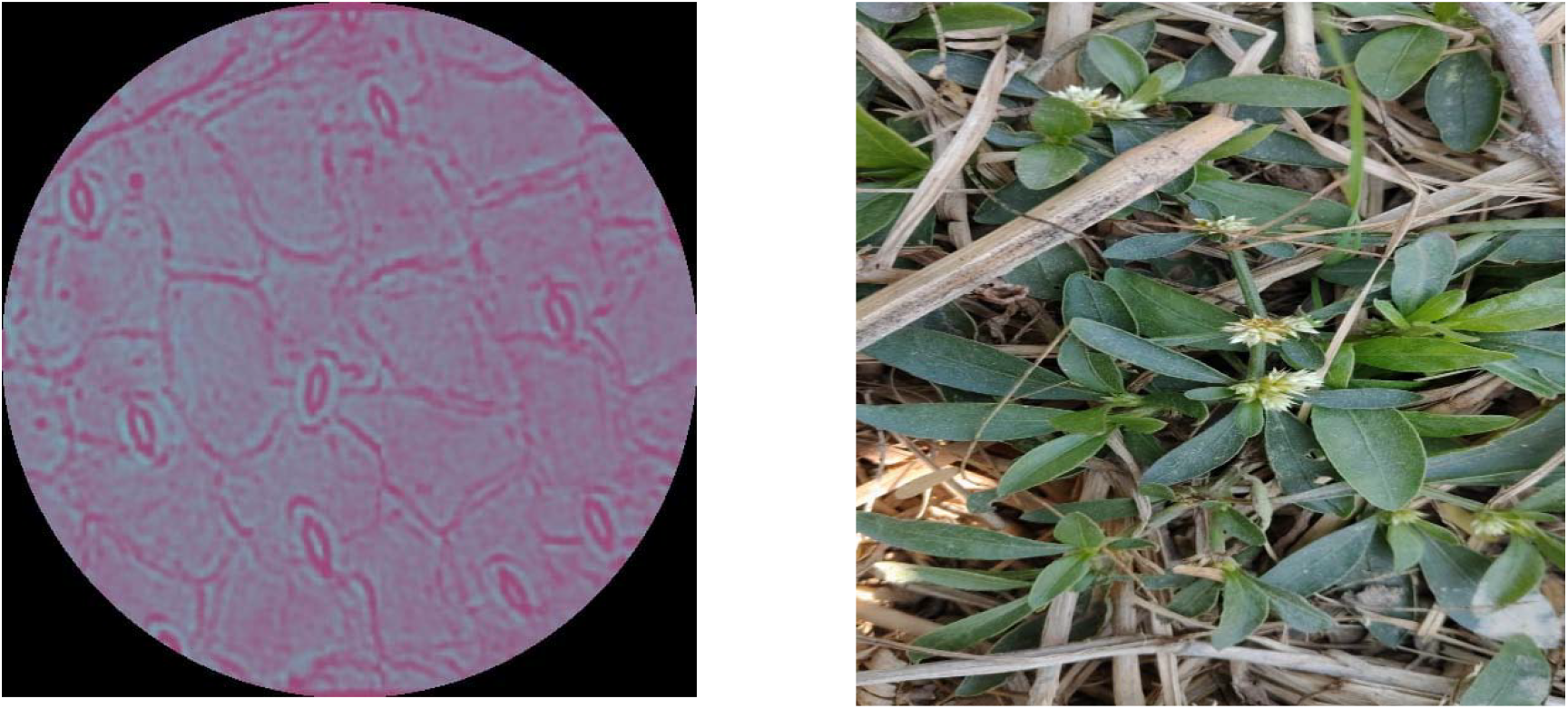
Pictre 12 ***Alternanthera sessillis*** (Sessile Joyweed) **Stomatal Type**: **Paracytic. Distribution**: **Amphistomatic**

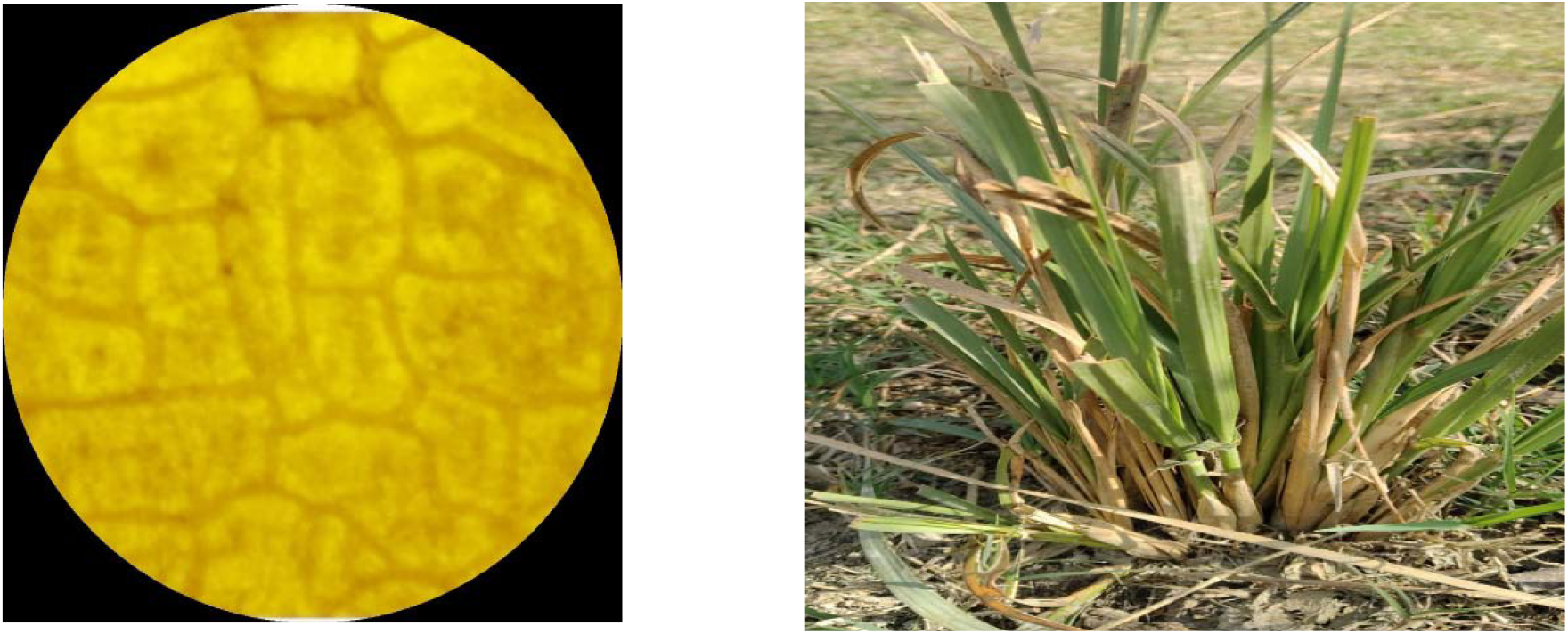
**Picture 13 *Calamagrostis epigejos*** (Wood Small-reed) **Stomatal Type**: **Dumbbell-shaped** guard cells, typical of grasses. **Distribution**: **Amphistomatic**.

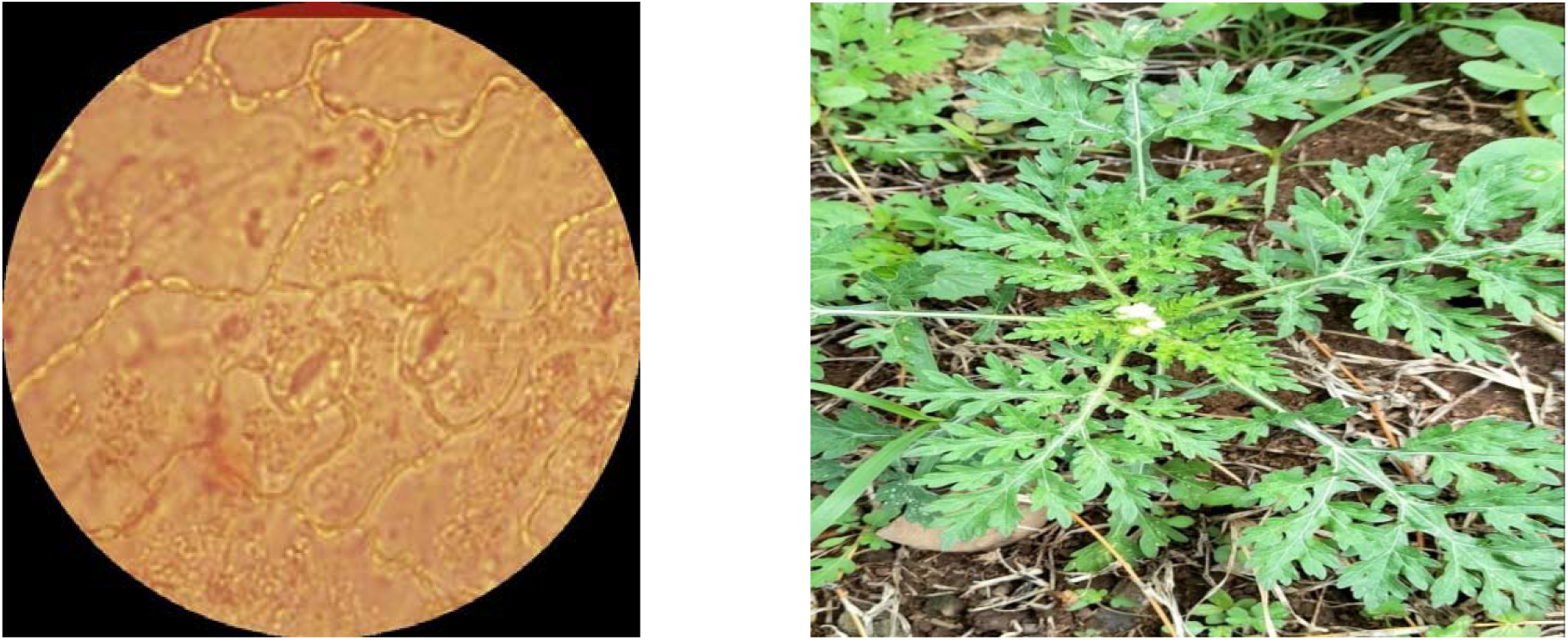
Picture ***14 Parthenium hysteriophorous*** (Santa Maria Feverfew) **Stomatal Type**: **Tetracytic** (four subsidiary cells, one on each side and two at the ends).

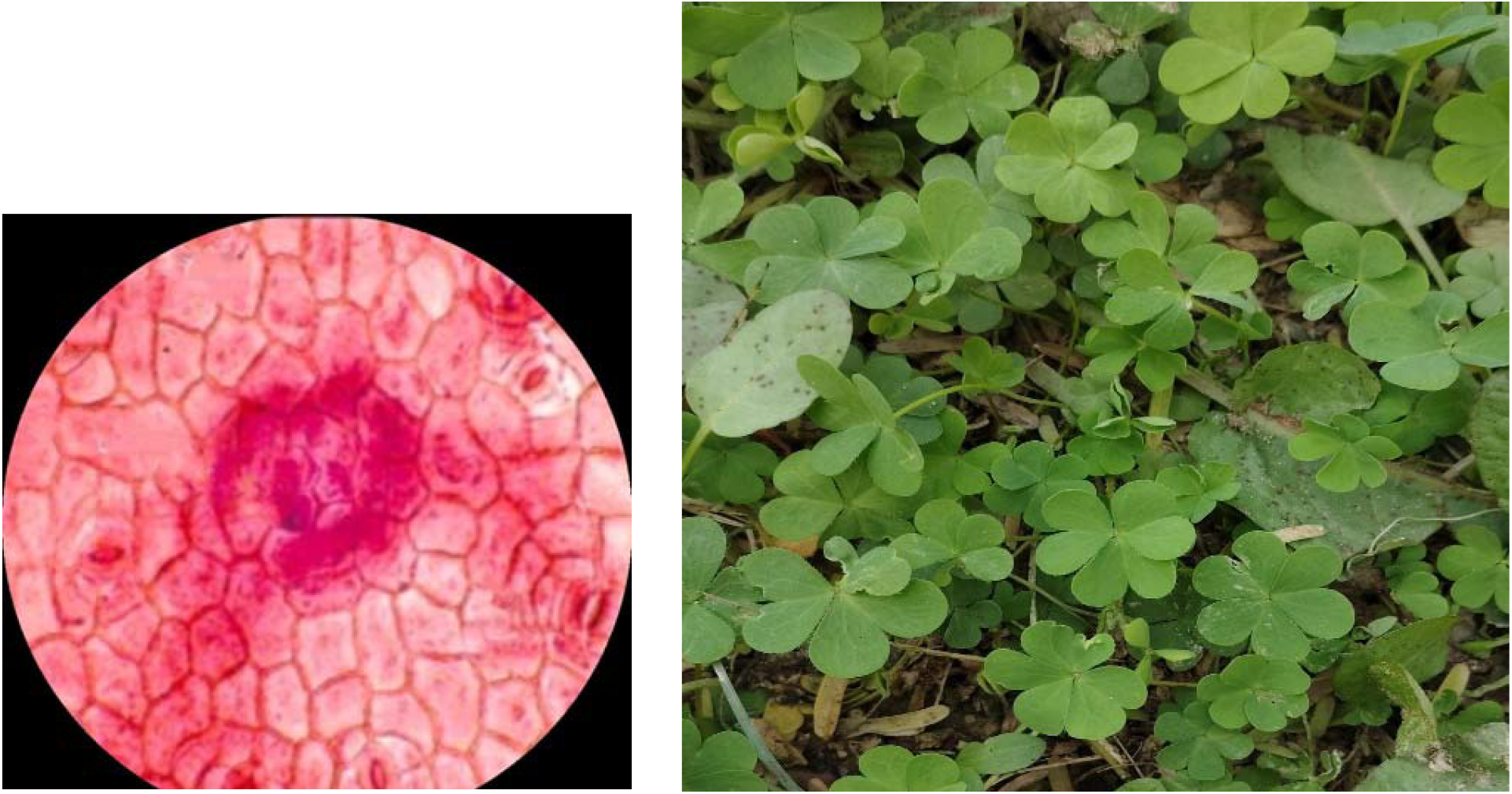
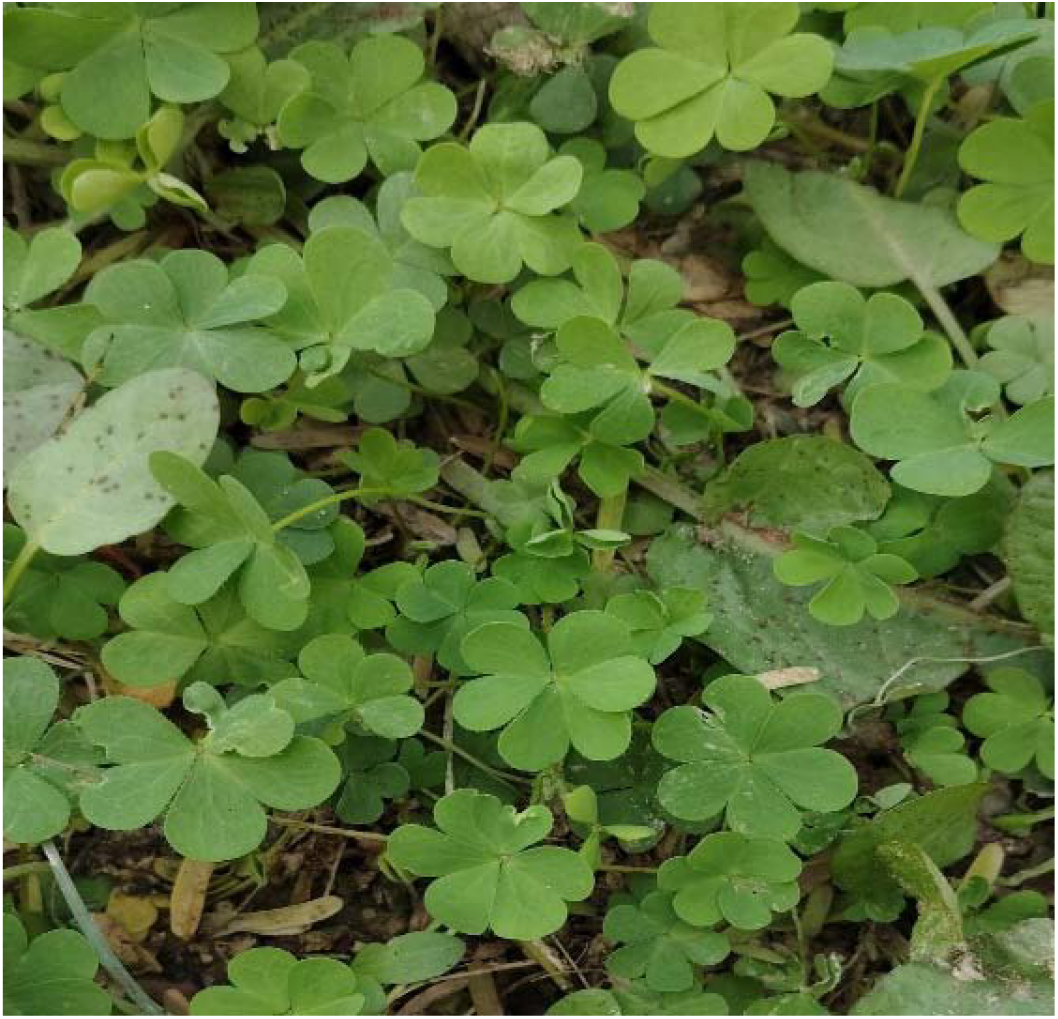
Picture 15 ***Oxalis acetosella*** (Wood Sorrel) **Stomatal Type**: **Anomocytic.** **Distribution**: **Amphistomatic**.

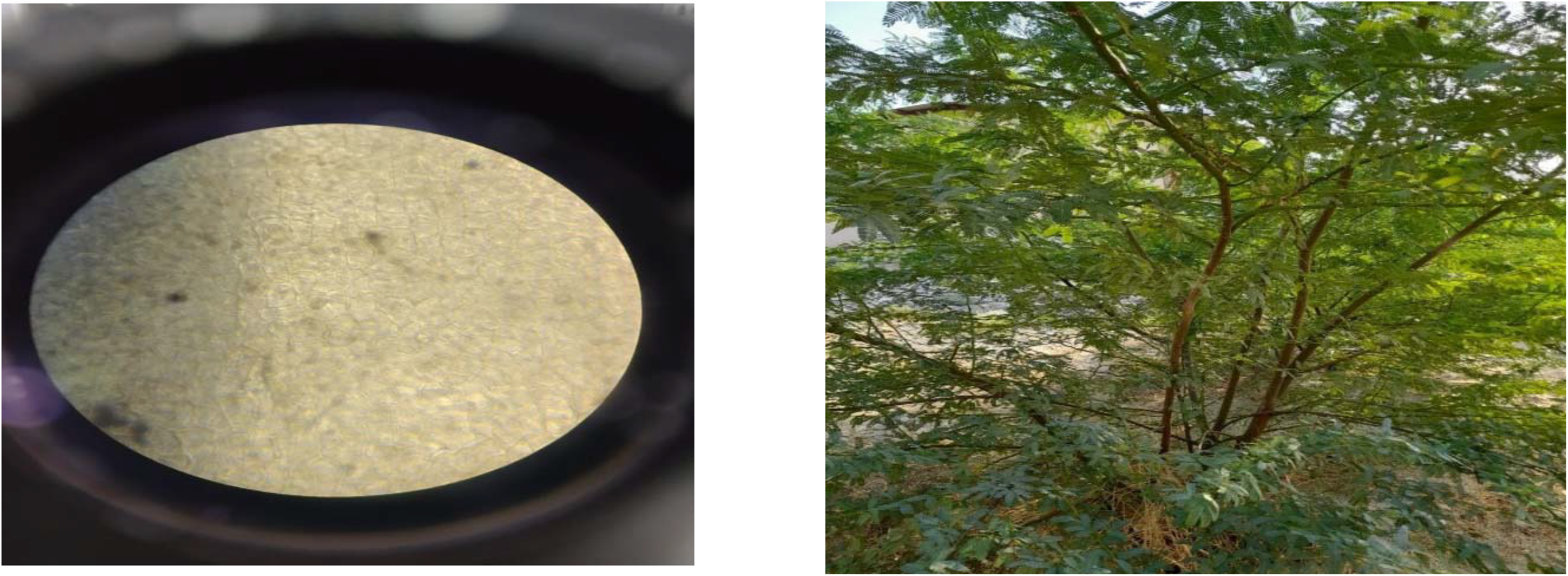
Picture **16 *Prosipus juliflora*** (Mesquite) **Stomatal Type**: **Anomocytic**.

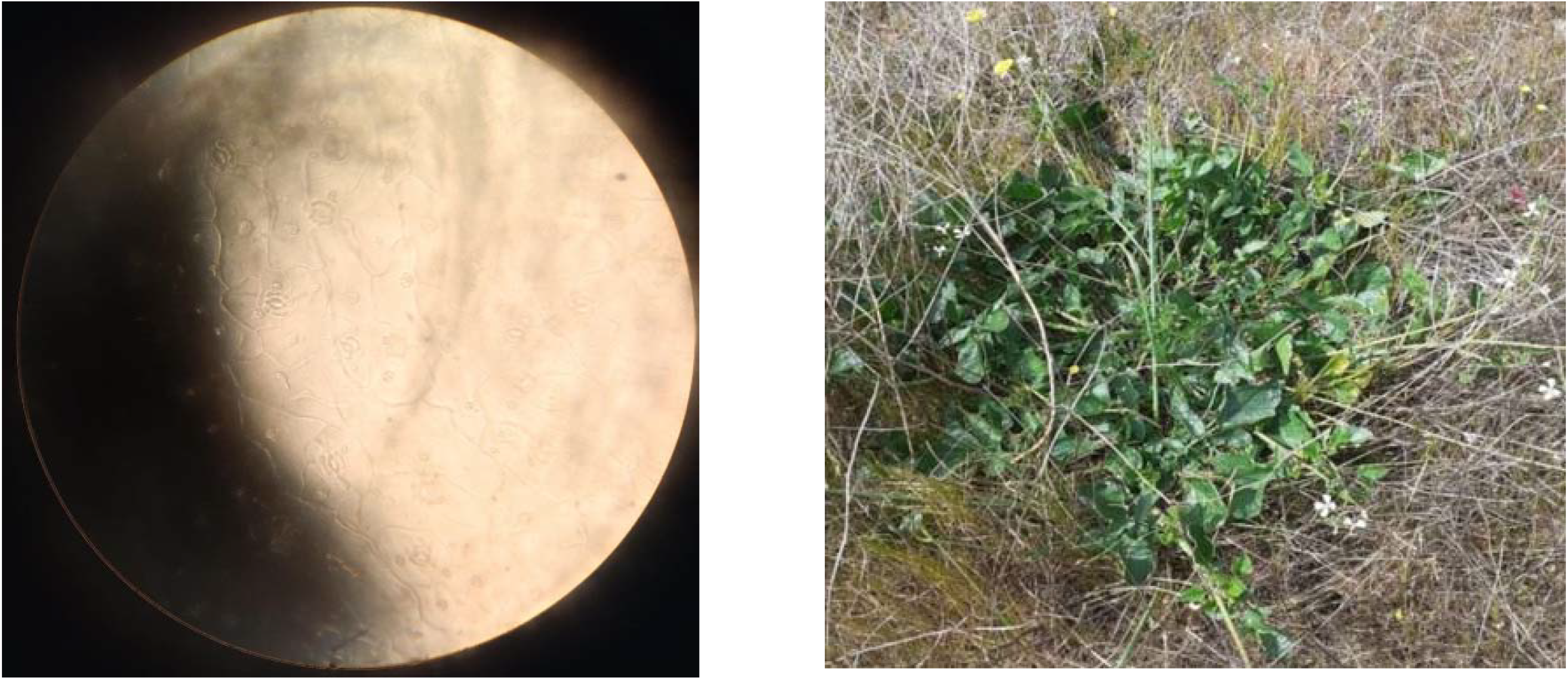
Picture 17 ***Eruca vesicaria*** **Distribution**: **Amphistomatic**

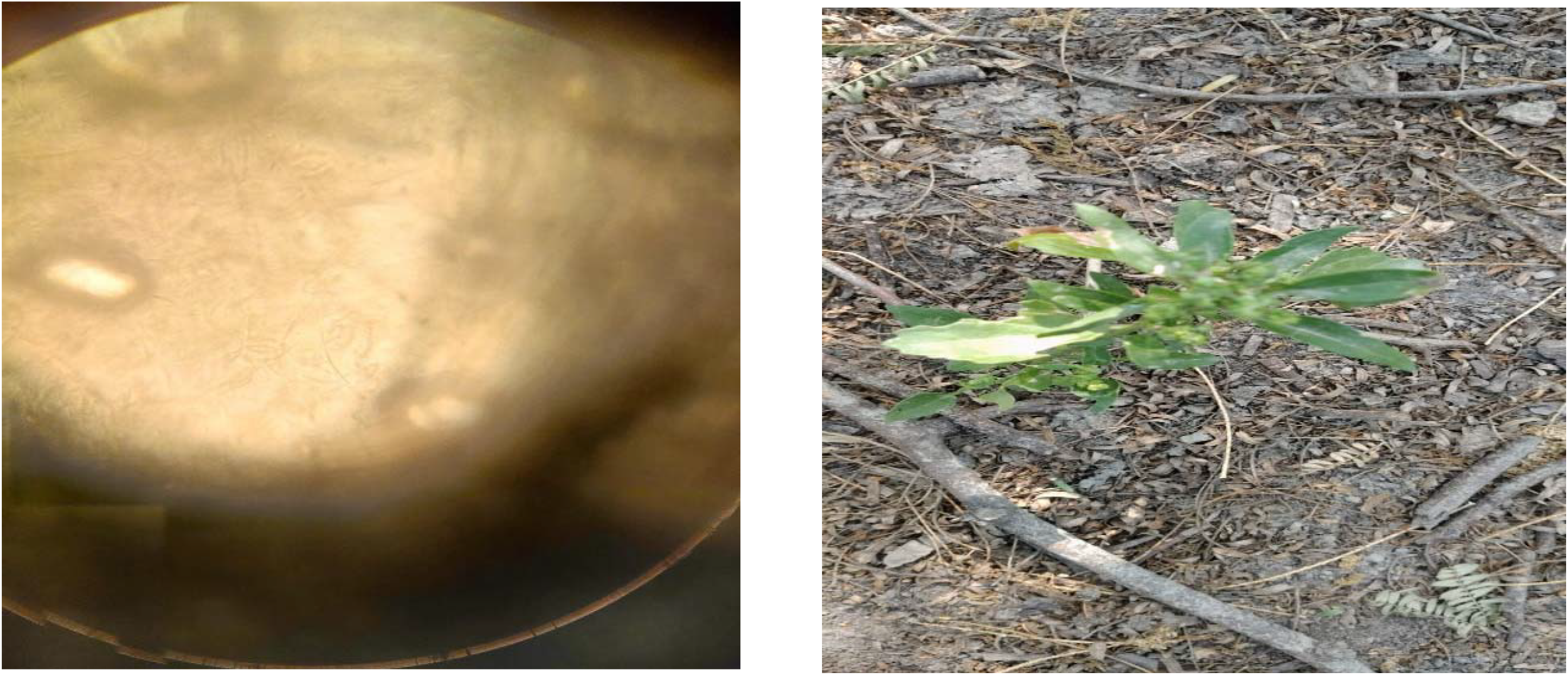
18. ***Chenopodium murale*** (Nettle-leaved Goosefoot) Stomatal Type: **Anisocytic**. Distribution: **Amphistomatic**.

