## Supplementary material for "Winter Foraging Selectivity of Blackbuck in Semi-Arid Agricultural Landscapes": Blackbuck_Supplementary_Data_Yunus.zip

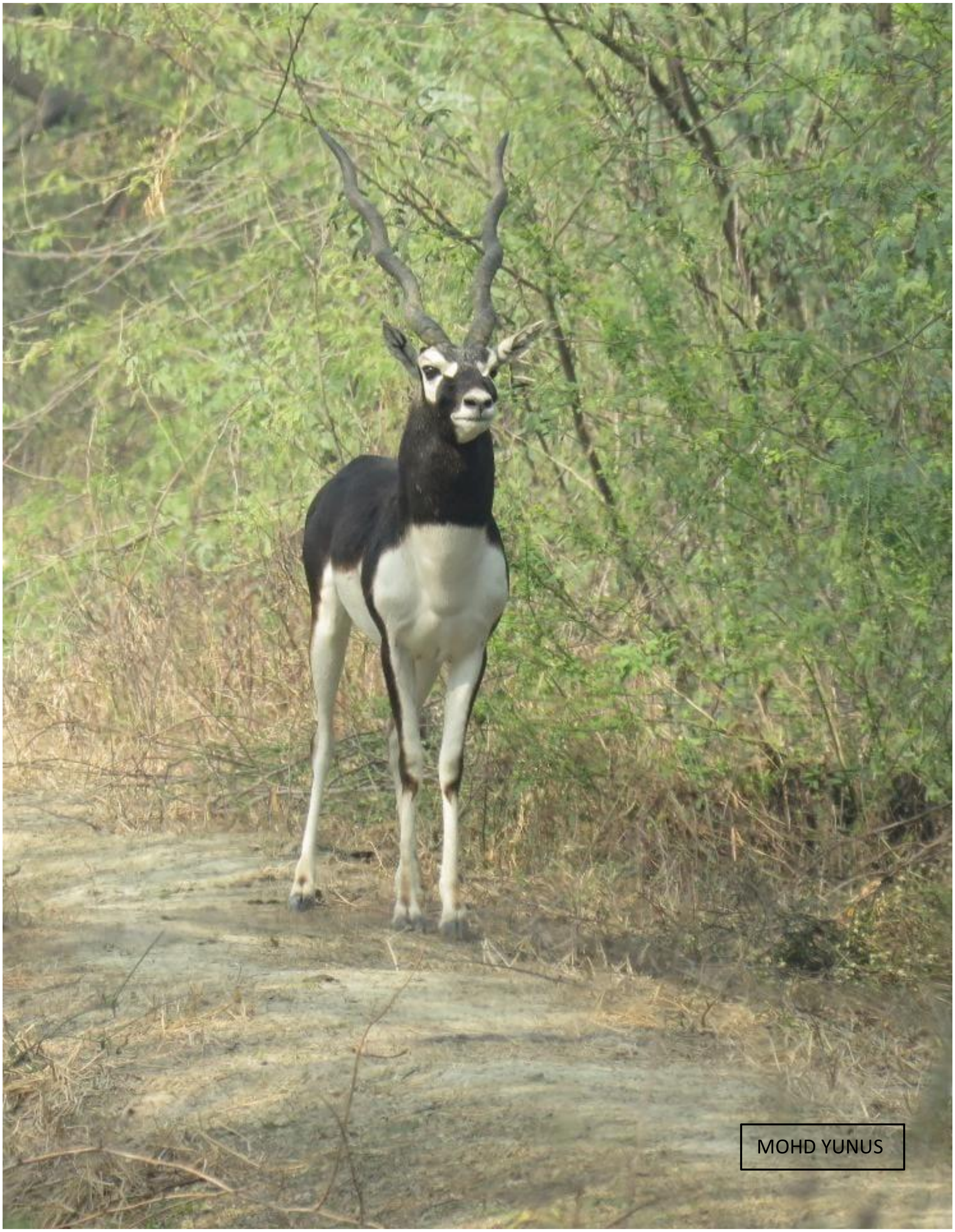


*1 BLACK BUCK (Antilope cervicapra)*

### FEEDING BEHAVIOUR OF BLACK BUCK (*Antilope cervicapra)* DURING WINTER SEASON IN AND AROUND ALIGARH DISTRICT

#### ALIGARH MUSLIM UNIVERSITY

##### DISSERTATION

**MASTER'S OF BIODIVERSITY STUDIES AND MANAGEMENT**

Submitted By

**MOHD YUNUS**

Under the Supervision of

**DR. ORUS ILYAS**

**SESSION: 2023–2025**

**DEPARTMENT OF WILDLIFE SCIENCES**

**FACULTY OF LIFE SCIENCES**

**ALIGARH MUSLIM UNIVERSITY**

### Declaration

I hereby declare that the dissertation entitled "**Feeding Behavior of Black Buck in Winter Around Aligarh"** submitted to **Professor Orus Ilyas**, in partial fulfillment of the requirements for the award of the degree of **Biodiversity and Management**, is a record of original and independent research work carried out by me under the supervision of Professor Orus Ilyas.

This work has not been submitted previously, in whole or in part, to any other university or institution for the award of any degree, diploma, fellowship, or other similar titles or recognition.

All sources of information and assistance have been duly acknowledged. I bear full responsibility for the contents of this dissertation.

Date: **25/05/2025**

Place: **Aligarh**

**Mohd Yunus**

**GP7936**

### Acknowledgement

I would like to express my deepest gratitude to everyone who contributed to this research and made it possible. First and foremost, I extend my sincere thanks to my supervisor, Professor ORUS ILYAS, for their invaluable guidance, continuous support, and encouragement throughout this project. Their expertise and insights have been instrumental in shaping this research. From its inception to its completion, Professor ORUS ILYAS has devoted so much to my study.

From Professor ORUS ILYAS, I learned, not only the knowledge of wildlife but also the scientific approach and the dedicated spirit for work. In addition, he has infused me with enthusiasm for researching and an open mind for different cultures and opinions. Without his supervision, I would not have completed this challenging dissertation.

I am very thankful to Prof. Satish Kumar, Chairman, Department of Wildlife Sciences, Aligarh Muslim University. I am also grateful to him for his guidance, suggestions, and ample facilities to complete the task efficiently.

I express my heartfelt gratitude to my respected teachers Prof. Afifullah Khan, Dr. Nazneen Zehra, Dr. Sharad Kumar, Dr. Kaleem Ahmed, Dr. Ahmad Masood Khan and the last but not the least Professor Jamal Ahmad Khan for their valuable teaching and cooperation.
I would also like to thank to the farmers and local communities of Aligarh for their cooperation and participation in this study. Their willingness to share information and allow access to their farmlands was essential for the fieldwork.

I would like to thank my parents for well, everything. I thank my sister Adeeba and brother Dr. Yusuf for being so supportive from the time when I was in school. I thank all my brothers and sisters for always making me believe that I am a ‘born to be wild’ person.

I would also like to express my heartfelt gratitude to my friend and senior, Mohd Intakhab, for standing by me and supporting me through difficult times. I would like to thank all my friends who have borne my insanity from high school to graduation—Saad-ur-Rehman and Mohd Anas and all the others I have missed.

### Abstract

This study investigates the winter feeding behavior of blackbuck (Antilope cervicapra) across three semi-arid landscapes in Aligarh, Uttar Pradesh—Palla Salu, Rathgaon, and Sikandra Rao. Using vegetation sampling and micro-histological analysis of fecal pellets, we assessed plant availability and dietary composition. A total of 4,125 plant individuals were recorded, with *Prosopis juliflora* and *Cynodon dactylon* emerging as dominant species. However, dietary analysis revealed a clear preference for *Cynodon dactylon, Poa annua*, and *Rumex obtusifolius*, while *Prosopis juliflora,* despite its abundance, was selectively avoided. Bonferroni-adjusted confidence intervals confirmed statistically significant foraging preferences, indicating that blackbuck prioritize nutrient-rich grasses and selectively avoid less palatable or chemically defended plants. These findings emphasize the importance of conserving native forage species and controlling invasive taxa like *P. juliflora* to support blackbuck populations. The study underscores the value of integrating ecological research with habitat management for effective conservation of blackbuck in human-dominated.

### CONTENT

**Chapters** **Page No.**


*1. Introduction*……………………….……………………………..……….1

*1.1 Background of the Study*……………… …..…………………...1

*1.2 Rationale*………………….……………………………...…....4

*1.3 Threats*…………………………………………………………...6

*2. Literature Review*…………………….………………….……..….…8

*3. Study Area*……………………………………………………….......10

*3.1Rathgaon*………………………………………………………11

*3.2 Palla Salu*.......................................................................................12

*3.3 Sikandra Rao*…………………………………….……………….…...13

*4. Methodology*……………………………………………………..……14

*4.1 Vegetation sampling*……………………………………...…...14

*4.2 Study area and habitat diversity*……………………………...….15

*4.3 Sampling design*……………………….…………………….....…15

*4.4 Fecal Sample Collection and Micro-histology*………………......15

*4.5 Laboratory Analysis and Micro-histology*……...……..…………....16

*4.6 Data analysis*………………………...………….……………….16

*5. Result* …………………………………………………………………18

*5.1 Plants Details*.....................................................................................18

*5.2 IVI*……………………………..……………………………...20

*5.3 Pellet Analysis*…………………………………………..….......21

*5.4 Results: Food and Feeding Habits*……………………..……...22

*6. Discussion*…………………………………………………....….24

*7. Conclusion*………………………………………..……………..27

*8. References*…………………………………………...…………29

*9. Tables*…………………………………………..…………..……33

*10. Pictures of Micro-histological slides with plants*…………..………...41

*11. Pictures*…………………………………………………………45

### 1. INTRODUCTION

#### 1.1 Background

The Blackbuck (*Antilope cervicapra*) represents a quintessential symbol of the Indian subcontinent's rich faunal heritage. In the present work, I have examined its distinctive morphological and evolutionary traits, drawing on foundational studies by (Dharmakumarsinhji and Gaekwad 1958), who highlighted the species' appeal to early naturalists and zoologists. As a member of the Antilopinae subfamily within Bovidae, the Blackbuck is a medium-sized, sexually dimorphic antelope that showcases a wide array of ecological and physiological adaptations. The evolutionary backdrop of this species can be traced to the rapid radiation of the bovids approximately 12–15 million years ago, which led to the emergence of the Antilopini tribe (Hassanin & Douzery, 1999), situating the Blackbuck within a complex phylogenetic framework.

One of the most defining features of adult males is their dichromatic pelage, consisting of a dark brown to black dorsal coat in contrast with stark white underparts, legs, buttocks, eye rings, muzzle, and nose. In contrast, females and juveniles display a lighter fawn or tan dorsal coloration with similarly white ventral areas and marked white eye rings. The spiraled horns of males—typically comprising one to four twists—are another key morphological trait. Interestingly, the darker pelage in males is linked to testosterone levels, reflecting the intricate hormonal regulation of secondary sexual traits.

Taxonomically, two subspecies have been identified: *Antilope cervicapra rajputanae* and *A. c. cervicapra*, distinguished on the basis of morphological characteristics such as coat texture, coloration, and horn length (Groves, 1982; Ranjitsinh, 1989). The former, found predominantly in northwestern India, is characterized by its larger size, broader eye rings, spotless white shanks, longer horns, and paler females. In contrast, *A. c. cervicapra*, which occurs in eastern and southern India, tends to be smaller, with finer pelage, shorter horns, and darker males. However, as Ranjitsinh (1989) noted, overlapping ranges and high phenotypic plasticity make the sub specific classification somewhat ambiguous.

Beyond biology, the Blackbuck holds immense cultural and spiritual value in India, revered in ancient mythology and represented in traditional art and religious symbolism (Dharmakumarsinhji & Gaekwad, 1958). Recent molecular phylogenetic research further elucidates its evolutionary history, indicating that Blackbuck diverged from gazelles approximately 1.9–3.4 million years ago (Hassanin & Douzery, 1999), offering deeper insights into the lineage diversification of antelopes in arid and semi-arid landscapes. Blackbuck (*Antilope cervicapra*) are primarily grazers, and their diet reflects strong seasonal patterns (Prasad, 1981; Ghosh et al., 1984; Ranjitsinh, 1989; Jhala, 1997). During the monsoon, they concentrate in grasslands where they feed mainly on tender grass shoots. When available, they also consume fresh leaves from browse species that sprout with the rains. As the season transitions into winter and the protein content in grasses declines, blackbuck modify their foraging strategy. In landscapes like Mudmal (Andhra Pradesh) and Velavadar (Gujarat), they frequently move into agricultural fields, feeding on crops such as wheat, barley, and gram (Ranjitsinh, 1989; Jhala, 1993a). Their dependence on crop fields tends to increase until the harvest season. In addition to crops, blackbuck also consume seed pods of species like *Prosopis cineraria* and the invasive *P. juliflora*, which can make up a substantial portion of their diet—up to 41% of dry weight in some winters (Jhala, 1991).

During the summer, when grass quality is at its poorest and fields are often fallow, browse becomes more important in their diet (Ranjitsinh, 1989; Jhala, 1991). In some areas, such as Guru Bishnoiyan—a Bishnoi village—blackbuck heavily feed on *P. cineraria* pods during this period (Goyal et al., 1988). In Mudmal, they are also known to forage on groundnut crops and *Acacia* pods (Prasad, 1981). Although browse may be consumed in smaller amounts compared to grasses, it provides essential nutrients due to its higher energy and protein content. Remarkably, even in the absence of browse, blackbuck can survive on low-quality forage such as coarse grasses and dry plant matter (Jhala, 1991).

Habitat use in blackbuck is influenced not only by forage availability but also by the risk of predation. They generally avoid areas with dense or tall vegetation where predators may hide (Ranjitsinh, 1989; Jhala, 1991). Instead, they prefer open landscapes and tend to rest in exposed areas even during peak daytime heat. Studies have shown that they exhibit heightened vigilance near tall grass or shrub cover, likely due to increased perceived threat from predators (Isvaran, 2007). Similar to other antelopes that have evolved in open habitats, blackbuck rely heavily on early detection of predators and swift flight to escape danger (Mungall, 1978; Ranjitsinh, 1989). Consequently, their preference for open short grasslands seems to be shaped by a combination of dietary needs and anti-predator strategies.

#### Coping Strategies of Blackbuck in Arid Environments

In the arid and semi-arid regions where blackbuck naturally occur, they often face tough conditions, especially during the summer. Water becomes scarce, and the nutritional quality of available forage drops significantly. According to Jhala (1997), during the peak of summer, the digestibility of dry matter in their diet can fall to just 32%, and the crude protein content may go as low as 3.6%. This led me to think about how blackbuck manage to survive during these difficult months.

One way they seem to cope is by minimizing their overall energy expenditure. Since the energy they gain from forage is reduced, and the cost of digestion increases—especially with coarse, dry forage—it makes sense that they would try to reduce their food intake. Jhala (1991) even observed that blackbuck might lose more protein in their feces than they gain from food, which probably means they're breaking down body tissues to meet their needs. Cutting down on food intake would reduce digestive stress, and possibly foraging movements too, helping conserve more energy overall.

These assumptions are backed by both physiological and behavioral studies. For example, a study on captive blackbuck showed a noticeable drop in food intake from the monsoon season to summer (Jhala, 1997). Similarly, in the wild, blackbuck were found to forage less during March–April compared to the cooler months of November–December (Isvaran & Jhala, 2000), which supports the idea that they are adapting their foraging activity based on energy availability and environmental stress.

Water availability is another major challenge, but blackbuck seem well adapted to this too. In the absence of water sources, they mainly rely on preformed water in the plants they eat (Jhala et al., 1992). During the monsoon, when plant moisture can be as high as 70%, they can go without drinking for a couple of days. But when summer arrives and forage moisture drops to about 20%, they are usually seen drinking once a day. Another interesting behavior I came across is their tendency to feed intensively in the early morning hours. This likely helps them take advantage of dew, since plant moisture is higher when humidity is high in the morning (Jhala et al., 1992).

Overall, blackbuck show a combination of physiological and behavioral adaptations that allow them to cope with harsh summer conditions in their natural habitats.

#### 1.2 Rationale

The blackbuck (*Antilope cervicapra*) is primarily a grazing ungulate, as established by multiple studies (Prasad, 1981; Ghosh et al., 1984; Ranjitsinh, 1989; Jhala, 1997). During the monsoon season, their diet is dominated by fresh grass shoots, and herds commonly aggregate in open grasslands. The sprouting of new leaves on browse species—stimulated by seasonal rains—supplements their nutritional intake in this period. However, as the protein content of grasses declines during the winter months, blackbuck gradually shift toward alternative food sources. In regions like Mudmal (Andhra Pradesh) and Velavadar (Gujarat), this includes agricultural crops such as wheat, barley, and gram, with crop field usage peaking in late winter just prior to harvest (Ranjitsinh, 1989; Jhala, 1993a). Additionally, seed pods from native and exotic leguminous species, particularly *Prosopis cineraria* and *P. juliflora*, become vital dietary components, comprising up to 41% of their dry weight intake during certain winters (Jhala, 1991).

In the summer, as grass quality further deteriorates and agricultural fields lie fallow, blackbuck diets become increasingly reliant on browse species (Ranjitsinh, 1989; Jhala, 1991). For instance, in the Bishnoi village of Guru Bishnoiyan, *P. cineraria* seed pods constitute a major food source during this period (Goyal et al., 1988). In Mudmal, individuals have been observed feeding on groundnut crops and pods from *Acacia* species (Prasad, 1981). While browse comprises a smaller proportion of their annual diet compared to grasses, it remains a crucial resource due to its higher protein and energy content. Nevertheless, blackbuck have shown remarkable adaptability, often subsisting on coarse grasses and plant litter when higher-quality forage is unavailable (Jhala, 1991).

Predation risk is another key factor influencing blackbuck foraging and habitat preferences. Several studies indicate an aversion to tall grasslands and dense shrublands, likely due to reduced visibility and increased vulnerability to predators (Ranjitsinh, 1989; Jhala, 1991). Increased vigilance in proximity to tall vegetation supports this hypothesis, suggesting that blackbuck assess predator threat levels based on habitat structure (Isvaran, 2007). Like many antelope species inhabiting open plains, blackbuck rely heavily on early predator detection and rapid flight responses, a behavior well-documented across their range (Mungall, 1978; Ranjitsinh, 1989). This preference for open landscapes persists even during the heat of the day when individuals rest in exposed areas, possibly prioritizing visibility over thermoregulation.

As one of the last remaining wild antelope species in India, the blackbuck plays a critical ecological role within grassland ecosystems. It also holds immense cultural and spiritual value in the Indian subcontinent. Although currently listed as “Least Concern” by the International Union for Conservation of Nature (IUCN), blackbuck populations are increasingly threatened by habitat fragmentation, expanding agriculture, poaching, and competition with livestock. Given their ecological sensitivity, blackbuck serve as effective bioindicators for grassland health. Understanding seasonal dietary shifts, particularly during nutritionally challenging periods like winter, is therefore essential for informing conservation policies and land management strategies aimed at ensuring the long-term viability of blackbuck in the wild

#### 1.3 Threats

The dramatic decline of blackbuck (*Antilope cervicapra*) populations across India is primarily attributed to habitat loss and fragmentation (Rahmani, 2001; Sankar & Acharya, 2004). Blackbuck are dependent on semi-arid grasslands and open scrub habitats, which have become increasingly degraded and subdivided due to expanding human settlements, intensive agriculture, and livestock grazing pressures (Kumar et al., 2019). These habitats, once widespread, are now largely restricted to small protected areas such as Rehekuri, Tal Chhapar, and Velavadar, along with fragmented patches embedded in agricultural landscapes and rural settlements (Jhala, 1991). Overgrazing by high densities of livestock not only depletes forage but also exacerbates competition, while instances of crop damage by blackbuck often strain human-wildlife relationships, undermining conservation initiatives (Sharma & Singh, 2017).

Hunting has also played a significant role in reducing blackbuck densities (Ranjitsinh, 1989). Nevertheless, in regions where local communities—such as the Bishnois—uphold religious or cultural taboos against harming wildlife, blackbuck populations tend to persist in better numbers (Dharmakumarsinhji & Gaekwad, 1958). A growing ecological concern is the aggressive invasion of grasslands by exotic species such as *Prosopis juliflora* and *Opuntia dillenii*, which promote woody vegetation encroachment and degrade open habitats essential for blackbuck survival (Mishra et al., 2020). Invasive plant infestations, particularly around regions like the Mudumalai Wildlife Sanctuary, highlight the urgency of weed eradication programs and habitat restoration efforts (Kumar et al., 2019).

Effective conservation strategies must prioritize sustaining blackbuck populations in multi-use, human-dominated landscapes (Jhala, 1991). This involves preserving contiguous networks of grassland habitats and promoting traditional land uses such as pastoralism, which are more compatible with grassland conservation than intensive agriculture or industrial development (Kumar et al., 2019). In addition to managing invasive species, reducing hunting pressure and mitigating crop damage conflicts through community engagement and compensation schemes are crucial (Mishra et al., 2020). Another important yet often overlooked threat is predation by free-ranging dogs, which must be addressed through local control measures, especially in blackbuck strongholds (Sankar & Acharya, 2004).

Historically, blackbuck have also suffered from unjustified persecution. During the colonial period, exaggerated claims of crop damage led to widespread hunting, with the species often classified as vermin (Ranjitsinh, 1989). Nomadic tribes, including the Dafers, were known to hunt blackbuck for commercial purposes (Dharmakumarsinhji & Gaekwad, 1958). Even after India's independence in 1947, crop raiding was frequently used as a rationale to issue weapons under the guise of crop protection, further intensifying pressure on blackbuck populations (Ranjitsinh, 1989). Nonetheless, traditional reverence among communities such as the Bishnois continues to offer a buffer against local extinctions, underscoring the importance of culturally rooted conservation ethics (Dharmakumarsinhji & Gaekwad, 1958).

Protected areas like Velavadar have successfully maintained relatively large populations of blackbuck. However, ongoing challenges such as livestock encroachment, periodic droughts, and limited water availability continue to affect their long-term survival (Sankar & Acharya, 2004). Although disease outbreaks are rare, past epidemics like the rinderpest outbreak in Orissa during the 1940s have demonstrated the potential vulnerability of isolated populations (Ranjitsinh, 1989). Combined with illegal hunting and trapping, these pressures have historically contributed to significant population declines (Ranjitsinh, 1989).

### 2. LITRATURE REVIEW

The blackbuck (*Antilope cervicapra*) is a prominent herbivore species native to the Indian subcontinent, and I chose to study it due to its ecological relevance and adaptability across seasons (Rahmani, 2001; Sankar & Acharya, 2004). Its survival is closely linked to the availability and nutritional quality of forage, which varies with seasonal changes and habitat conditions (Isvaran, 2005; Ranjitsinh, 1989). In winter, when natural vegetation becomes scarce and less nutritious, understanding their dietary habits becomes crucial, especially in semi-arid landscapes like Aligarh (Khan et al., 1996; Geeta & Acharya, 2012). Blackbuck mainly feed on graminoids, but they also consume forbs and shrubs based on local availability (Mishra et al., 2020). During winter, they tend to rely more on dry grasses and browse, which reflects their adaptive strategies for coping with seasonal food shortages. Studies from northern and central India show that blackbuck often move into nearby agricultural fields during lean periods, which increases the risk of conflict with farmers (Sharma & Singh, 2017). The Aligarh region, with its mix of semi-arid conditions and farmland, presents an important context for studying these interactions. Their winter diet often includes crop residues, legumes, and dry grasses, which not only help meet their nutritional needs but also raise concerns about crop damage and habitat degradation. Techniques like fecal analysis and direct observation have been used to study their diet, with fecal analysis being especially useful due to its non-invasive nature and accuracy (Prasad et al., 2018). Comparative studies have shown that habitat management significantly influences forage availability, but most research so far has focused on large protected areas, leaving a gap in our understanding of blackbuck living near human settlements like Aligarh. I believe that studying their winter diet in this region can offer valuable insights for conservation planning, habitat management, and reducing human-wildlife conflict, ultimately promoting better coexistence between blackbuck and agricultural communities.

#### Research questions:

- What plant species are present in the winter diet of blackbuck across the three selected study sites, as determined by micro-histological analysis of faecal pellets?
- What is the Important Value Index (IVI) of forage plant species available in all sites, and how does it reflect the dominance and availability of forage resources?
- "Which plant species are preferred by blackbuck in Palla Salu, Sikandra Rao, and Rathgaon during winter, based on a comparison between their dietary occurrence and field availability (IVI)?"

#### Research Objective:

- To investigate the vegetation comparison in Palla salu, Rathgaon and Sikandra Rao.
- To investigate in winter food and feeding habit of black buck in selected areas.
- To investigate the micro-histological features of plant availability in study areas.
- To prepare plant and plant slides.

### 3. STUDY AREA

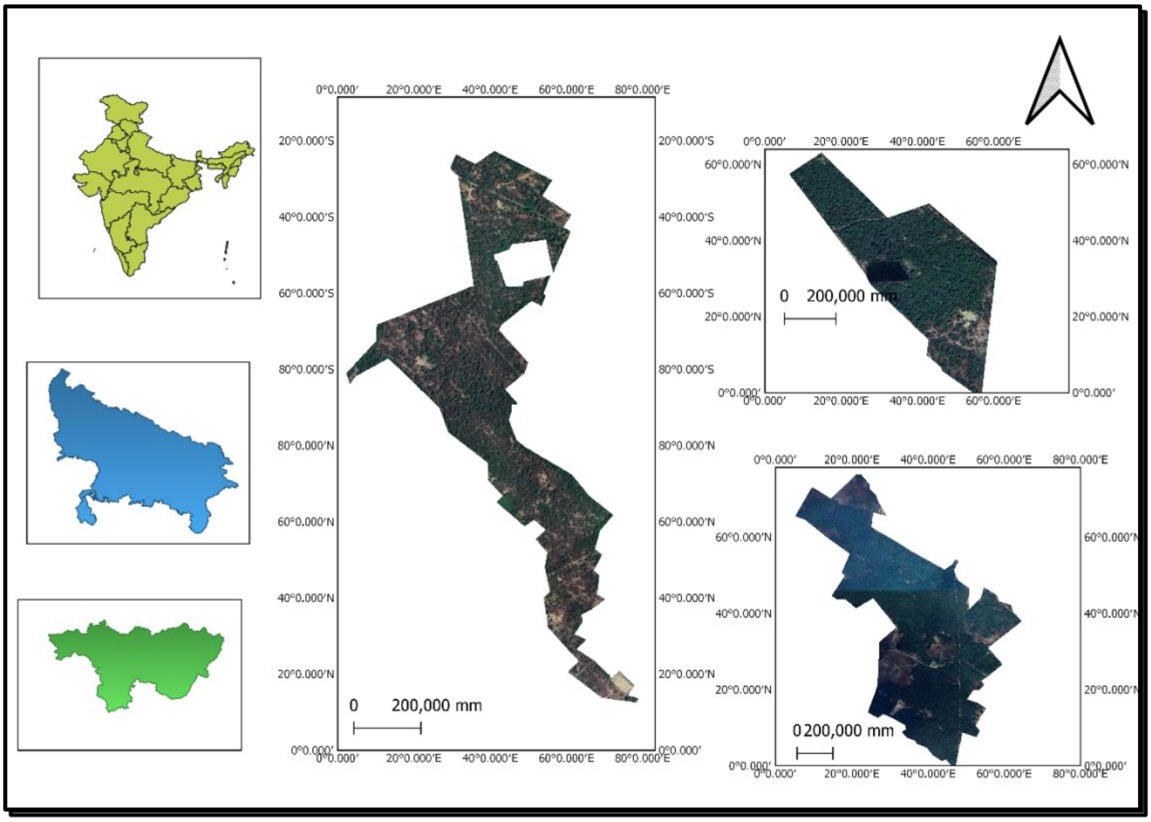


***Figure2 Map of Study Area***

Aligarh, situated in western Uttar Pradesh, falls on the southern margin of the Upper Ganga–Yamuna Doab, approximately 100 km southeast of Delhi and about 40 km southwest of the Ganges River (Encyclopedia Britannica). The city stands at an elevation of around 178 meters above sea level and covers an area of 5,019 square kilometers, stretching about 112 km east to west and 72 km north to south. The climate is classified as hot semi-arid (Köppen BSh), characterized by intense summer heat beginning in April, with peak temperatures in May. The monsoon sets in by late June and lasts till early October, contributing the majority of the annual rainfall, which averages around 800 mm.Winter are generally mild, with temperatures ranging from 2°C to 22°C. The soil in this region is primarily light-textured loam with patches of sandy soil, influenced by loess deposits blown from the Thar Desert in Rajasthan. This geographical and climatic context supports diverse agricultural practices, making Aligarh a significant center for agricultural production and trade.

The study was conducted in Rathgaon, Pallasalu, and Sikandra Rao, located in the Aligarh district of Uttar Pradesh, India. These regions are characterized by semi-arid climatic conditions, interspersed with agricultural landscapes, fallow lands, and patches of natural vegetation.

#### 3.1 Rathgaon

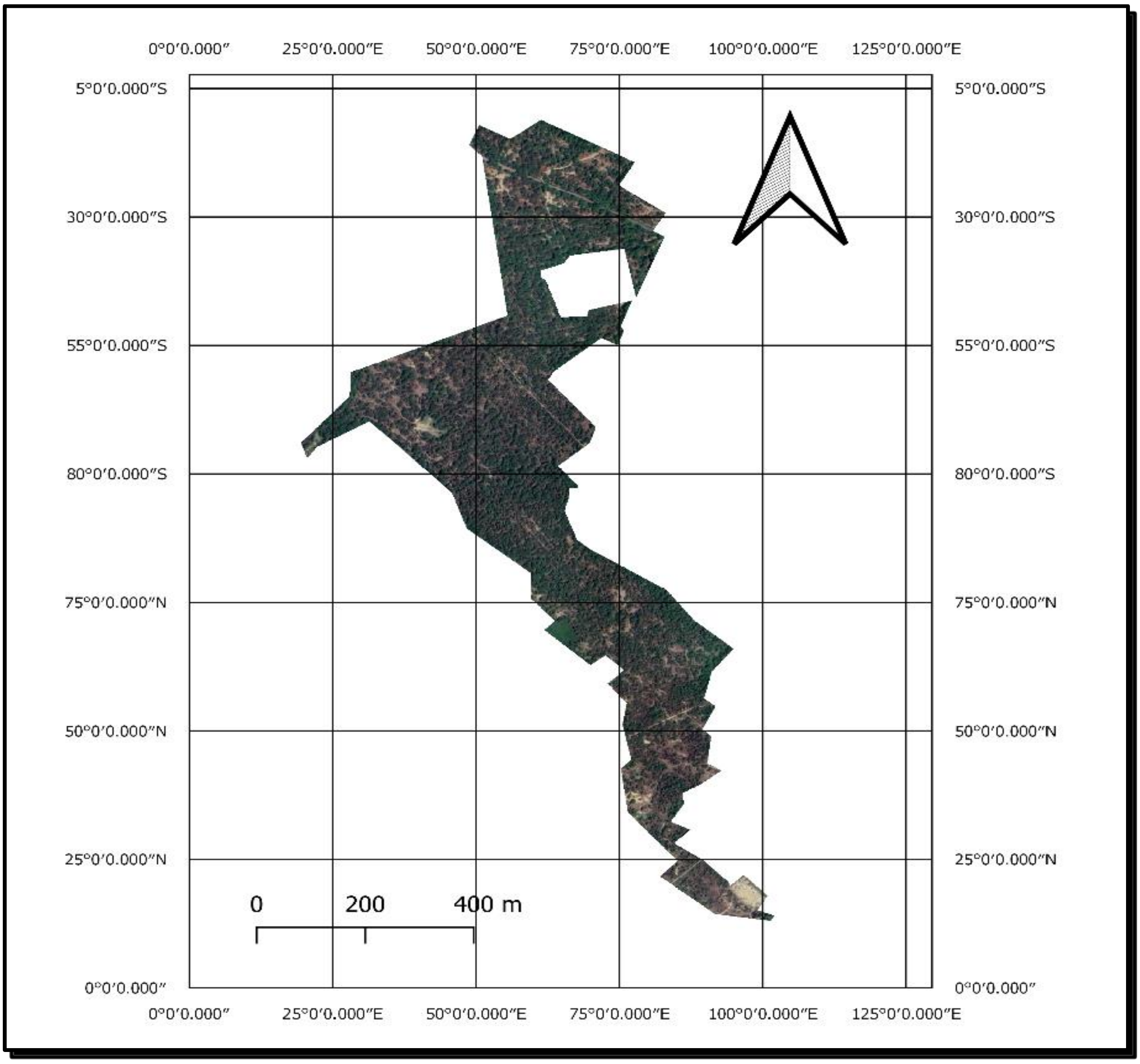


*Figure 3 Rathgaon*

Rathgaon is a small region in the Aligarh district of Uttar Pradesh with around 0.4 km² of forest area. The climate here is humid subtropical (Cwa), which means the summers are hot, monsoon is strong, and winters are cool (Kottek et al., 2006). The area receives about 1,000 to 1,200 mm of rainfall every year, mostly during the southwest monsoon from June to September (Singh et al., 2014). The land is mostly alluvial plains, making the soil fertile and supporting a variety of plant life (Krishnan, 1982). This mix of weather and land makes Rathgaon an interesting place for ecological research.

#### 3.2 Palla Salu

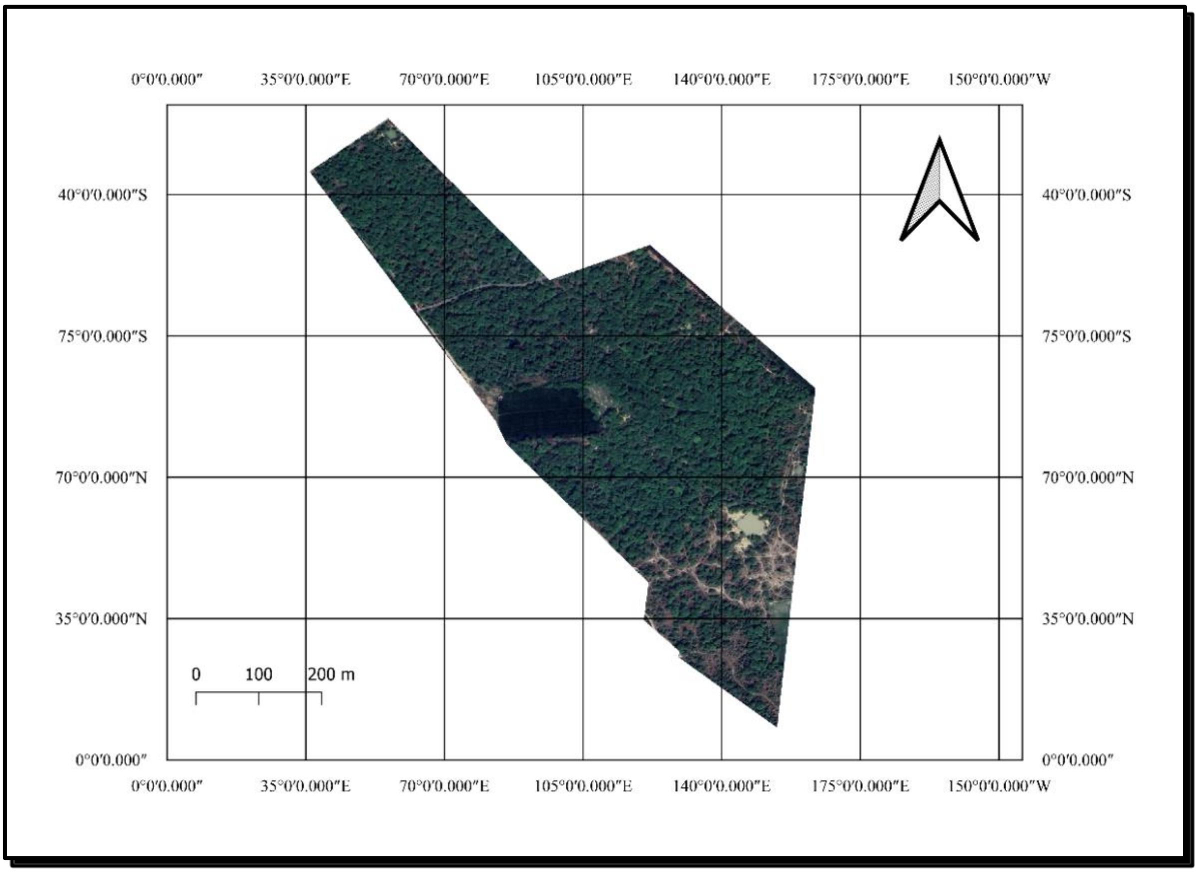


*Figure 4 Map of Palla Salu*

Palla Salu is also located in the Aligarh district and has a similar humid subtropical (Cwa) climate with very hot summers and relatively cooler winters (Kottek et al., 2006). It gets around 800–900 mm of rainfall every year, especially during the southwest monsoon from mid-June to September (Singh et al., 2014). In May and June, temperatures often go beyond 40°C, while in January, they drop to about 10°C (Yadav & Dubey, 2018). These weather patterns affect how plants grow and how animals like blackbuck use the area.

#### 3.3 Sikandra Rao

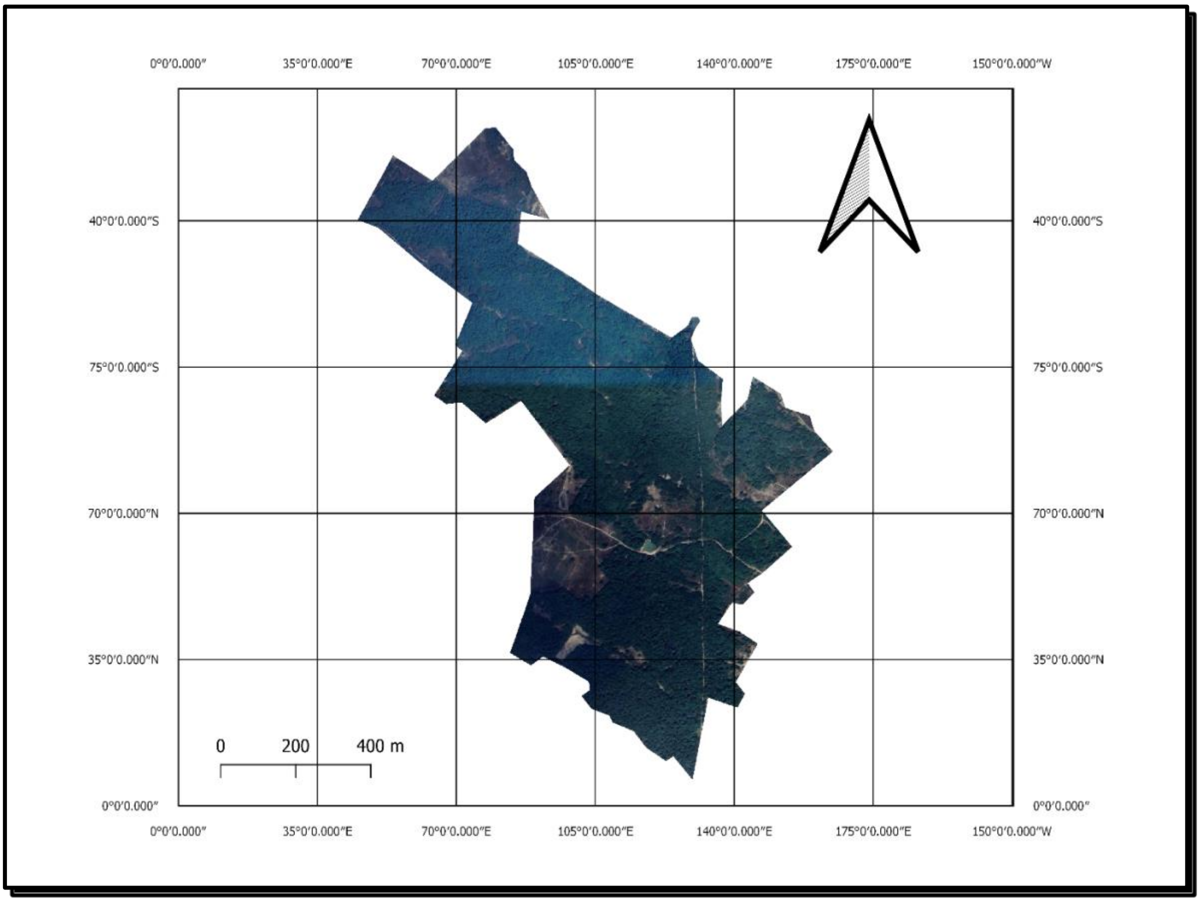


*Figure 5 Map of Sikandra Rao*

Sikandra Rao is in Hathras district, near Aligarh, and has about 0.9 km² of forest area. The climate is also humid subtropical (Cwa), with scorching summers and mild winters (Kottek et al., 2006). In May and June, the temperature crosses 40°C, while in winter it drops to around 10°C (Singh et al., 2014; Yadav & Dubey, 2018). The region gets about 1,000 mm of rain every year, mostly during the monsoon season from June to September. This rainfall and the fertile soil here support diverse plant species, which is important for herbivores like the blackbuck (Gupta & Kumar, 2019).

### 4. METHODOLOGY

#### 4.1Vegetation Sampling

To investigate winter foraging patterns of blackbuck (Antilope cervicapra), a systematic vegetation sampling approach was employed across three key winter habitats within the semi-arid landscape of Aligarh district: Palla Salu (0.33 $km^2$), Sikandra Rao (0.90 $km^2$), and Rathgaon (0.40 $km^2$). These sites were selected based on prior observational data indicating regular blackbuck presence during winter (Jhala,1991).

Sampling was conducted monthly from January to March 2025, on fixed dates (15th, 16th, and 17th of each month) to maintain temporal consistency . A stratified line transect design was used, with three linear 200-meter transects established at each site (Kumar & Shahabuddin, 2005). Along each transect, vegetation was sampled using an alternating lateral quadrat placement pattern—at every 20-meter interval, quadrats were placed 10 meters perpendicular to the transect axis, alternating between the left and rightsides (Mishra,1968).

Quadrat dimensions were standardized by vegetation type following established ecological protocols (Kershaw, 1973). Specifically, 1 m × 1 m quadrats were used for sampling herbaceous plants and graminoids, 5 m × 5 m quadrats for shrubs, and 10 m × 10 m quadrats for trees. In each quadrat, floristic inventories were carried out, and species presence, abundance, and phenological status were recorded.

To aid accurate taxonomic identification, the PlantNet mobile application was used, which employs AI-based image recognition to assist with plant species identification in the field (PlantNet, n.d.). Using these data, standard phytosociological parameters—frequency, density, and relative abundance—were computed based on established methodologies (Mishra, 1968).

To determine the preference or avoidance of plant species by blackbuck, Bonferroni confidence intervals were used to compare the proportion of each plant species in the diet with its availability in the environment. This approach provided a statistically robust assessment of blackbuck foraging behavior and habitat use during winter (Byers et al., 1984).

#### 4.2 Study Area and Habitat Diversity

The study focused on three distinct locations—Palla Salu, Rathgaon, and Sikandra Rao—representing a gradient of natural and human-modified habitats. This section was based on habitat selection theory, which suggests that herbivores choose habitats based on resource availability, safety, and ecological pressures (Manly et al., 2002). By studying these areas, the research aimed to understand how land-use changes affected dietary resources for blackbuck, a species that adapts well to grassland and agricultural landscapes (Rahmani, 2001).

#### 4.3 Sampling Design

A random sampling strategy was used to collect fecal pellets from different habitat types, such as grasslands, agricultural fields, and mixed-use landscapes. Random sampling was chosen to minimize bias and ensure a representative dataset (Levy & Lemeshow, 2013). Transects of 500–1000 meters were set up to cover various microhabitats and capture spatial variations in plant availability. Approximately 15 samples per site were collected to ensure statistical reliability in ecological studies (Gotelli & Ellison, 2013). The total were 45 samples for all of 3 different sites.

#### 4.4 Fecal Sample Collection and Handling

Fecal analysis was used as a non-invasive method to study dietary habits (Putman, 1984). Fresh fecal pellets were identified based on moisture content, collected using sterilized tools, and labeled with location, date, and habitat type. The samples were stored in airtight containers and transported in insulated boxes to prevent degradation (Holechek et al., 1982). This method minimized disturbance to animal populations and followed ethical guidelines for wildlife research (Sutherland, 2006).

#### 4.5 Laboratory Analysis and Micro-histology

Micro-histological analysis was employed as an essential technique to examine herbivore diets by identifying plant fragments in feces or pellets (Stewart, 1967; Sparks & Maleckas, 2004). Reference slides were prepared by crushing plant material into small pieces resembling chewed fragments (Baumgardt et al., 1964). The material was treated with concentrated nitric acid ($HNO_3$) and distilled water (1:3 ratio) to break down cell walls and make the material transparent (Holechek et al., 1982). It was then dehydrated through a series of alcohol and water washes, finishing with absolute alcohol to remove moisture (Baumgardt et al., 1964). Finally, the dehydrated material was mounted on slides using glycerine or Canada balsam for detailed microscopic examination (Storr, 1961). These slides helped in identifying plant species in the blackbuck's diet, contributing to a better understanding of their foraging behavior and ecological interactions.

#### 4.7 Data Analysis

In the study, carpological analysis was employed to examine plant remains, specifically seeds, fruits, and other plant parts, found in the feces of the blackbuck (*Antilope cervicapra*). This method was used to identify and quantify the plant species consumed by the blackbuck. Carpological analysis proved crucial for understanding dietary preferences and foraging behavior, particularly in the winter months when direct observation of feeding behavior was difficult (Sparks & Maleckas, 2004; Stewart, 1967).

Fecal samples from various habitats across the study areas were analyzed under a microscope to identify plant fragments such as seeds and fruits. These identifications allowed for the determination of species composition in the blackbuck's diet (Stewart, 1967). The results were used to quantify the proportional contributions of different plant species to the diet, enhancing understanding of the herbivore’s interaction with its environment (Holechek et al., 1982; Sparks & Maleckas, 2004). This technique was valuable in determining dietary selectivity and assessing which plant species were preferred during the winter season.

Moreover, carpological analysis was combined with other methods—vegetation sampling and micro-histology—to cross-validate findings and gain a comprehensive understanding of blackbuck foraging preferences (Holechek et al., 1982). This multi-method approach provided insights into how blackbucks responded to the availability of different plant species and how their diet changed in response to environmental factors (Putman, 1984). Therefore, carpological analysis served as an essential tool to explore dietary composition and habitat use by blackbucks during the winter months.

### 5. RESULT

#### 5.1 Plants Details

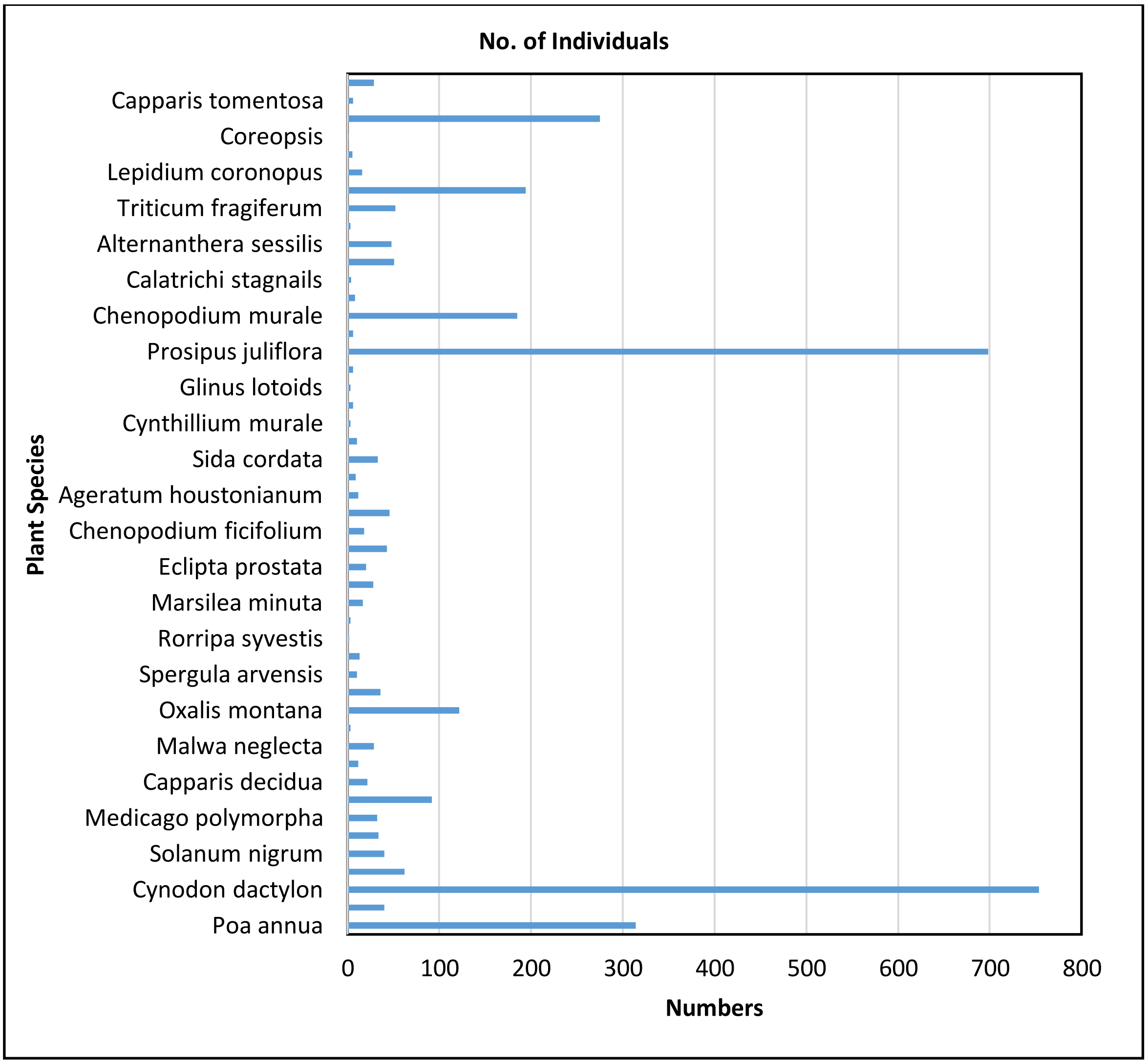


*Figure 6 Plants Species List*

The vegetation survey conducted in the study area revealed a diverse range of plant species, with varying numbers of individuals. According to the results, the most abundant species were Cynodon dactylon (754 individuals), Prosipus juliflora (698 individuals), and Poa annua (314 individuals). These species are likely to be well adapted to the local environmental conditions and may play a crucial role in the ecosystem (Kershaw, 1973).

The abundance of these species can be attributed to their ability to thrive in a variety of habitats and their tolerance to different environmental factors such as temperature, moisture, and soil type (Mishra, 1968). For example, Cynodon dactylon is a highly adaptable species that can grow in a range of environments, from tropical to temperate regions (Skerman & Riveros, 1990).

In contrast, some species had very low numbers, such as Rorripa syvestis (2 individuals), Atriplex patula (3 individuals), Dhatura innoxia (3 individuals), and Carduus teniflorous (3 individuals). These species may be rare or endangered in the study area, and their low abundance could be due to various factors such as habitat degradation, competition with other species, or environmental stress (Jain & Singh, 1985).

Understanding the abundance and distribution of plant species is essential for informing conservation and management strategies for the ecosystem (Kent & Coker, 1992). For example, species with low abundance may require special conservation efforts, such as habitat restoration or protection from invasive species (Given, 1994).

#### 5.2 IVI

Vegetation sampling across the blackbuck habitats in Palla Salu, Rathgaon, and Sikandra Rao revealed the presence of 4125 individual plants belonging to various herb and shrub species. Among all, *Prosopis juliflora* (Mesquite) showed the highest Importance Value Index (IVI) of 47.81, making it the most dominant species in the study area. This was followed by *Cynodon dactylon* (Bermuda grass) with an IVI of 45.84, and *Poa annua* (Annual Meadowgrass) with 14.62. These species were found in most of the quadrats and also had high numbers, showing that they are well adapted to the environment of the study sites.

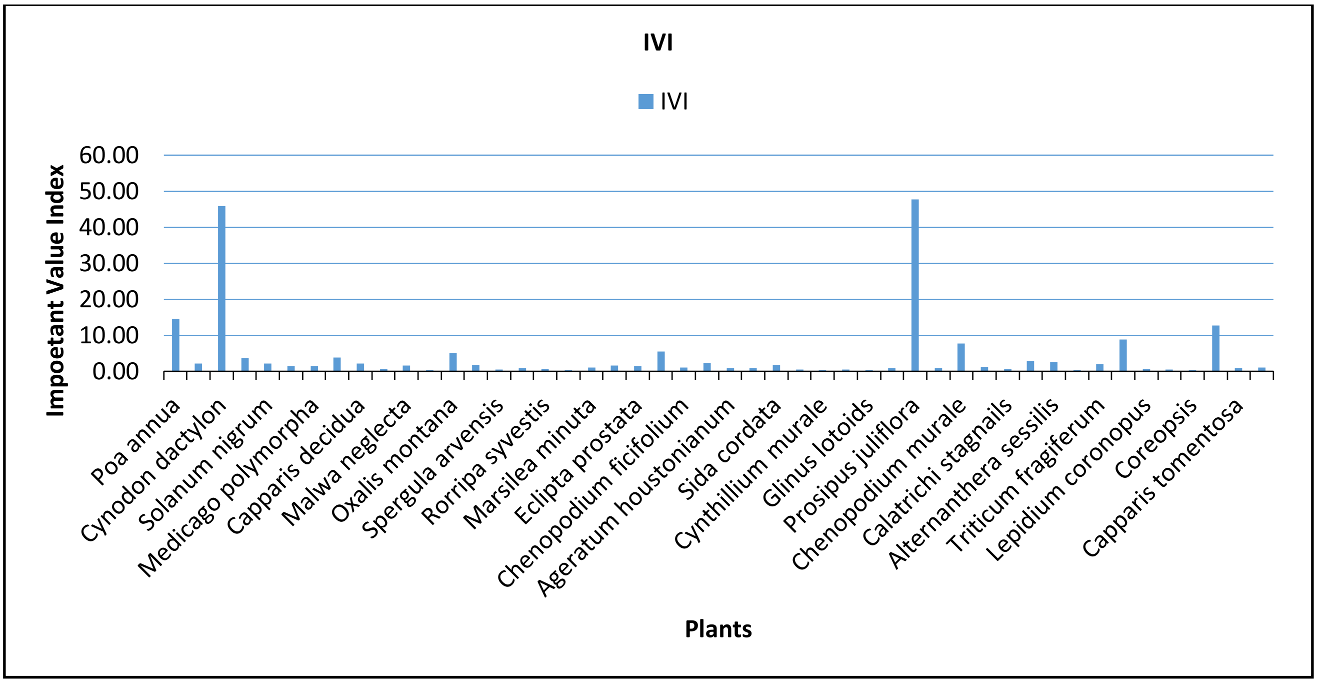


***Figure 7 IVI of Plants***

*Prosopis juliflora* is an invasive species commonly found in dry areas, and it often spreads quickly, reducing space for other native plants (Pasiecznik et al., 2001). On the other hand,

*Cynodon dactylon* is a valuable forage grass known to be eaten by blackbucks and other herbivores (Sankaran, 2005; Isvaran, 2007). Its high IVI suggests that it could be an important part of the blackbuck's winter diet. Other species like *Calamagrostis epigejos*, *Chenopodium vulvaria,* and *Chenopodium murale had* moderate IVI values, showing their regular presence. Many species such as *Coreopsis, Rumex obtusifolius,* and *Cynthillium murale* had very low IVI values, which means they were rare or seen only in a few places.

The overall plant community was uneven. A few species were found everywhere and in large numbers, while many were present in small numbers or only in certain spots. This is common in open grassland areas, especially those used for grazing or disturbed by human activities (Tripathi et al., 2016; Mishra et al., 2002).The dominance of forage grasses like *Cynodon dactylon* indicates that the area still supports palatable species, which could attract blackbuck. However, the spread of invasive plants like *Prosopis juliflora* may affect the availability of native grasses and herbs in the long run, possibly changing the feeding habits of blackbuck over time (Rai & Tripathi, 2018).

#### 5.3 Pallet Analysis

The dietary analysis of blackbuck (*Antilope cervicapra*) using Bonferroni-adjusted confidence intervals revealed distinct patterns of forage selectivity during the winter season across the study sites in Aligarh. Among the 18 identified plant species, *Cynodon dactylon* (usage: 34.89%) and *Poa annua* (10.41%) were significantly preferred over their field availability (27.7% and 7.75% respectively), suggesting their critical role as high-quality forage grasses in winter diets (Jhala, 1997; Sankaran, 2005). *Rumex pulcher* and *Atriplex patula* were also consumed disproportionately more than their availability, indicating selective intake possibly due to higher palatability or essential micronutrients (Goyal et al., 1988; Robbins, 1993). In contrast, species such as *Prosopis juliflora*, despite being ecologically dominant (25.65% availability), were markedly avoided (17.28% usage). This avoidance may be attributed to its known chemical defenses, spiny morphology, or lower digestibility (Pasiecznik et al., 2001; Rai & Tripathi, 2018). Similarly, *Solanum nigrum*, *Lactuca floridana*, and *Parthenium hysterophorus* were all underutilized despite their presence in the environment, likely due to their content of toxic alkaloids or unpalatable secondary metabolites (Robbins, 1993). Several other species, including *Chenopodium vulvaria*, *Oxalis acetosella*, and *Calamagrostis epigejos*, were used in proportion to their availability, thus falling into the neutral category, indicating their role as opportunistic or fallback forage during resource-limited periods (Isvaran, 2007). These findings emphasize the blackbuck's ability to forage selectively even under winter stress, maximizing nutrient intake while minimizing ingestion of suboptimal plant material—a strategy crucial for survival in semi-arid, human-dominated landscapes.

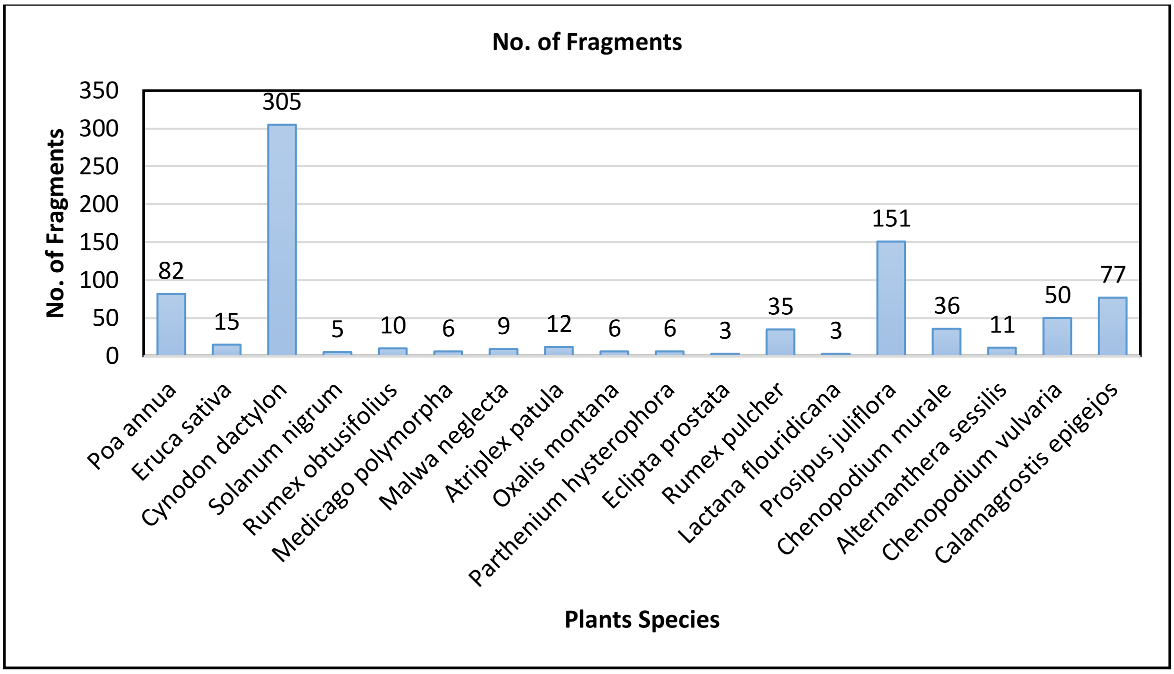


***Figure 8 Plants Fragments in Micro-histology***

The presence of these plant species in the pellets indicates that they are an important part of the blackbuck's diet. The dominance of certain plant species, such as Bermuda Grass and Mesquite, may indicate a preference for these species by blackbuck. Understanding the dietary preferences of blackbuck can inform habitat management and conservation efforts, ensuring that the habitat provides a sufficient supply of preferred plant species (Rodgers & Panwar, 1988; Kumar & Shahabuddin, 2005).

This study highlights the importance of micro-histological analysis in understanding the dietary habits of wildlife. By analyzing the plant fragments in blackbuck pellets, researchers can gain insights into the feeding behavior and habitat requirements of this species. This information can be used to develop effective conservation strategies that protect the habitat and ensure the long-term sustainability of blackbuck populations (Groves, 1980; Champion & Seth, 1968).

#### 5.4 Results: Food and Feeding Habits

**Pie-Zα/2k Pie(1-Pie) / n ≤ Pie ≤ Pie + Zα/2k Pie (1-Pie) / n.**

The analysis of blackbuck dietary selectivity during winter revealed clear patterns of forage preference and avoidance. *Cynodon dactylon* (Bermuda Grass) was the most prominent preferred species, with a usage proportion of 34.9% significantly higher than its availability at 27.7%. Other preferred species included *Poa annua* (Annual Meadowgrass), with a usage proportion of 10.4% compared to its availability of 7.5%, and *Rumex pulcher* (Fiddle Dock), with a usage proportion of 4.0% versus its availability of 1.6%. These species were consumed disproportionately more than expected, underscoring their critical nutritional

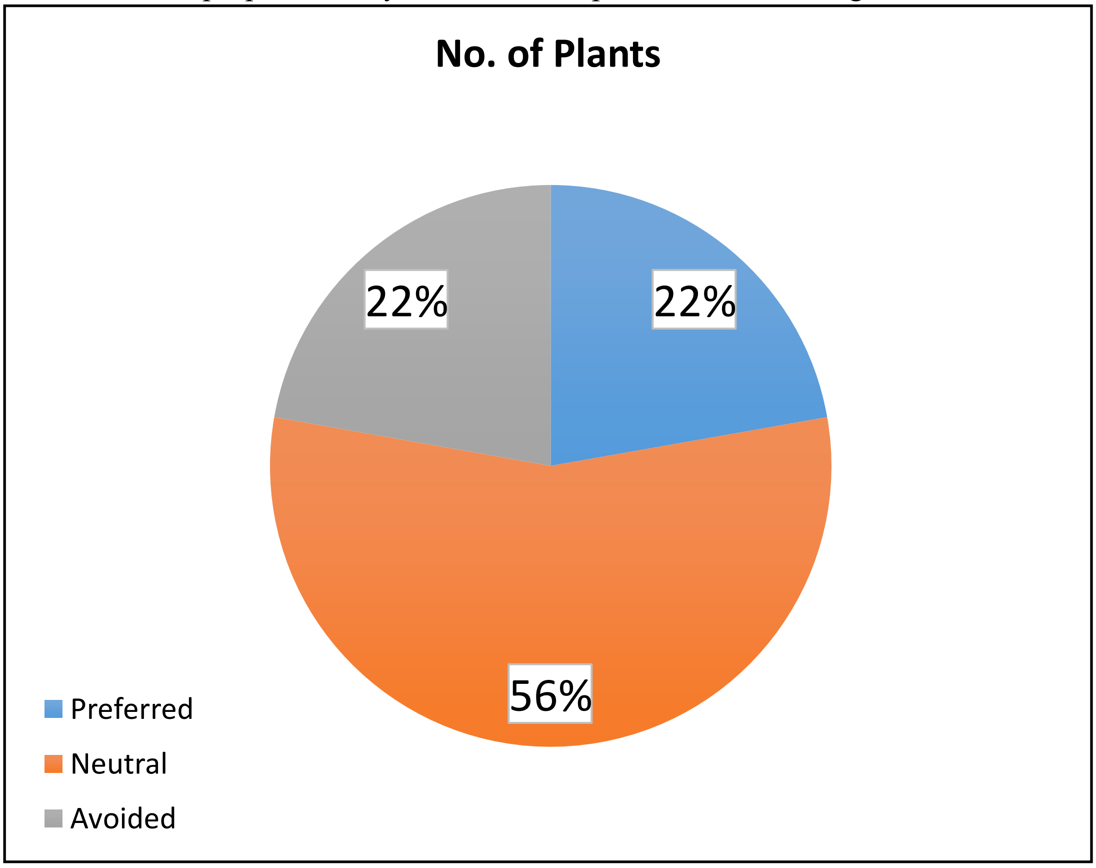


***Figure 9 Preference pie chart***

value

during resource-scarce winter months.

The study revealed interesting patterns in blackbuck dietary preferences, with Atriplex patula (Common Orache) being significantly preferred despite its low availability. Conversely, certain species were avoided, including Prosipus juliflora, likely due to its chemical defenses or low palatability, as well as Solanum nigrum (Black Nightshade) and

Lactuca floridana (Florida Lettuce), which showed significantly lower usage relative to availability. Other species, such as Eruca sativa (Aragula), Rumex obtusifolius (Broad-Leaved Dock), and Malva neglecta (Common Mallow), had neutral usage rates, suggesting they constitute incidental or opportunistic forage. These findings highlight the importance of maintaining key forage species in blackbuck habitats to support their nutritional needs during winter.

### 6. DISCUSSION

#### Vegetation Composition and Structure in Blackbuck Habitats

The vegetation survey across the blackbuck habitats of **Palla Salu, Rathgaon, and Sikandra Rao** revealed a total of 4,125 individuals encompassing a variety of herbaceous and shrubby plant species. The dominance of *Prosopis juliflora* (IVI: 47.81), an invasive species, is particularly noteworthy. This species' high Importance Value Index reflects its aggressive colonization and ability to outcompete native flora, particularly in arid and semi-arid ecosystems (Pasiecznik et al., 2001). Its proliferation poses a potential long-term threat to biodiversity and grassland functionality, particularly by reducing the availability of native forage species essential for herbivores such as the blackbuck (*Antilope cervicapra*).

*Cynodon dactylon* (IVI: 45.84), commonly known as Bermuda grass, was the second most dominant species and is widely known for its palatability and nutritional value to herbivores, including blackbuck (Sankaran, 2005; Isvaran, 2007). Its dominance suggests that the habitat still retains considerable ecological value for foraging ungulates. Similarly, *Poa annua* (IVI: 14.62) and other moderately represented species such as *Calamagrostis epigejos* and *Chenopodium murale* add to the heterogeneity of the grassland, supporting diverse foraging opportunities.

The data reflect a highly uneven plant community structure, with a few dominant species and many rare or infrequent ones. This "log-normal" distribution of plant species is typical in ecosystems subjected to grazing and anthropogenic disturbance (Mishra et al., 2002; Tripathi et al., 2016). While *Cynodon dactylon* supports blackbuck foraging, the unchecked spread of *Prosopis juliflora* may suppress native herbaceous growth, potentially leading to a decline in forage diversity (Rai & Tripathi, 2018). Effective habitat management must therefore balance conservation of key forage species with control of invasive taxa.

#### Blackbuck Dietary Preferences and Foraging Behavior

The **micro-histological analysis** of blackbuck fecal pellets uncovered 860 identifiable fragments belonging to 18 plant species. The prevalence of *Cynodon dactylon* (305 fragments) and *Prosopis juliflora* (151 fragments) aligns with their abundance in the habitat, suggesting a correlation between plant availability and diet (Jhala, 1991; Isvaran, 2004). However, frequency alone is not sufficient to infer preference.

The **Bonferroni-adjusted confidence interval analysis** provides a more nuanced picture of blackbuck dietary selectivity. *Cynodon dactylon* was clearly preferred (34.9% usage vs. 27.7% availability), confirming its role as a core winter forage species. This supports previous findings that blackbuck show selective grazing behavior favoring nutrient-rich C4 grasses during resource-scarce seasons (Jhala, 1997; Sankaran, 2005). Similarly, *Poa annua* and *Rumex obtusifolius* (*Fiddle Dock*) were also disproportionately consumed, possibly due to higher palatability or nutrient content.

Of particular interest is the significant preference for *Chenopodium vulvaria* despite its minimal availability (0.11% in quadrats). This suggests a highly selective foraging behavior potentially aimed at acquiring specific micronutrients or secondary metabolites that play a role in digestion or parasite resistance (Goyal et al., 1988; Robbins, 1993). Selectivity for such species underscores the importance of maintaining microhabitats within the grassland matrix that support rare but nutritionally critical plant species.

Conversely, *Prosopis juliflora* was **avoided** (usage 17.28% vs. availability 25.65%), highlighting the species' limited palatability or possible presence of antinutritional factors (Prasad, 1981; Pasiecznik et al., 2001). This avoidance is ecologically significant — while the plant dominates the landscape, it provides relatively little value to blackbuck, which could lead to habitat degradation and reduced carrying capacity if its spread is left unchecked.

Species such as *Sonchus oleraceus* (*Florida Lettuce*) and *Solanum nigrum* (*Black Nightshade*) were also underutilized, possibly due to chemical defenses (alkaloids, nitrates) that deter herbivory (Robbins, 1993; Kumar & Shahabuddin, 2005). Others like *Aragula* and *Broad-Leaved Dock* fell within the neutral range, consumed in proportion to their availability, reflecting opportunistic rather than preferential foraging.

#### Implications for Conservation and Habitat Management

These findings have direct implications for blackbuck conservation. The selective foraging patterns observed highlight the importance of maintaining populations of high-value forage species, particularly *Cynodon dactylon*, *Poa annua*, and other palatable grasses. In contrast, the dominance and low palatability of *Prosopis juliflora* raise concerns about long-term habitat quality.

To safeguard the blackbuck population and their foraging needs:

- **Invasive species control** (especially *P. juliflora*) should be prioritized to prevent further loss of palatable native flora (Rai & Tripathi, 2018).
- **Grassland restoration** efforts should focus on re-establishing native C4 grasses and herbaceous species that are nutritionally valuable (Rodgers & Panwar, 1988).
- **Seasonal monitoring** of dietary composition should be continued to track changes in forage availability and blackbuck preferences across wet and dry seasons (Singh et al., 2014).
- **Community involvement** in habitat management can reduce overgrazing and encourage sustainable land-use practices near blackbuck habitats (Kumar & Shahabuddin, 2005).

### 7. CONCLUSION

The study conducted across the habitats of **Palla Salu**, **Rathgaon**, and **Sikandra Rao** offers important ecological insights into the intricate relationship between vegetation composition and the foraging behaviour of **blackbuck (*Antilope cervicapra*)**. A critical finding of the research is the **dominance of *Prosopis juliflora*** within these landscapes—an aggressive, non-native species known to alter native plant communities through competition, allelopathy, and habitat modification. Its proliferation poses a substantial threat to the native grassland ecosystem by **displacing indigenous forage species** that are essential to the dietary requirements of herbivores such as blackbuck.

Despite the widespread encroachment of *P. juliflora*, the presence and persistence of **native grasses**, particularly *Cynodon dactylon* and *Poa annua*, reflect the **resilience of certain forage species** under pressure from invasive plant dominance. These grasses continue to serve as critical food sources, especially during the resource-scarce winter season. Additional native herbaceous species such as *Chenopodium vulvaria*, *Chenopodium murale*, *Calamagrostis epigejos*, and *Oxalis montana* were also recorded in notable frequencies within blackbuck diets, indicating **a selective feeding strategy grounded in both availability and palatability**.

The **micro-histological analysis of blackbuck faecal samples** confirmed these dietary patterns, highlighting a strong preference for select species that presumably offer optimal nutritional value. The **low representation of *P. juliflora* in pellet analysis** despite its high abundance in the habitat strongly suggests **active avoidance**, likely due to its **chemical defenses, spiny morphology**, or **low digestibility**. This selective foraging strategy reflects the blackbuck's ability to **maximize nutrient intake** while minimizing the consumption of suboptimal or chemically defended species.

To statistically evaluate foraging preferences, **Bonferroni-adjusted confidence intervals** were applied, enabling a robust categorization of plant species into **preferred, avoided, and neutral groups**. This analytical approach enhances the reliability of dietary preference assessment, distinguishing true selection from random encounter. Species such as *C. dactylon* and *P. annua* consistently fell within the preferred category, reinforcing their importance as **core forage species** during winter.

The findings of this study underscore the **urgent need for adaptive management interventions** that prioritize the **restoration of native forage grasses and control of invasive species** such as *P. juliflora*. The plant community structure observed—marked by **a few dominant species and a long tail of rare or sparsely distributed species**—reflects a pattern typical of degraded ecosystems where ecological balance has been disrupted. Such uneven species distribution not only limits dietary options for herbivores but also affects overall **ecosystem functionality and resilience**.

Restoring native vegetation not only enhances forage availability for blackbuck but also promotes greater **floristic diversity and habitat complexity**, benefiting a broader range of species. By integrating blackbuck dietary preferences with vegetation composition data, **evidence-based conservation strategies** can be formulated. These strategies should aim at sustaining healthy herbivore populations while restoring ecological integrity, ensuring the **long-term conservation of grassland ecosystems and biodiversity in the region**.

### 9. TABLES

#### *Table 9.1 IVI OF PLANTS*

| Scientific Name | No. of Individuals | Frequency | Relative frequency | Density | Relative density | IVI |
|---|---|---|---|---|---|---|
| *Achyranthus aspera* | 51 | 5 | 1.59 | 0.57 | 1.24 | 2.83 |
| *Ageratum houstonianum* | 12 | 2 | 0.64 | 0.13 | 0.29 | 0.93 |
| *Alternanthera sessilis* | 48 | 4 | 1.27 | 0.53 | 1.16 | 2.44 |
| *American germender* | 5 | 1 | 0.32 | 0.06 | 0.12 | 0.44 |
| *Argemone mexicana* | 6 | 2 | 0.64 | 0.07 | 0.15 | 0.78 |
| *Atriplex patula* | 3 | 1 | 0.32 | 0.03 | 0.07 | 0.39 |
| *Calamagrostis epigejos* | 275 | 19 | 6.05 | 3.06 | 6.67 | 12.72 |
| *Calatrichi stagnails* | 4 | 2 | 0.64 | 0.04 | 0.10 | 0.73 |
| *Cannabis sativa* | 6 | 2 | 0.64 | 0.07 | 0.15 | 0.78 |
| *Capparis decidua* | 22 | 5 | 1.59 | 0.24 | 0.53 | 2.13 |
| *Capparis tomentosa* | 6 | 2 | 0.64 | 0.07 | 0.15 | 0.78 |
| *Carduus teniflorous* | 3 | 1 | 0.32 | 0.03 | 0.07 | 0.39 |
| *Chenopodium ficifolium* | 18 | 2 | 0.64 | 0.20 | 0.44 | 1.07 |
| *Chenopodium murale* | 185 | 10 | 3.18 | 2.06 | 4.48 | 7.67 |
| *Chenopodium vulvaria* | 194 | 13 | 4.14 | 2.16 | 4.70 | 8.84 |
| *Coreopsis* | 1 | 1 | 0.32 | 0.01 | 0.02 | 0.34 |
| *Cyanthillium cinerem* | 10 | 1 | 0.32 | 0.11 | 0.24 | 0.56 |
| *Cynodon dactylon* | 754 | 59 | 18.79 | 12.40 | 27.05 | 45.84 |
| *Cynthillium murale* | 3 | 1 | 0.32 | 0.03 | 0.07 | 0.39 |
| *Dhatura innoxia* | 3 | 1 | 0.32 | 0.03 | 0.07 | 0.39 |
| *Eclipta prostata* | 20 | 3 | 0.96 | 0.22 | 0.48 | 1.44 |
| *Eruca sativa* | 40 | 4 | 1.27 | 0.44 | 0.97 | 2.24 |
| *Eryngium aristulatum* | 92 | 5 | 1.59 | 1.02 | 2.23 | 3.82 |
| *Glinus lotoids* | 3 | 1 | 0.32 | 0.03 | 0.07 | 0.39 |
| *Herniaria hirsuta* | 9 | 2 | 0.64 | 0.10 | 0.22 | 0.86 |
| *Ipomea carnea* | 8 | 3 | 0.96 | 0.09 | 0.19 | 1.15 |
| *Lactana flouridicana* | 46 | 4 | 1.27 | 0.51 | 1.12 | 2.39 |
| *Lepidium coronopus* | 16 | 1 | 0.32 | 0.18 | 0.39 | 0.71 |
| *Malwa neglecta* | 29 | 3 | 0.96 | 0.32 | 0.70 | 1.66 |
| *Marsilea minuta* | 17 | 2 | 0.64 | 0.19 | 0.41 | 1.05 |
| *Medicago polymorpha* | 32 | 2 | 0.64 | 0.36 | 0.78 | 1.41 |
| *Oxalis corniculata* | 13 | 2 | 0.64 | 0.14 | 0.32 | 0.95 |
| *Oxalis montana* | 122 | 7 | 2.23 | 1.36 | 2.96 | 5.19 |

| | | | | | | |
|---|---|---|---|---|---|---|
| *Oxalis stricta* | 29 | 1 | 0.32 | 0.33 | 0.73 | 1.05 |
| *Parthenium hysterophora* | 36 | 3 | 0.96 | 0.40 | 0.87 | 1.83 |
| *Poa annua* | 314 | 22 | 7.01 | 3.49 | 7.61 | 14.62 |
| *Polygonum arenastrum* | 28 | 3 | 0.96 | 0.31 | 0.68 | 1.63 |
| *Prosipus juliflora* | 698 | 74 | 23.57 | 11.11 | 24.24 | 47.81 |
| *Rorripa syvestis* | 2 | 2 | 0.64 | 0.02 | 0.05 | 0.69 |
| *Rumex obtusifolius* | 34 | 2 | 0.64 | 0.38 | 0.82 | 1.46 |
| *Rumex pulcher* | 43 | 14 | 4.46 | 0.48 | 1.04 | 5.50 |
| *Sida cordata* | 33 | 3 | 0.96 | 0.37 | 0.80 | 1.76 |
| *Solanum dulcamara* | 62 | 7 | 2.23 | 0.69 | 1.50 | 3.73 |
| *Solanum nigrum* | 40 | 4 | 1.27 | 0.44 | 0.97 | 2.24 |
| *Spergula arvensis* | 10 | 1 | 0.32 | 0.11 | 0.24 | 0.56 |
| *Stellaria media* | 12 | 1 | 0.32 | 0.13 | 0.29 | 0.61 |
| *Sysimbrium irio* | 6 | 1 | 0.32 | 0.07 | 0.15 | 0.46 |
| *Triticum fragiferum* | 52 | 2 | 0.64 | 0.58 | 1.26 | 1.90 |
| Total | 4125 | 314 | 100.00 | 45.83 | 100.00 | 200.00 |

#### *Table 9.2BONFERRONI CI*

| Scientific Name | Fragments | Usage % | Availability % | Lower CI | Upper CI | Preference |
|---|---|---|---|---|---|---|
| *Poa annua* | 0.03 | 0.00 | 0.02 | 0.04 | 8.40 | Preferred |
| *Eruca vesicaria* | 0.01 | 0.00 | 0.00 | 0.01 | 1.10 | Neutral |
| *Cynodon dactylon* | 0.25 | 0.01 | 0.23 | 0.28 | 30.00 | Preferred |
| *Solanum nigrum* | 0.00 | 0.00 | 0.00 | 0.00 | 0.90 | Avoided |
| *Rumex obtusifolius* | 0.00 | 0.00 | 0.00 | 0.01 | 0.86 | Neutral |
| *Medicago polymorpha* | 0.00 | 0.00 | 0.00 | 0.00 | 0.78 | Neutral |
| *Malva sylvestris* | 0.00 | 0.00 | 0.00 | 0.00 | 0.08 | Neutral |
| *Atriplex patula* | 0.00 | 0.00 | 0.00 | 0.01 | 3.28 | Preferred |
| *Oxalis acetosella* | 0.00 | 0.00 | 0.00 | 0.00 | 0.97 | Neutral |
| *Parthenium hysterophorus* | 0.00 | 0.00 | 0.00 | 0.00 | 0.45 | Avoided |
| *Eclipta prostrata* | 0.00 | 0.00 | 0.00 | 0.01 | 1.15 | Neutral |
| *Rumex pulcher* | 0.05 | 0.00 | 0.04 | 0.06 | 26.90 | Preferred |
| *Lactuca floridana* | 0.01 | 0.00 | 0.00 | 0.02 | 4.98 | Avoided |
| *Prosopis juliflora* | 0.00 | 0.00 | 0.00 | 0.01 | 1.29 | Avoided |
| *Chenopodium murale* | 0.01 | 0.00 | 0.01 | 0.02 | 5.22 | Neutral |
| *Alternanthera sessilis* | 0.03 | 0.00 | 0.02 | 0.04 | 7.39 | Neutral |
| *Chenopodium vulvaria* | | | | | 93.75 | Neutral |
| *Calamagrostis epigejos* | 0.03 | 0.00 | 0.02 | 0.04 | 8.40 | Neutral |

#### *Table 9.1 IVI OF PLANTS*

| Scientific Name | No. of Individuals | Frequency | Relative frequency | Density | Relative density | IVI |
|---|---|---|---|---|---|---|
| *Achyranthus aspera* | 51 | 5 | 1.59 | 0.57 | 1.24 | 2.83 |
| *Ageratum houstonianum* | 12 | 2 | 0.64 | 0.13 | 0.29 | 0.93 |
| *Alternanthera sessilis* | 48 | 4 | 1.27 | 0.53 | 1.16 | 2.44 |
| *American germender* | 5 | 1 | 0.32 | 0.06 | 0.12 | 0.44 |
| *Argemone mexicana* | 6 | 2 | 0.64 | 0.07 | 0.15 | 0.78 |
| *Atriplex patula* | 3 | 1 | 0.32 | 0.03 | 0.07 | 0.39 |
| *Calamagrostis epigejos* | 275 | 19 | 6.05 | 3.06 | 6.67 | 12.72 |
| *Calatrichi stagnails* | 4 | 2 | 0.64 | 0.04 | 0.10 | 0.73 |
| *Cannabis sativa* | 6 | 2 | 0.64 | 0.07 | 0.15 | 0.78 |
| *Capparis decidua* | 22 | 5 | 1.59 | 0.24 | 0.53 | 2.13 |
| *Capparis tomentosa* | 6 | 2 | 0.64 | 0.07 | 0.15 | 0.78 |
| *Carduus teniflorous* | 3 | 1 | 0.32 | 0.03 | 0.07 | 0.39 |
| *Chenopodium ficifolium* | 18 | 2 | 0.64 | 0.20 | 0.44 | 1.07 |

| | | | | | | |
|---|---|---|---|---|---|---|
| *Chenopodium murale* | 185 | 10 | 3.18 | 2.06 | 4.48 | 7.67 |
| *Chenopodium vulvaria* | 194 | 13 | 4.14 | 2.16 | 4.70 | 8.84 |
| *Coreopsis* | 1 | 1 | 0.32 | 0.01 | 0.02 | 0.34 |
| *Cyanthillium cinerem* | 10 | 1 | 0.32 | 0.11 | 0.24 | 0.56 |
| *Cynodon dactylon* | 754 | 59 | 18.79 | 12.40 | 27.05 | 45.84 |
| *Cynthillium murale* | 3 | 1 | 0.32 | 0.03 | 0.07 | 0.39 |
| *Dhatura innoxia* | 3 | 1 | 0.32 | 0.03 | 0.07 | 0.39 |
| *Eclipta prostata* | 20 | 3 | 0.96 | 0.22 | 0.48 | 1.44 |
| *Eruca sativa* | 40 | 4 | 1.27 | 0.44 | 0.97 | 2.24 |
| *Eryngium aristulatum* | 92 | 5 | 1.59 | 1.02 | 2.23 | 3.82 |
| *Glinus lotoids* | 3 | 1 | 0.32 | 0.03 | 0.07 | 0.39 |
| *Herniaria hirsuta* | 9 | 2 | 0.64 | 0.10 | 0.22 | 0.86 |
| *Ipomea carnea* | 8 | 3 | 0.96 | 0.09 | 0.19 | 1.15 |
| *Lactana flouridicana* | 46 | 4 | 1.27 | 0.51 | 1.12 | 2.39 |
| *Lepidium coronopus* | 16 | 1 | 0.32 | 0.18 | 0.39 | 0.71 |
| *Malwa neglecta* | 29 | 3 | 0.96 | 0.32 | 0.70 | 1.66 |
| *Marsilea minuta* | 17 | 2 | 0.64 | 0.19 | 0.41 | 1.05 |
| *Medicago polymorpha* | 32 | 2 | 0.64 | 0.36 | 0.78 | 1.41 |
| *Oxalis corniculata* | 13 | 2 | 0.64 | 0.14 | 0.32 | 0.95 |
| *Oxalis montana* | 122 | 7 | 2.23 | 1.36 | 2.96 | 5.19 |
| *Oxalis stricta* | 29 | 1 | 0.32 | 0.33 | 0.73 | 1.05 |
| *Parthenium hysterophora* | 36 | 3 | 0.96 | 0.40 | 0.87 | 1.83 |
| *Poa annua* | 314 | 22 | 7.01 | 3.49 | 7.61 | 14.62 |
| *Polygonum arenastrum* | 28 | 3 | 0.96 | 0.31 | 0.68 | 1.63 |
| *Prosipus juliflora* | 698 | 74 | 23.57 | 11.11 | 24.24 | 47.81 |
| *Rorripa syvestis* | 2 | 2 | 0.64 | 0.02 | 0.05 | 0.69 |
| *Rumex obtusifolius* | 34 | 2 | 0.64 | 0.38 | 0.82 | 1.46 |
| *Rumex pulcher* | 43 | 14 | 4.46 | 0.48 | 1.04 | 5.50 |
| *Sida cordata* | 33 | 3 | 0.96 | 0.37 | 0.80 | 1.76 |
| *Solanum dulcamara* | 62 | 7 | 2.23 | 0.69 | 1.50 | 3.73 |
| *Solanum nigrum* | 40 | 4 | 1.27 | 0.44 | 0.97 | 2.24 |
| *Spergula arvensis* | 10 | 1 | 0.32 | 0.11 | 0.24 | 0.56 |
| *Stellaria media* | 12 | 1 | 0.32 | 0.13 | 0.29 | 0.61 |
| *Sysimbrium irio* | 6 | 1 | 0.32 | 0.07 | 0.15 | 0.46 |
| *Triticum fragiferum* | 52 | 2 | 0.64 | 0.58 | 1.26 | 1.90 |
| Total | 4125 | 314 | 100.00 | 45.83 | 100.00 | 200.00 |

### 10.PICTURES OF MICRO-HISTOLOGICAL SLIDES WITH PLANTS

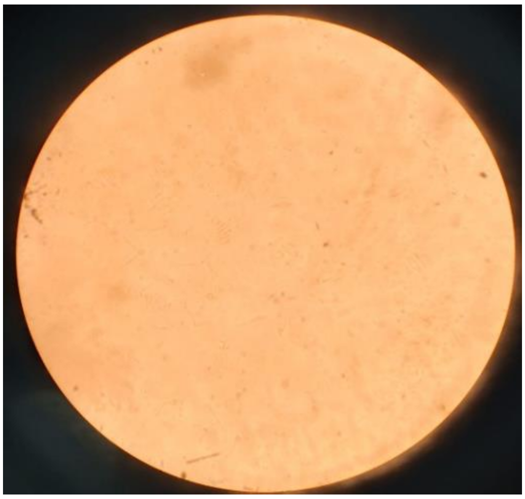

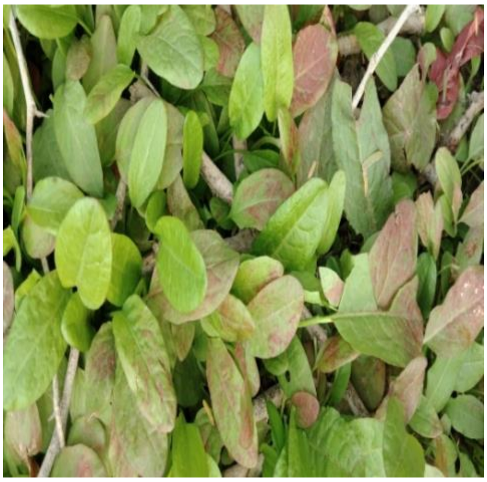

***Picture 1 Rumex pulcher***

***Rumex pulcher* (Fiddle Dock)**

**Stomatal Type**: **Anomocytic**.

**Distribution**: Primarily **hypostomatic**.

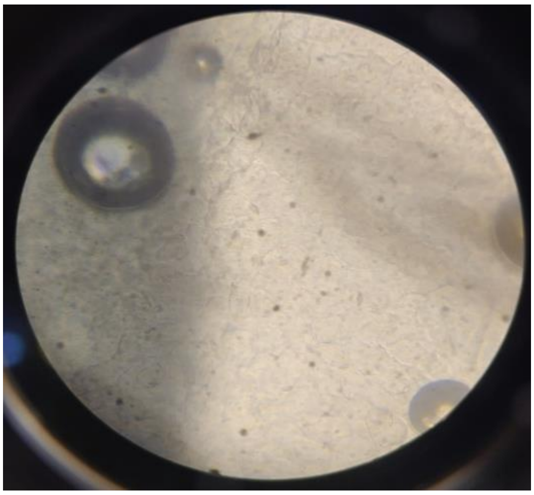

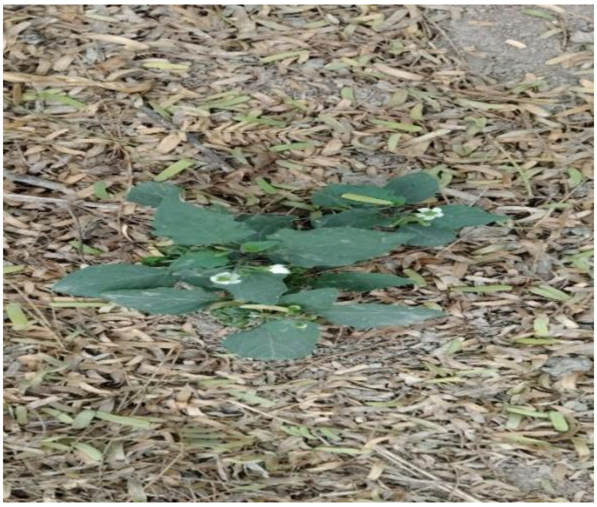

**Picture 2 Solanum nigrum**

***Solanum nigrum* (Black Nightshade)**

- **Stomatal Type**: **Anisocytic** (surrounded by three unequal subsidiary cells).
- **Distribution**: **Amphistomatic**.
- **Additional Features**: Presence of glandular trichomes .

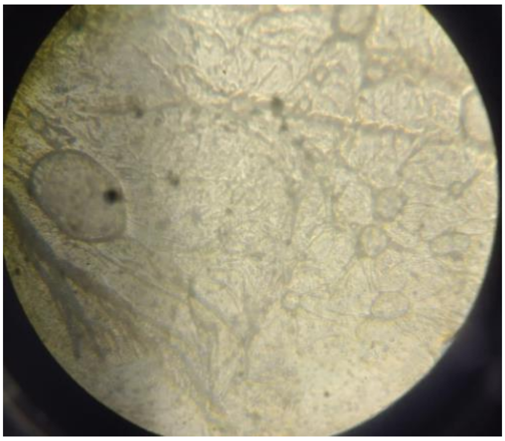

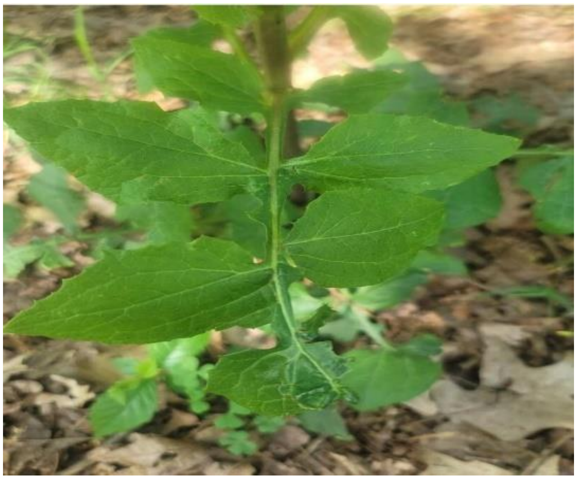

***Pictures 3 Lactana flouridicana***

***Lactuca floridana*** **(Woodland Lettuce)**

**Stomatal Type: Anomocytic.**

**Distribution: Amphistomatic**

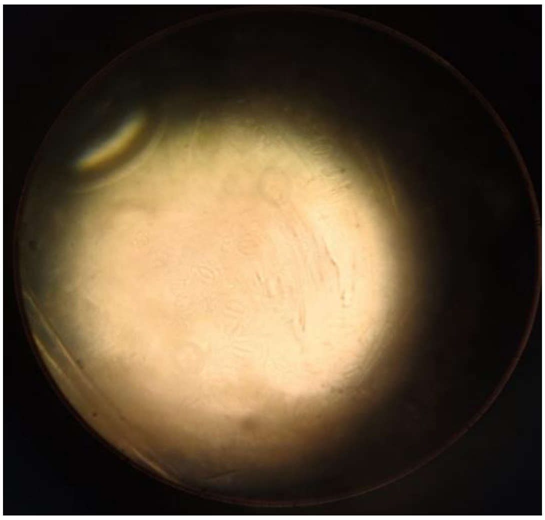

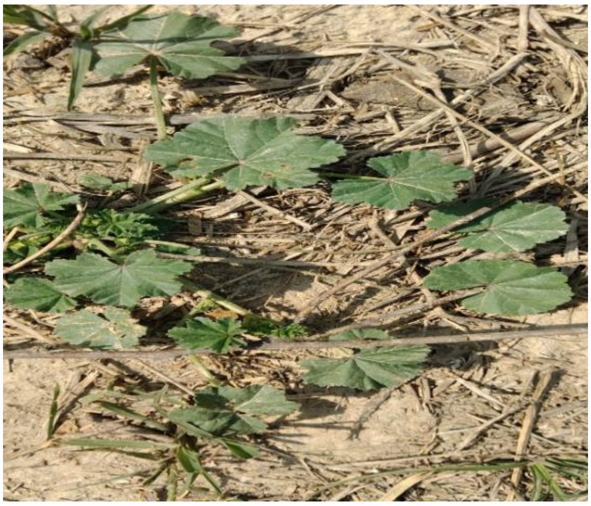

***Pictures 4 Malwa sylvestris***

***Malva sylvestris*** **(Common Mallow)**

**Stomatal Type: Anisocytic.**

**Distribution: Amphistomatic**

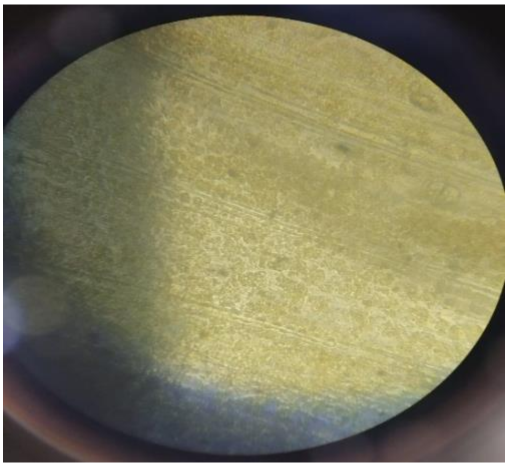

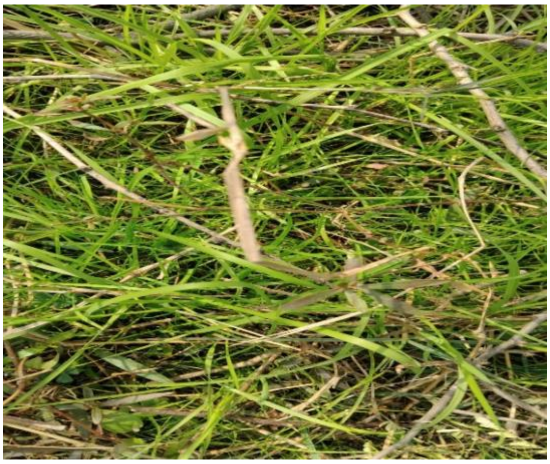

**Pictures 5 Cynodon dactylon**

#### *Cynodon dactylon* (Bermuda Grass)

**Stomatal Type**: **Paracytic** (each stoma accompanied by two subsidiary cells parallel to the pore).

**Distribution**: **Amphistomatic**.

**Additional Features**: Epidermal cells are sinuous; silica bodies present.

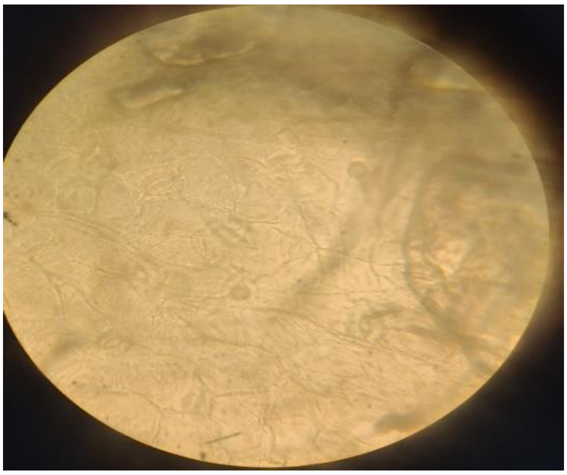

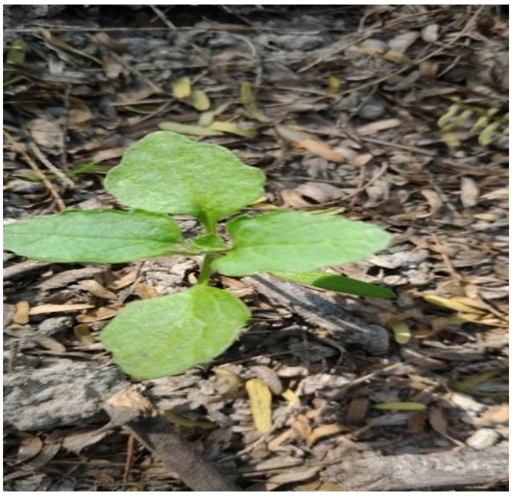

**Pictures 6 Chenopodium vulvaria**

#### *Chenopodium vulvaria* (Stinking Goosefoot)

**Stomatal Type**: **Anisocytic**.

**Distribution**: **Amphistomatic**.

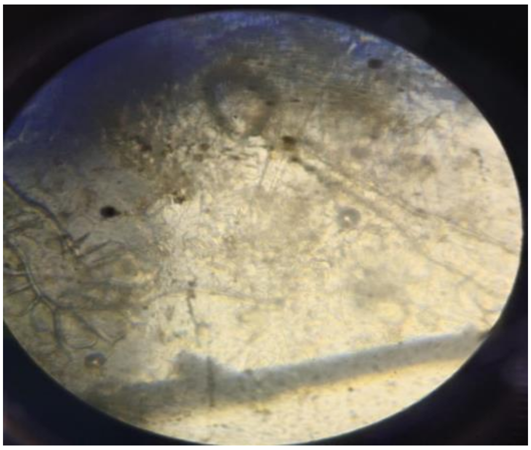
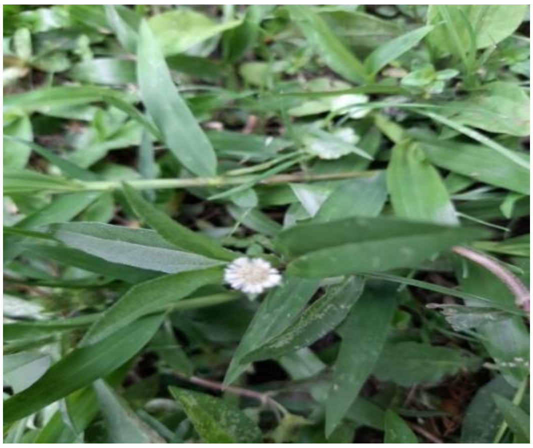

*Picture 7 Eclipta prostrata*

#### *Eclipta prostrata* (False Daisy)

**Stomatal Type**: **Anomocytic**.

**Distribution**: **Amphistomatic**.

**Additional Features**: Presence of capitate glandular trichomes

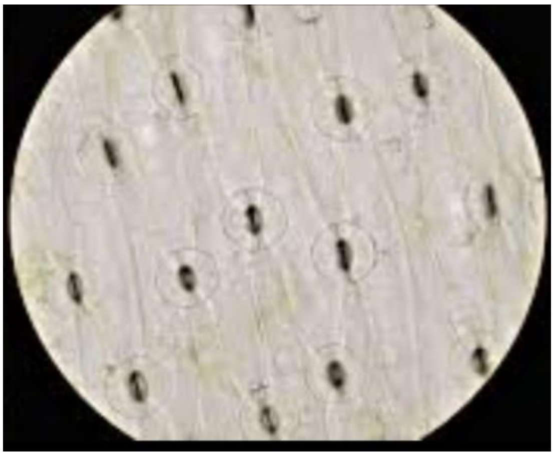
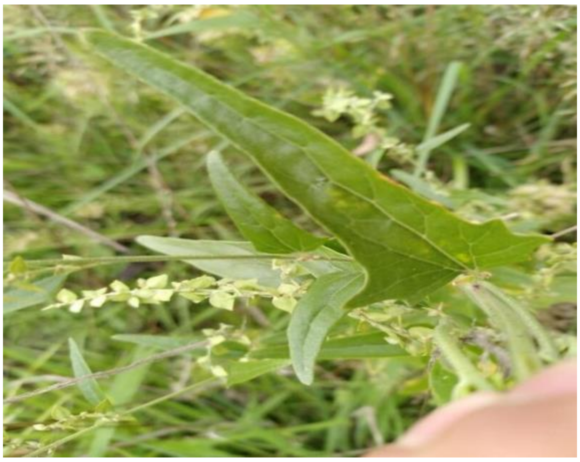

*Picture 8 Atriplex patula*

#### *Atriplex patula* (Common Orache)

**Stomatal Type**: **Anomocytic**.

**Distribution**: **Amphistomatic**

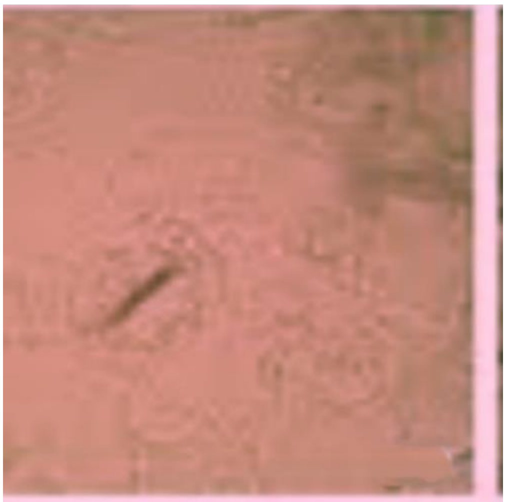
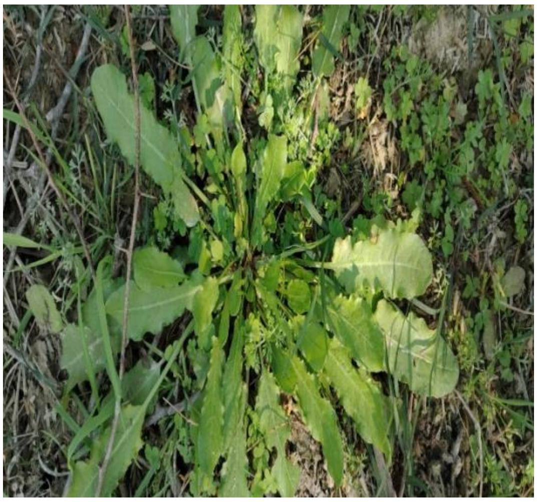

***Picture 9 Rumex obtusifolius***

#### *Rumex obtusifolius* (Broad-leaved Dock)

- ***S*tomatal Type**: **Anomocytic**.
- **Distribution**: Primarily **hypostomatic**

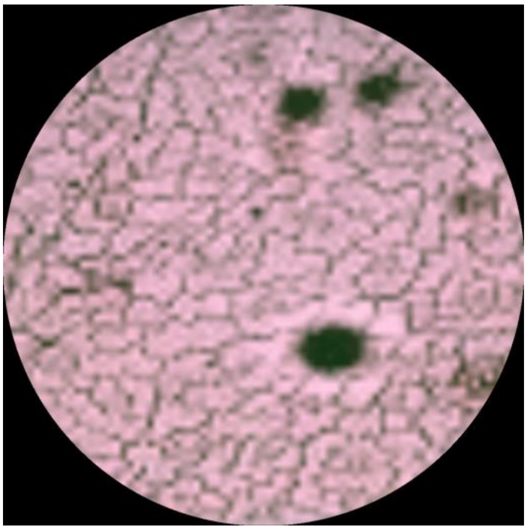
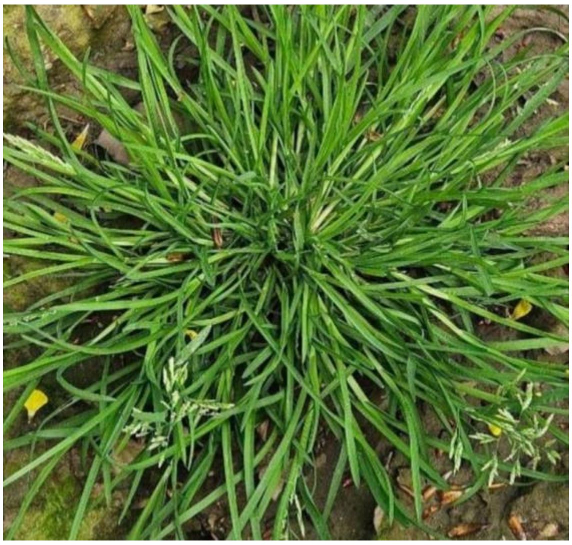

***Picture 10 Poa annua***

#### *Poa annua* (Annual Bluegrass)

**Stomatal Type**: Typically **dumbbell-shaped** guard cells, common in grasses.

**Distribution**: **Amphistomatic** (stomata on both leaf surfaces).

**Additional Features**: Stomata are arranged in parallel rows between veins.

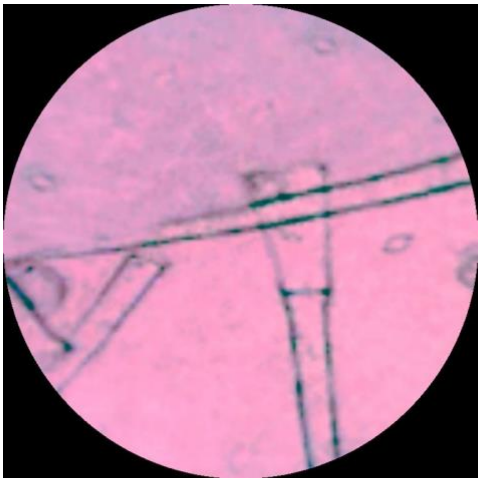

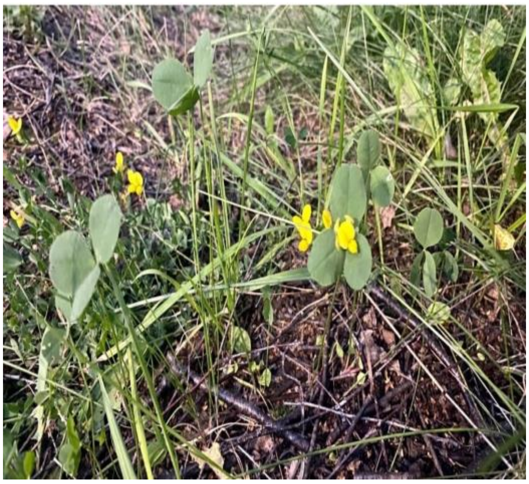

*Picture 11 Medicago polymorpha*

#### Medicago polymorpha (Burr Medic)

**Stomatal Type**: **Paracytic**.

**Distribution**: **Amphistomatic**

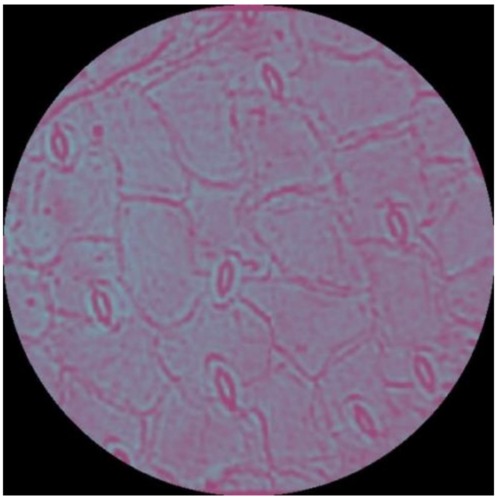

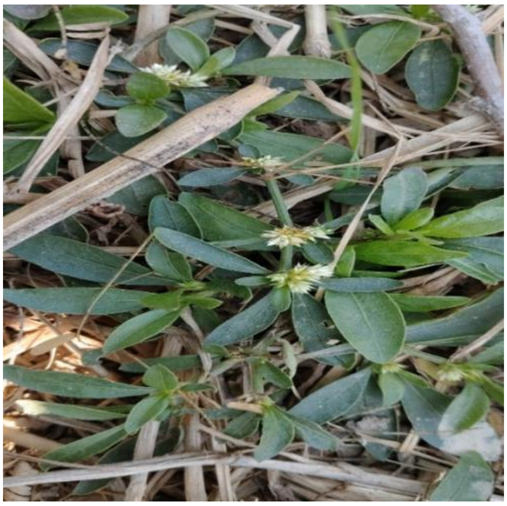

**Pictre 12 Alternanthera sessillis**

#### *Alternanthera sessilis* (Sessile Joyweed)

**Stomatal Type**: **Paracytic**.

**Distribution**: **Amphistomatic**

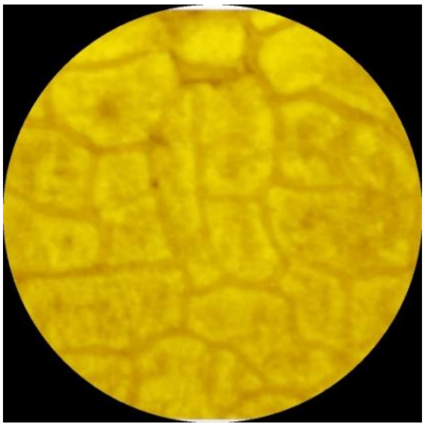

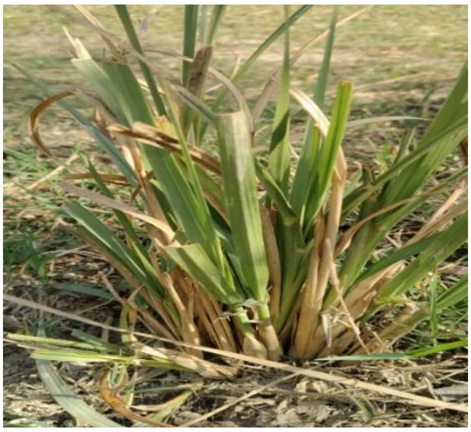

*Picture 13 Calamagrostis epigejos*

#### *Calamagrostis epigejos* (Wood Small-reed)

**Stomatal Type**: **Dumbbell-shaped** guard cells, typical of grasses.

**Distribution**: **Amphistomatic**.

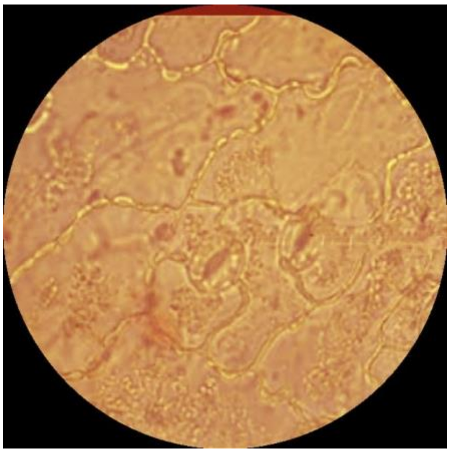

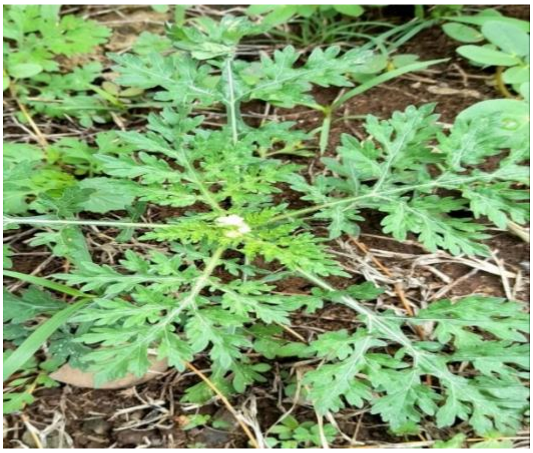

*Picture 14 Parthenium hysteriophorous*

#### *Parthenium hysterophorus* (Santa Maria Feverfew)

**Stomatal Type**: **Tetracytic** (four subsidiary cells, one on each side and two at the ends).

**Distribution**: **Amphistomatic** .

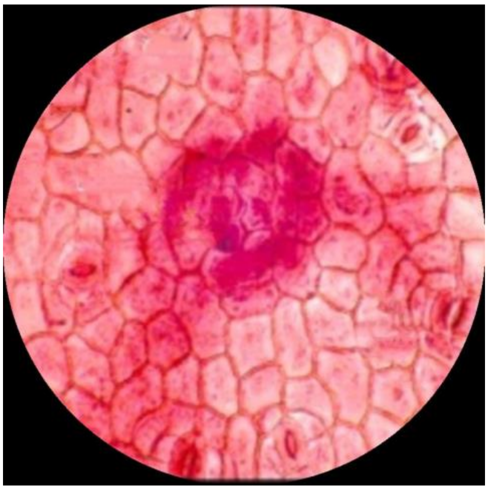

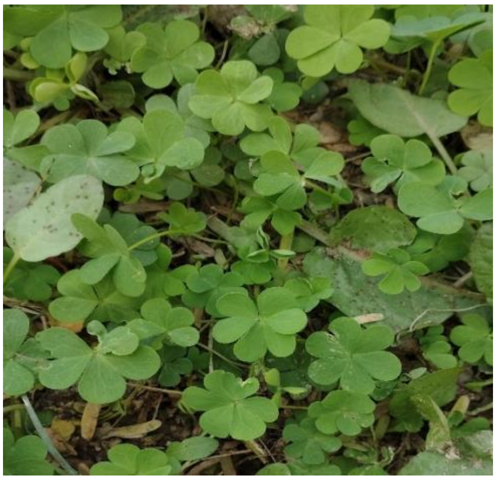

*Picture 15 Oxalis acetosella*

#### Oxalis acetosella (Wood Sorrel)

**Stomatal Type**: **Anomocytic**.

**Distribution**: Primarily **hypostomatic**.

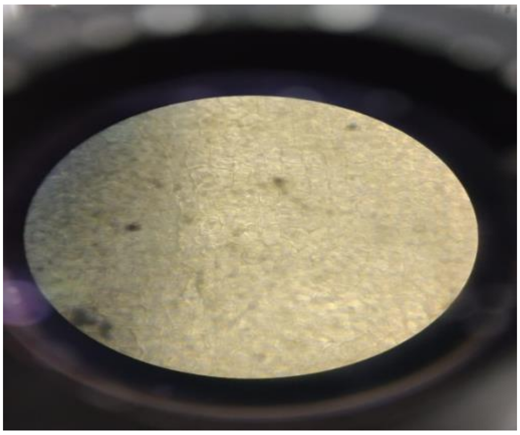

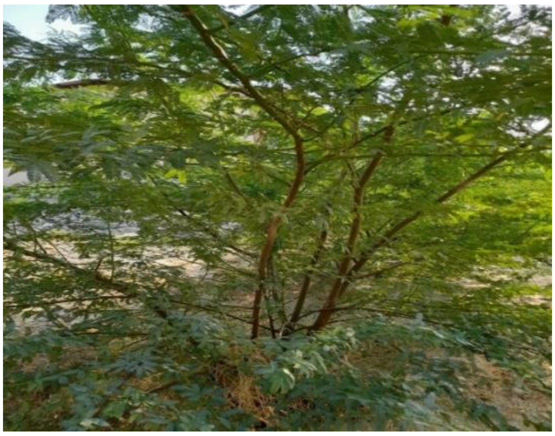

*Picture 16 Prosipus juliflora*

#### *Prosopis juliflora* (Mesquite)

**Stomatal Type**: **Anomocytic**.

**Distribution**: **Amphistomatic**

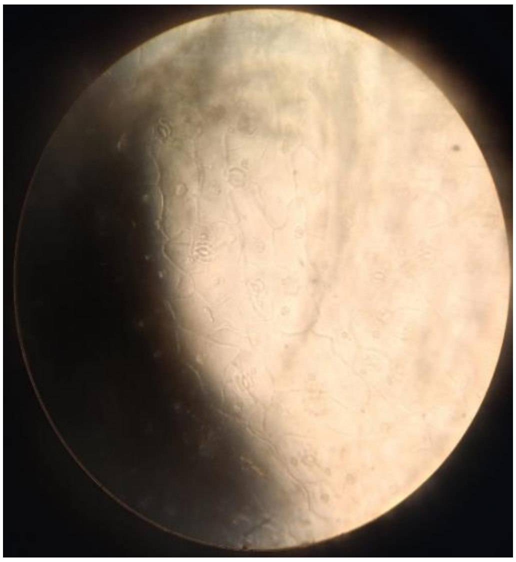

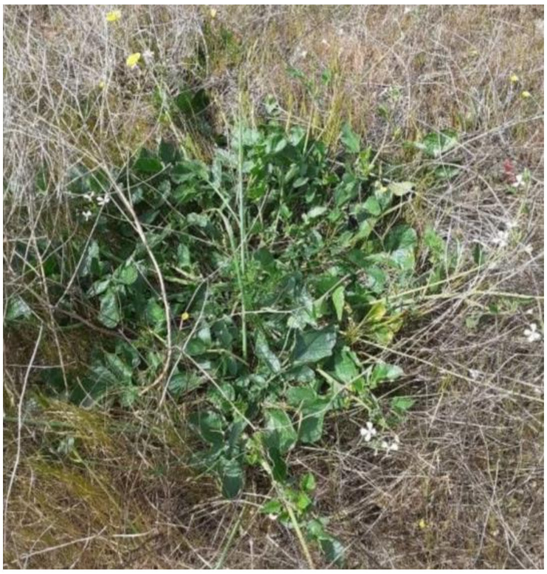

**Picture** ***17 Eruca vesicaria***

#### Eruca vesicaria (Arugula)

**Stomatal Type**: **Anomocytic** (surrounded by cells similar to other epidermal cells).

**Distribution**: Primarily **hypostomatic** (stomata mainly on the lower surface.

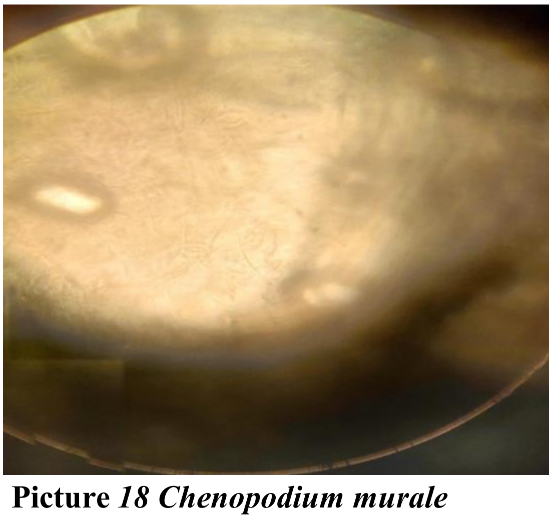

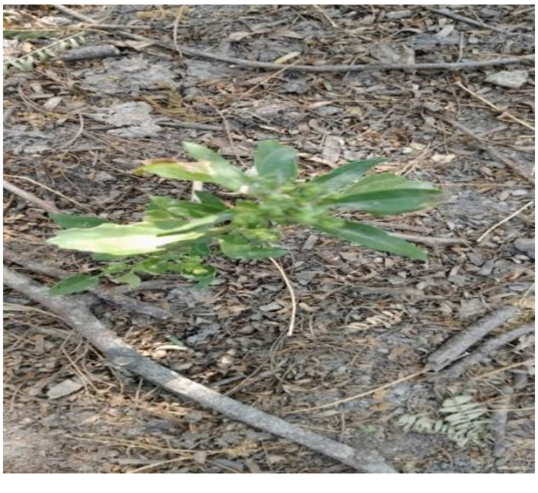

**Picture** ***18 Chenopodium murale***

#### *Chenopodium murale* (Nettle-leaved Goosefoot)

**Stomatal Type**: **Anisocytic**.

**Distribution**: **Amphistomatic** .

### 11.PICTURES

#### PALL SALU

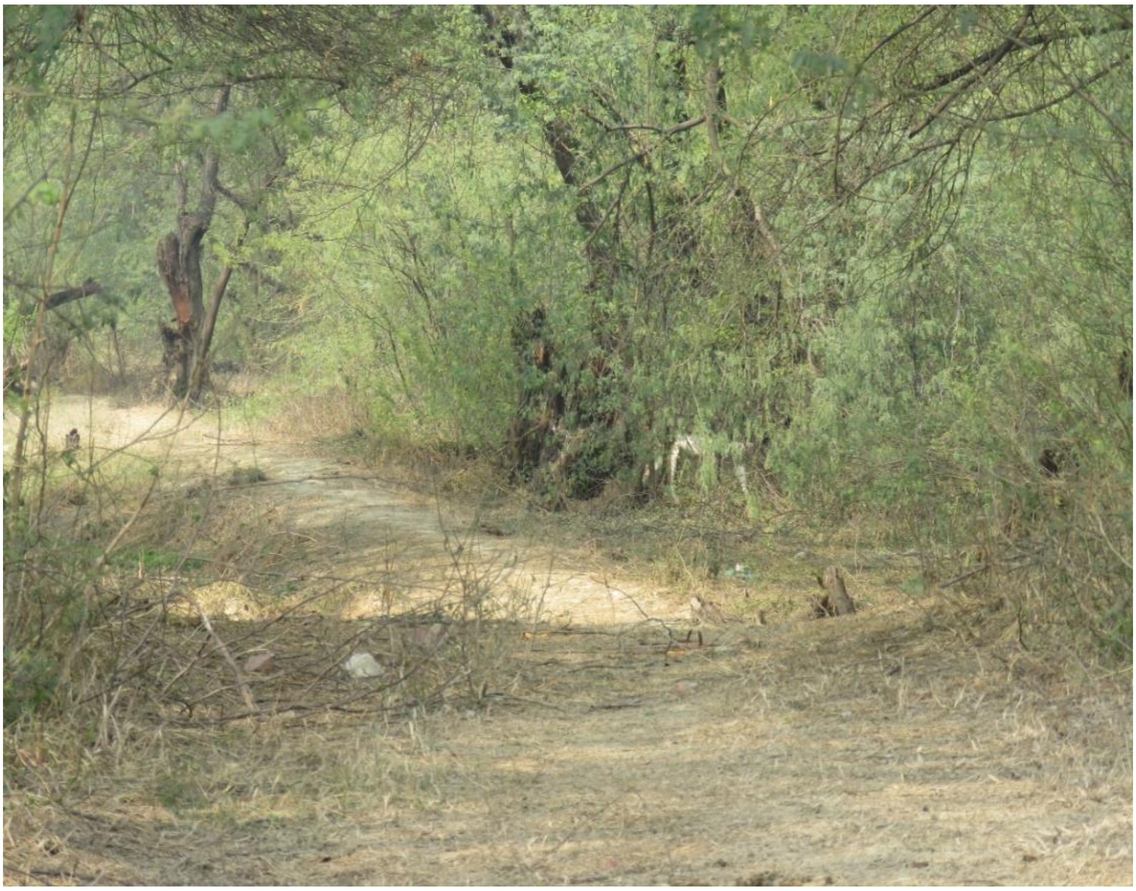

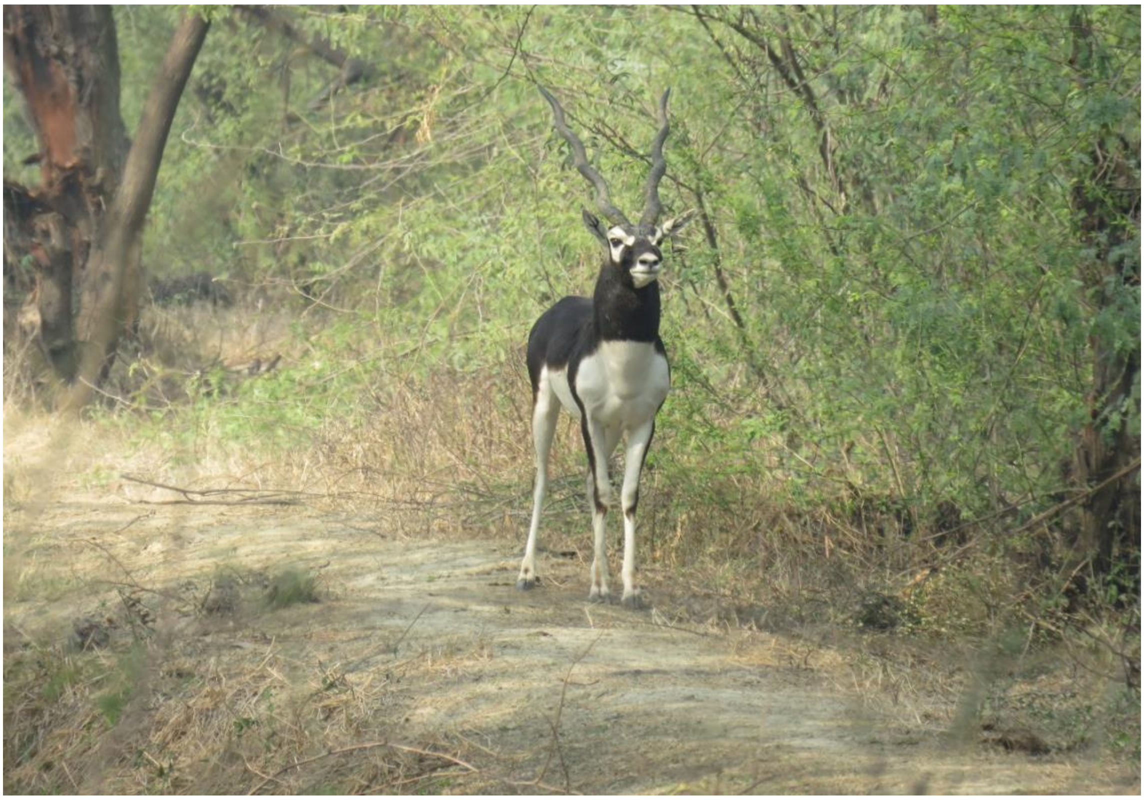

### RATHGAON

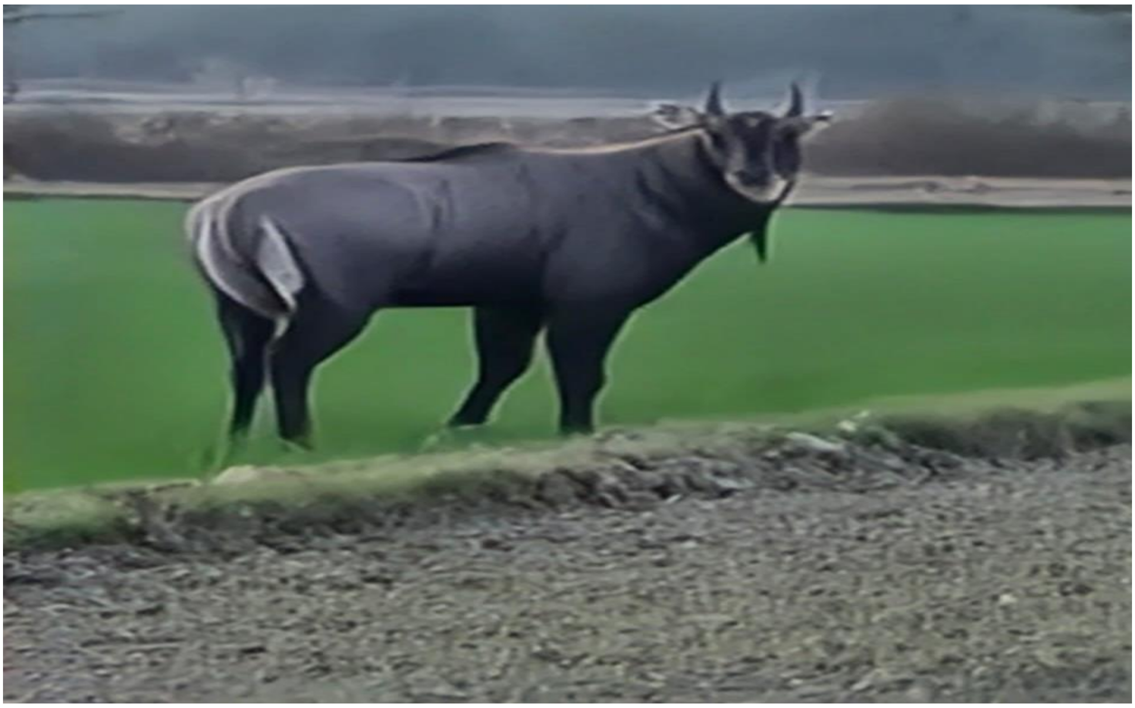

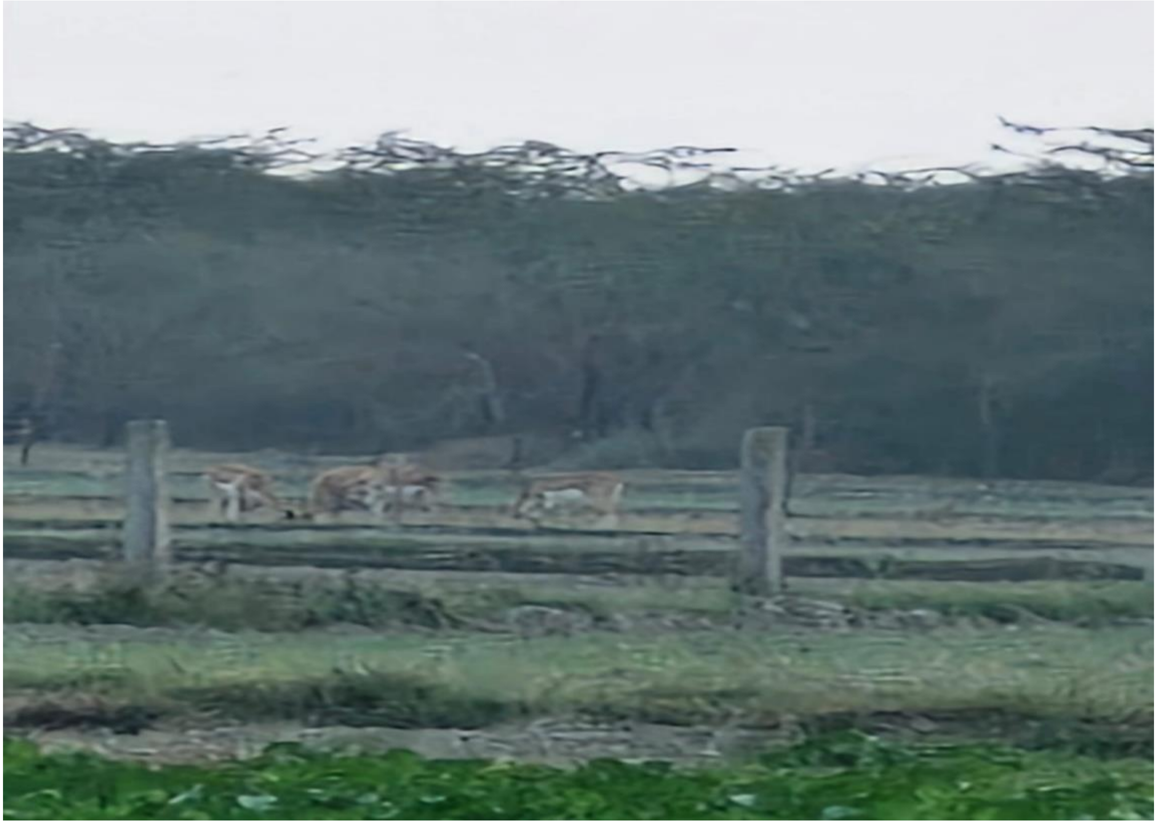

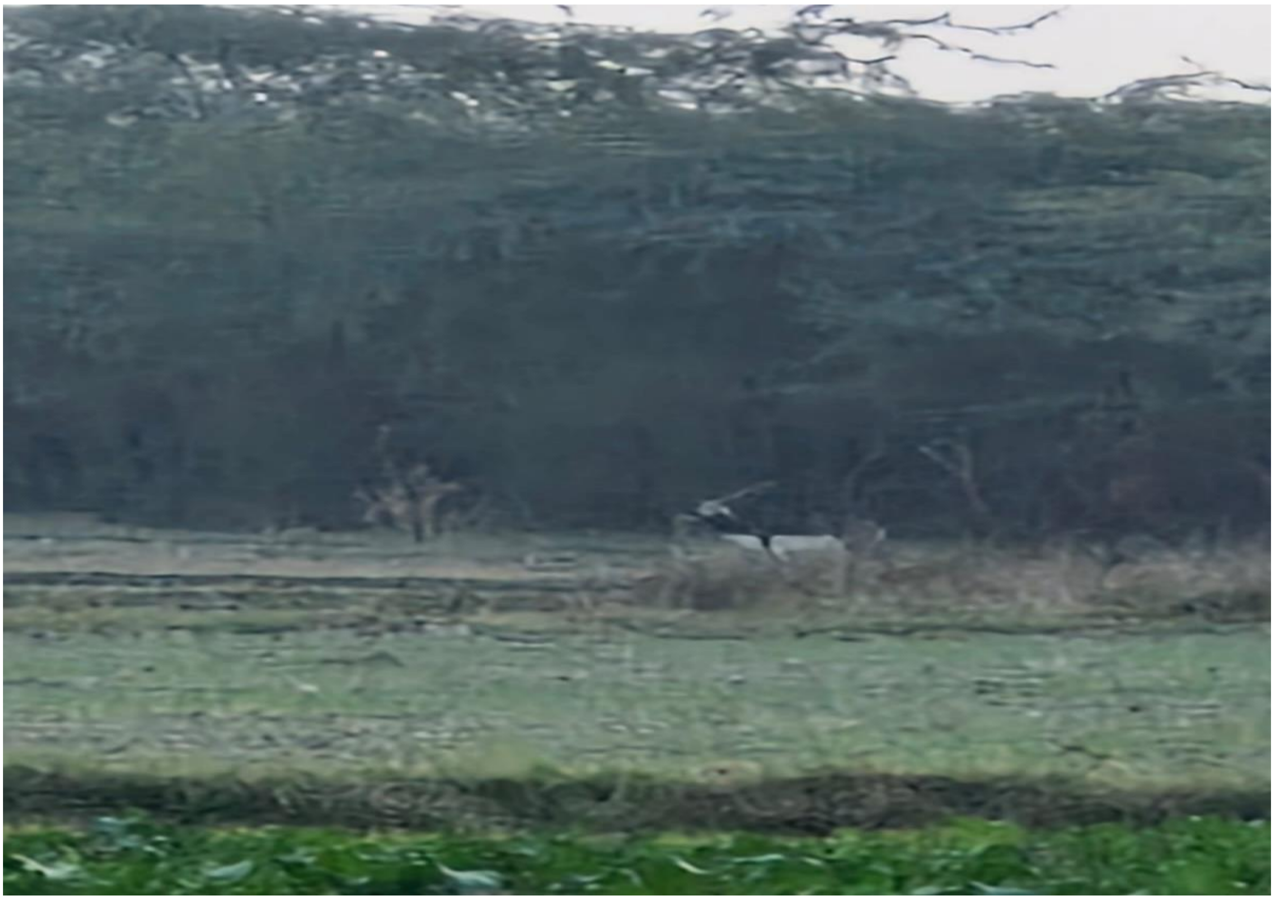

### SIKANDRA RAO

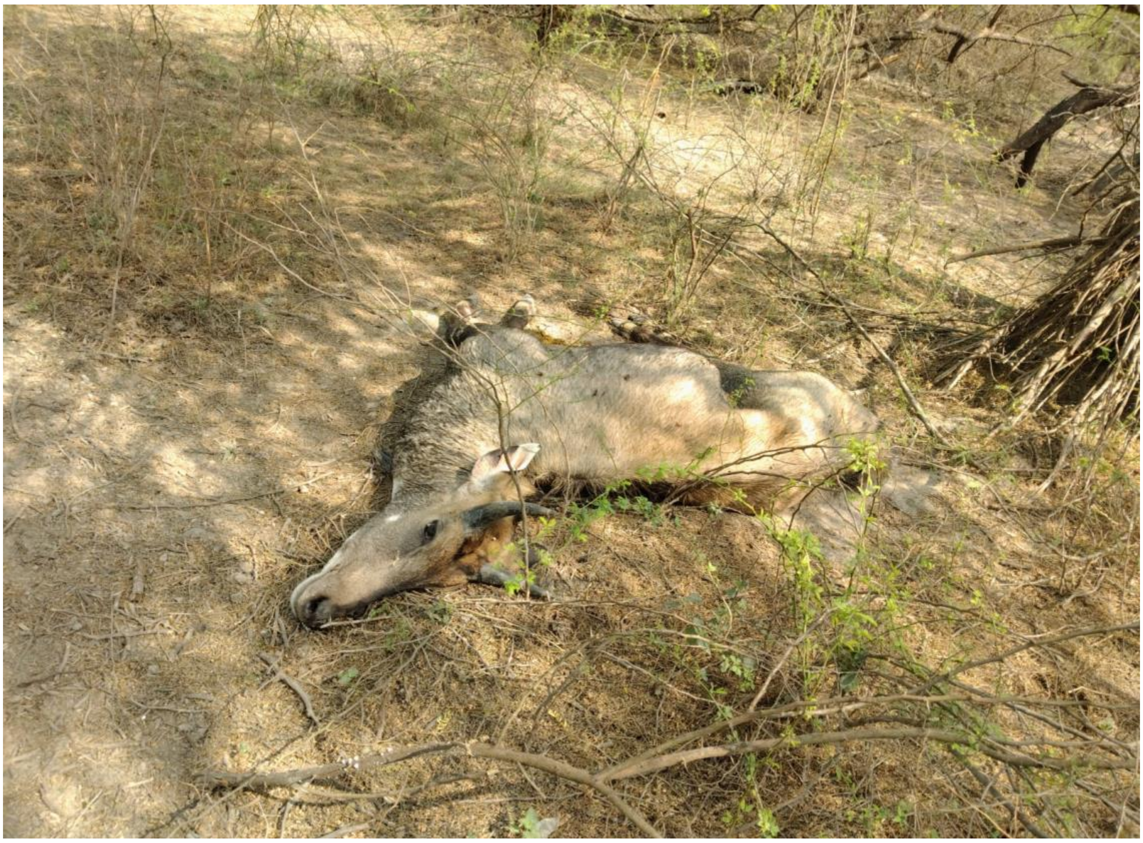

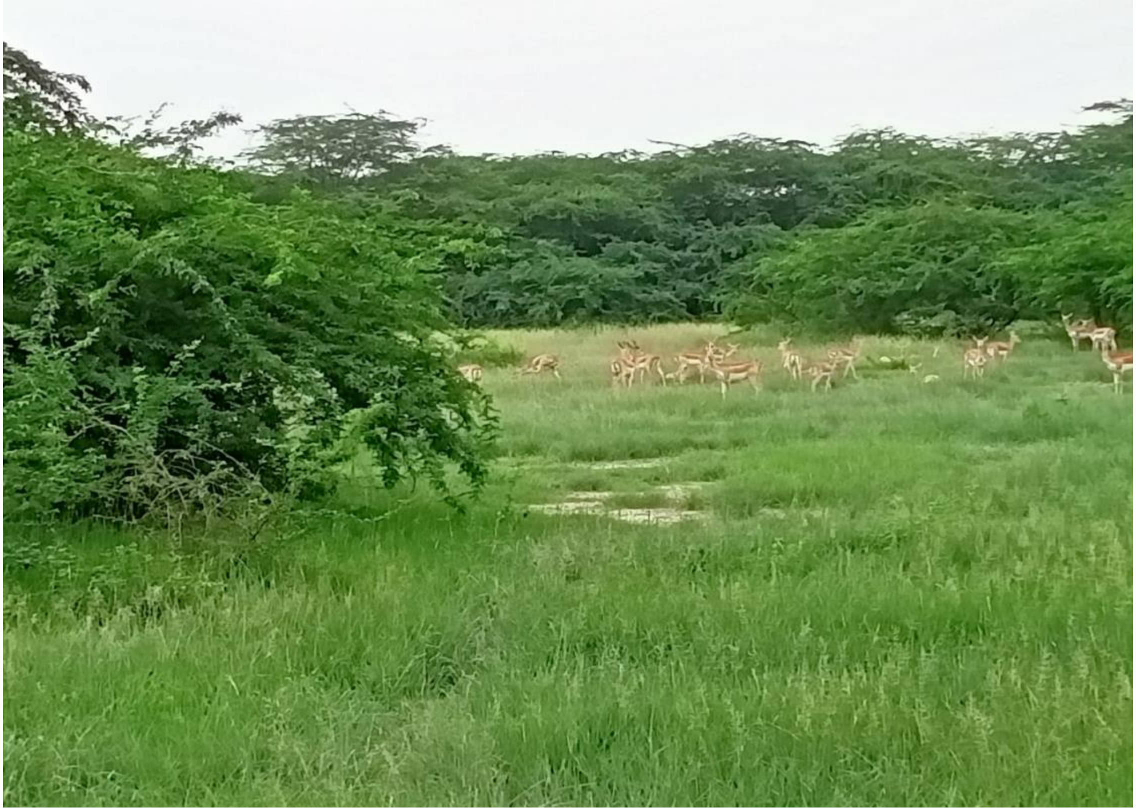

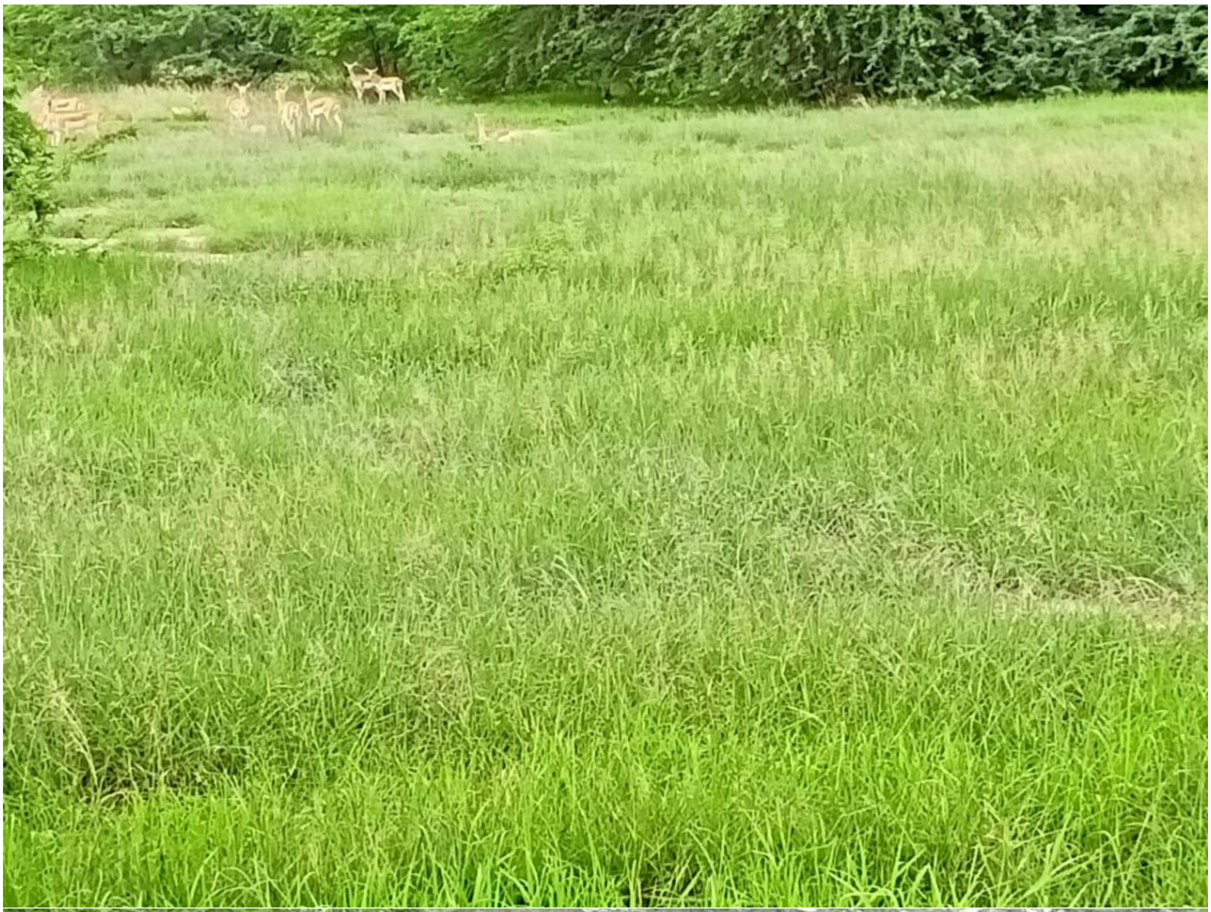

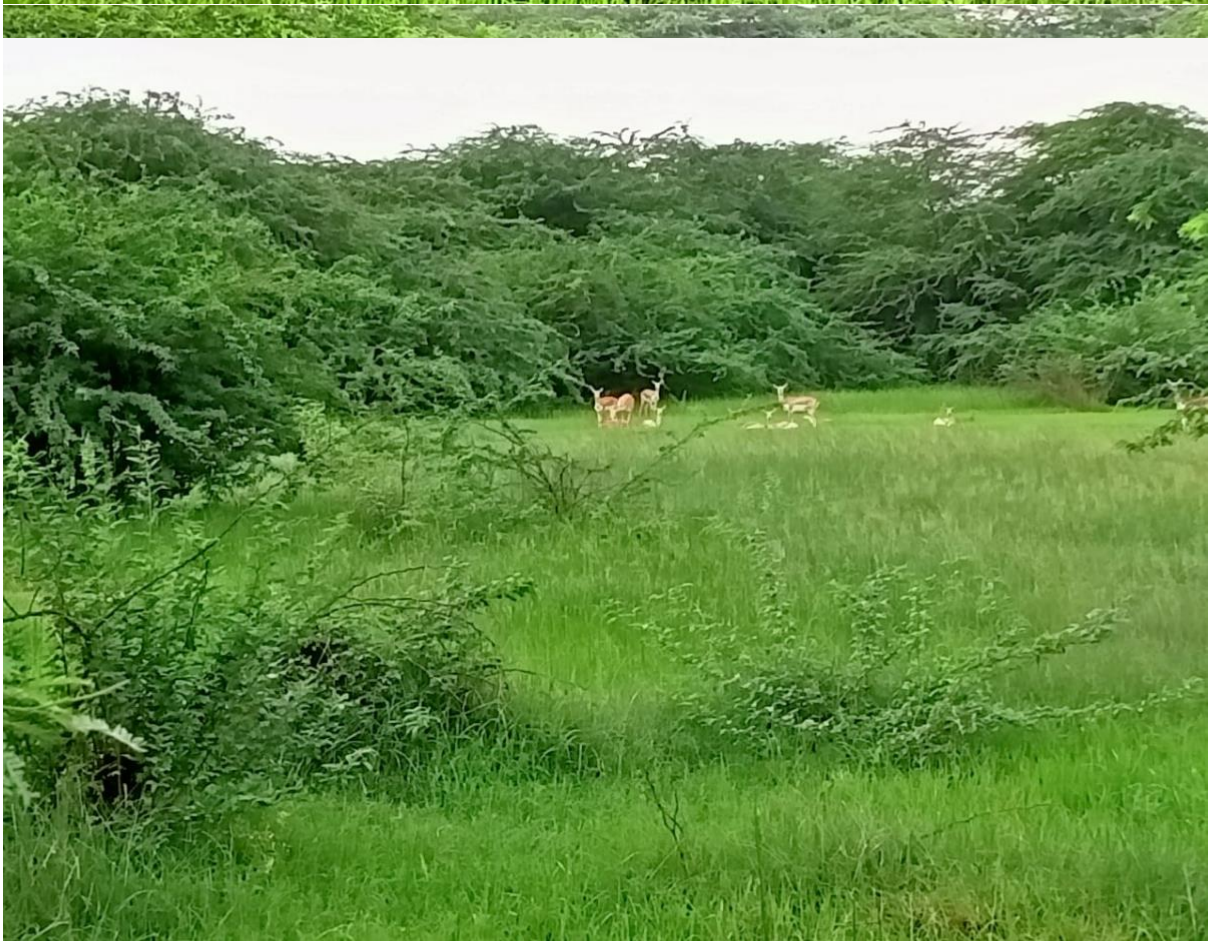

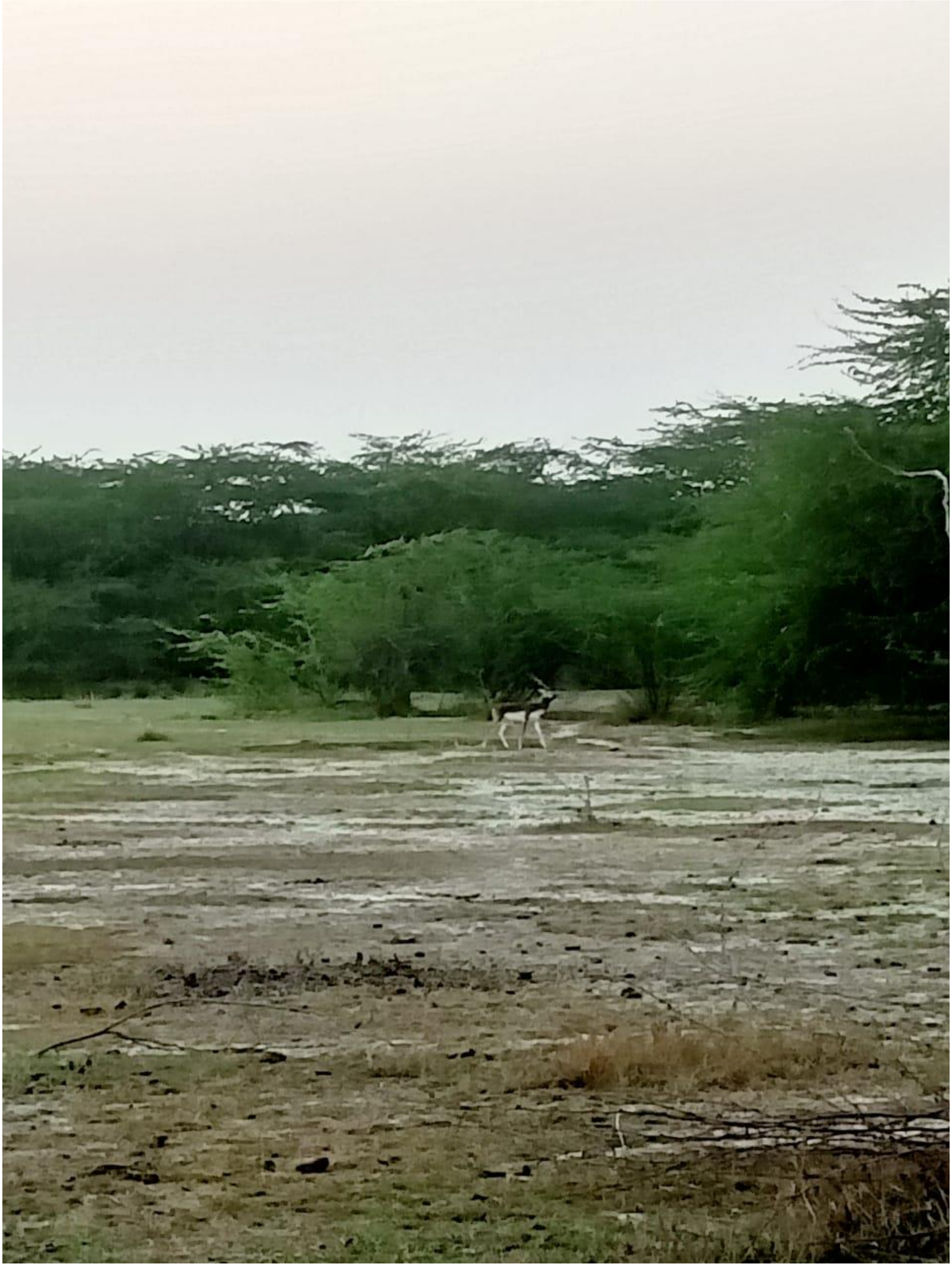

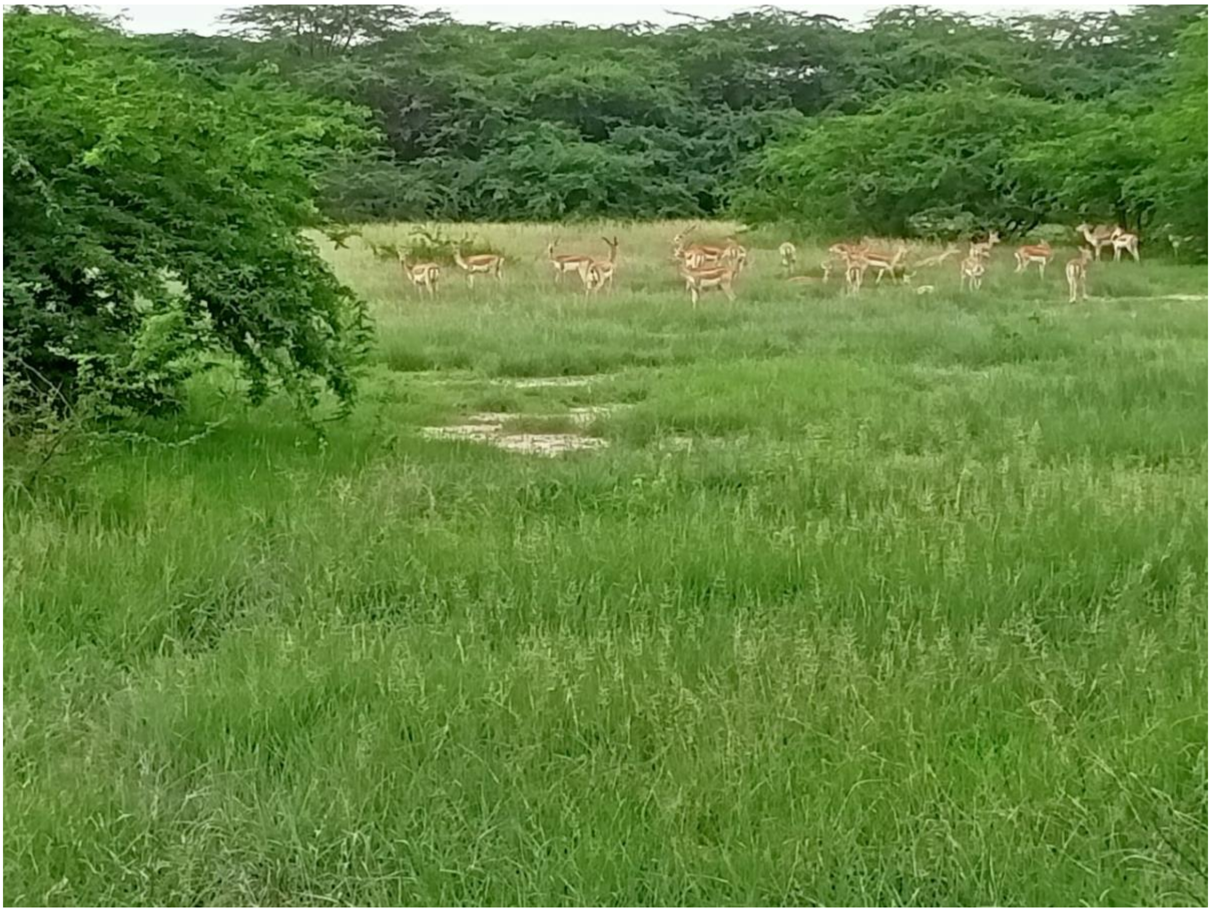

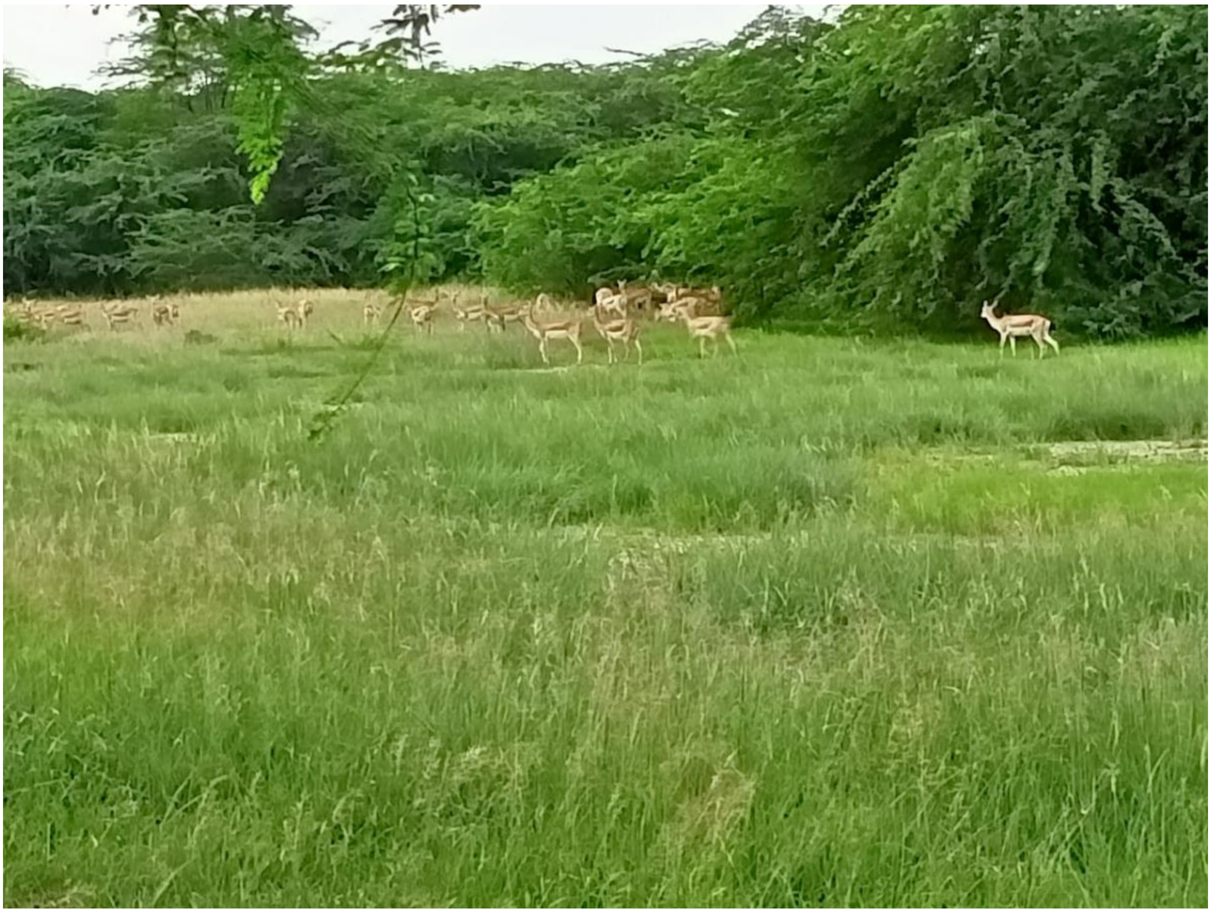
